# Structural basis for repeat-associated non-AUG translation initiation on C9orf72 mRNA

**DOI:** 10.64898/2026.10.25.761709

**Authors:** Ross Kaufhold, Wenzhao Dong, Ishita Purwar, Brenna Saladin, Clare M. Wieland, Amy Krans, Erinn Laimon, Indranil Malik, Suada Leskaj, Tal Havilio, Masaaki Sokabe, Christopher S. Fraser, Peter K. Todd, Jailson Brito Querido

**Affiliations:** Department of Biological Chemistry, University of Michigan, Ann Arbor, MI, USA; Life Sciences Institute, University of Michigan, Ann Arbor, MI, USA; Center for RNA Biomedicine, University of Michigan, Ann Arbor, MI, USA; Department of Neurology, University of Michigan, Ann Arbor, MI, USA; Department of Biotechnology, Indian Institute of Technology Hyderabad, Telangana, India; Department of Molecular and Cellular Biology, College of Biological Sciences, University of California, Davis, CA, USA

## Abstract

A GGGGCC (G_4_C_2_) hexanucleotide repeat expansion in the first intron of *C9orf72* is the most common monogenic cause of Amyotrophic Lateral Sclerosis (ALS) and Frontotemporal Dementia (FTD). This element serves as a template for repeat-associated non-AUG (RAN) translation, which generates neurotoxic dipeptide repeat (DPR) proteins that aggregate in patient brains and contribute to neurodegeneration in model systems. To understand the mechanisms underlying RAN translation initiation, we used cryo-electron microscopy to determine the structure of an *in vitro* assembled human late-stage C9orf72 translation initiation complex at 3.2 Å resolution and a near-native complex purified from human cell lysate at 3.5 Å resolution. RAN translation on C9orf72 mRNA initiates predominantly at a near-cognate CUG codon that is located 24 nucleotides upstream of the G_4_C_2_ repeat and is flanked by a strong endogenous Kozak sequence. The structures further reveal a direct interaction between the G_4_C_2_ repeat and 18S ribosomal RNA expansion segment 9 (ES9S) on the 40S small ribosomal subunit through a kissing-loop interaction. Preventing this RNA-RNA interaction using complementary antisense oligonucleotides significantly reduces RAN translation *in vitro* and in patient-derived iNeurons, indicating that this structural interaction is functionally important. These findings provide mechanistic insights into RAN translation initiation and highlight a novel potential therapeutic approach to mitigate toxic DPR production in C9orf72-associated neurodegenerative disease.

---

Translation of mRNA into protein is a complex process comprising four main steps: initiation, elongation, termination, and ribosome recycling. While cells can regulate each of these steps, most translation regulation occurs during initiation^1^. Canonical translation initiation involves mRNA binding to the ribosomal complex, followed by positioning of the start codon in the decoding center of the 40S small ribosomal subunit. This process requires binding of eukaryotic initiation factors (eIFs) and a ternary complex (TC) consisting of eIF2-GTP-tRNA_i_^Met^ to the 40S small subunit to form the 43S pre-initiation complex (43S)^1,2^. In parallel, the cap-binding complex eIF4F (composed of the cap-binding protein eIF4E, RNA helicase eIF4A, and scaffolding protein eIF4G) binds to the m^7^G cap at the 5′ end of the mRNA and interacts with poly(A)-binding protein (PABP) attached at the 3^*1*^ untranslated region (UTR) of mRNA. The eIF4F-mRNA-PABP complex recruits the 43S, leading to formation of the 48S initiation complex (48S)^1^. The assembled 48S scans along the 5′ UTR of mRNA until it recognizes and selects a start codon^1,2^, triggering the release of most eIFs (late-stage 48S) and allowing the GTPase eIF5B to bind and promote joining of the 60S large subunit and formation of the 80S initiation complex (80S IC)^3–7^. GTP hydrolysis by eIF5B leads to its release from the 80S IC^8,9^, marking the end of translation initiation and the beginning of peptidyl bond formation^9^.

Not all translation initiation events follow this canonical process. Translation can also initiate at non-AUG codons, including those associated with upstream open reading frames (uORFs)^10–17^ or via internal ribosome entry sites (IRES)^18^. One example of such an alternative mechanism is repeat-associated non-AUG (RAN) translation^19^. RAN translation occurs in nucleotide repeat expansion diseases, where instability at DNA short tandem repeats (STR) underlies over 50 different neurodegenerative disorders^20–23^. A well-described example is the hexanucleotide G_4_C_2_ repeat expansion within the first intron of the *C9orf72* (chromosome 9 open reading frame 72) gene. While the general population typically has <30 G_4_C_2_ repeats, expansion of this STR is the leading monogenic cause of both Amyotrophic Lateral Sclerosis (ALS) and Frontotemporal Dementia (FTD)^24,25^. Despite its location within an intron, the G_4_C_2_ repeat is transcribed as part of a linear, capped mRNA that recruits the ribosome and supports RAN translation to generate toxic dipeptide repeat (DPR) proteins^26–28^. The expanded *C9orf72* STR can undergo bidirectional transcription, generating both sense (G_4_C_2_) and antisense (C_4_G_2_) transcripts^20–23^. RAN translation of these transcripts in all three reading frames generates five distinct DPRs: glycine-alanine (GA), glycine-proline (GP), and glycine-arginine (GR) from the sense strand, and proline-arginine (PR), proline-alanine (PA), and an additional GP from the antisense strand^29^. These DPRs accumulate in ALS and FTD patient brains and spinal cords^26,27,30,31^ and induce neurodegeneration in model systems^4,28,32,33,36^, suggesting an active role in disease pathogenesis. RAN translation is most efficient in the GA reading frame^28,36–38^ and GA DPRs are the most abundant in human tissues, whereas GP and GR DPRs are produced less efficiently and accumulate at significantly lower levels^39^.

RAN translation is proposed to initiate in multiple reading frames at both near-AUG cognate and non-cognate codons, circumventing several principles of canonical initiation. How G_4_C_2_ repeats interact with the ribosomal complex to promote translation initiation remains unclear. To address this fundamental question, we used singleparticle cryo-electron microscopy (cryoEM) to determine the structure of human late-stage C9orf72 translation initiation complexes after start codon selection. The structures of an *in vitro* assembled complex and a near-native complex purified from human cell lysate show that RAN translation initiates predominantly at a near-cognate CUG codon located upstream of the repeat region within a strong Kozak context. Notably, these structures also reveal a direct interaction between the G_4_C_2_ repeats and 18S rRNA. Blocking this interaction with antisense oligonucleotides reduces DPR production in *in vitro* translation lysate systems, plated neurons and patient-derived *C9orf72*-ALS iNeurons expressing endogenous DPRs. Together, these findings support a potential role for this RNA–RNA contact in promoting RAN translation and identify it as a novel therapeutic target for *C9orf72* RAN translation.

## Structures of human late-stage C9orf72 translation initiation complexes

To generate a disease-relevant translation initiation complex, we designed a capped and polyadenylated mRNA construct containing the first intron of the *C9orf72* gene and seventy G_4_C_2_ repeats, positioned 162 nucleotides downstream of the transcription start site (Extended Data Table 1). This mRNA models an aberrant transcript we recently identified as the major endogenous template for RAN translation in *C9orf72*-ALS patients^28^ and was previously demonstrated to support RAN translation across multiple reading frames^36,40^. Previous studies have established that recruitment of the C9orf72 mRNA to the 43S complex can occur through an eIF4F-dependent mechanism^36,37^. Therefore, we assembled reconstituted late-stage RAN initiation complexes using purified eIFs, PABP, tRNA_i_^Met^, and the 40S small ribosomal subunit (Extended Data Fig. 1), following methods previously established for canonical AUG translation initiation^41,42^.

Using cryoEM, we determined the structure of a late-stage RAN translation initiation complex at an overall resolution of 2.8 Å (Extended Data Fig. 2). Within the cryoEM map, we assigned several known components, including the 40S ribosomal subunit, eIF1A, eIF2, tRNA_i_^Met^, and mRNA (Fig. 1a-f; Extended Data Fig. 2). Notably, the cryoEM reconstruction revealed an additional density on the head of the ribosome directly interacting with the 18S rRNA expansion segment 9 (ES9^S^). To improve the local resolution of critical regions of the initiation complex, we performed focused 3D classification with a mask targeting the ternary complex, followed by additional focus classification on the unassigned density at ES9^S^ (Extended Data Fig. 2). Consequently, these focused classifications enriched for particles containing the TC and ES9^S^-associated density at the expense of other components of the initiation complex, including eIF3, resulting in reconstructions with low occupancy of these factors but improved local resolution of ES9^S^ and the TC. To account for molecular motions and enhance the quality of the cryoEM reconstruction, we used DynaMight, implemented in Relion-5^43^. This approach generated a refined map with an overall resolution of 3.2 Å (Extended Data Fig. 3 and Extended Data Table 2). While global resolution was slightly reduced, the improved local resolution around ES9^S^ allowed us to distinguish between purines and pyrimidines, enabling us to build the ES9^S^-associated density as G_4_C_2_ repeat RNA.

**Figure 1:**
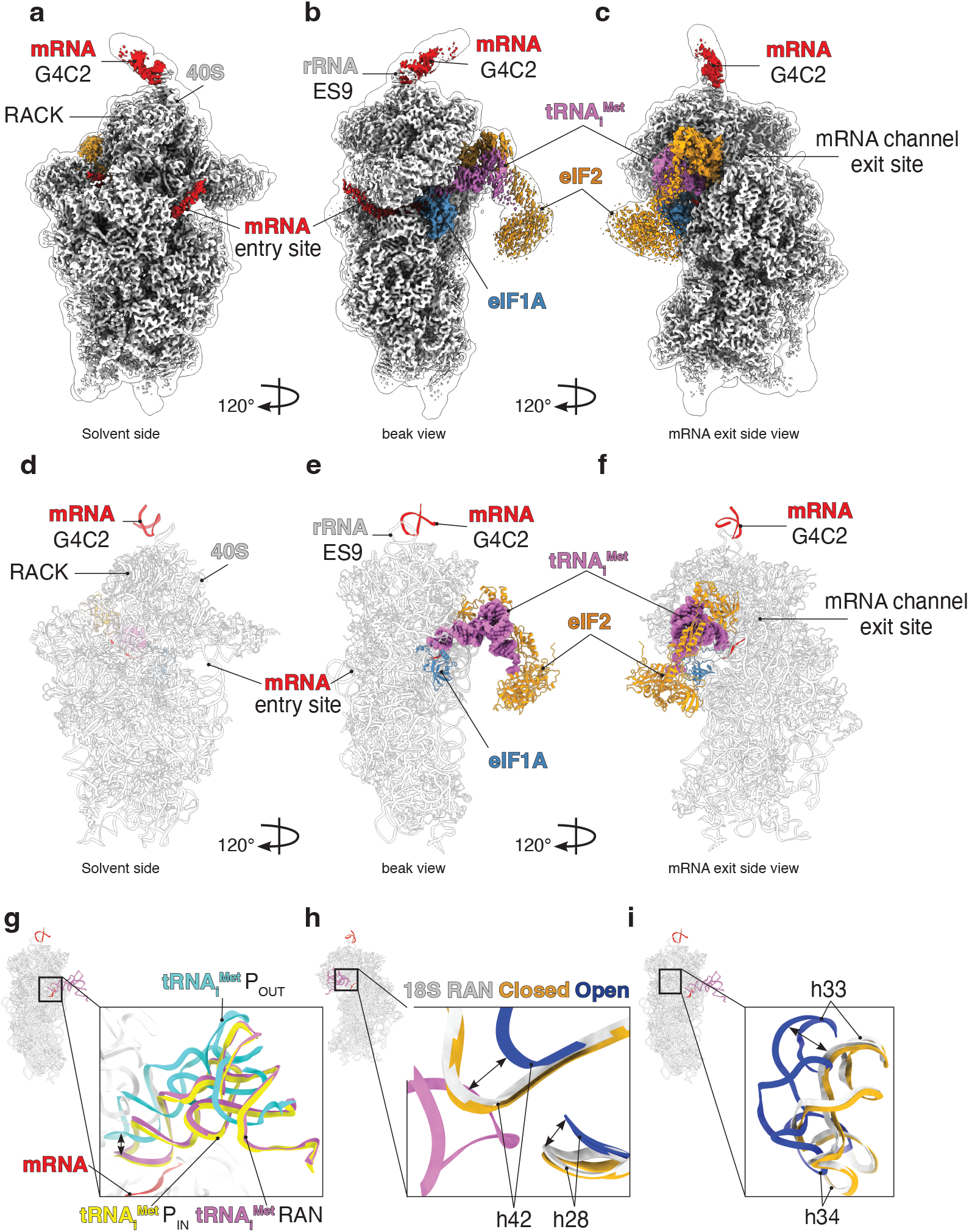
Structure of the *in vitro* late stage RAN translation initiation complex. **a-c**, Overview of the electron density map of the structure presented at the solvent side, the beak and the mRNA exit channel. **d-f**, Molecular model fitted within the electron density map. **g**, Superposition of the tRNA_i_^Met^ position between a canonical scanning 48S complex (cyan P_out_, PDB: 8PJ1) and a canonical late-stage 48S initiation complex (yellow P_in_, PDB: 8PJ2) reveals the late-stage RAN translation initiation complex is in a closed conformation (magenta). **h**, Comparisons of 18S rRNA head position in the RAN (light grey), open (blue) and closed (gold) 48S complexes at the posterior of the decoding center and **i**, at the beak further supported a RAN translation initiation complex in the closed position.

During the scanning of the 5′ UTR of mRNA, the head of the 40S small ribosomal subunit adopts an open conformation (away from the body)^42,44,45^, allowing tRNA_i_^Met^ to sample the codon in the ribosomal P-site (P_OUT_) until a start site is recognized and selected. Here, we observe that the overall conformation of the structure is a post-scanning intermediate in which the head of the 40S ribosome is in the closed conformation and the tRNA_i_^Met^ is fully inserted into the P site (P_IN_) (Fig. 1g-i). The conformation and the composition of this complex are consistent with that of a late-stage initiation complex, in which most eIFs, particularly eIF1, are released following start codon selection^41,44–50^ to allow for 60S large subunit joining.

While our *in vitro* reconstituted structure enabled us to examine translation initiation under controlled conditions, reconstitution may bias structural observations based on the specific factors included in the reaction. This is particularly relevant to RAN translation, which has been reported to proceed through both cap-dependent and cap-independent mechanisms^38,51–53^. To address this limitation and determine whether the structural features observed *in vitro* are recapitulated in a more physiologically relevant context, we determined the structure of a near-native late-stage RAN translation initiation complex isolated from human cell lysate at an overall resolution of 3.5 Å (Extended Data Fig. 4–6). This complex closely recapitulates the conformational features observed in our *in vitro* reconstituted structure, including a late-stage initiation state in which the tRNA_i_^Met^ is fully accommodated in the P site (P_IN_) (Extended Data Fig. 4k–l). To account for compositional heterogeneity within the dataset, we performed additional 3D classification, which resolved distinct particle populations, including a class containing eIF3 at 4.76 Å resolution and a class containing the G_4_C_2_-ES9^S^ interaction (Extended Data Fig. 4a–j and Extended Data Fig. 5).

### RAN translation on C9orf72 mRNA initiates at a near-cognate CUG codon

In the *C9orf72* gene, the G_4_C_2_ repeat expansion is positioned upstream of the annotated coding sequence (CDS)^24,25^, suggesting that RAN translation initiates at an upstream, non-canonical start site. In both the *in vitro* reconstituted and near-native complex structures, we were able to clearly distinguish purines and pyrimidines within the mRNA codon positioned in the P site of the ribosome and directly interacting with the anticodon stem-loop (ASL) of tRNA_i_^Met^. In both cases, *C9orf72* RAN translation initiates at a near-cognate CUG codon located at position −24 relative to the first nucleotide of the G_4_C_2_ repeat region (Fig 2a-c). This start codon is in the GA reading frame, which is the most abundant DPR detected in both reporter systems and patient brains (Extended Data Fig. 7)^36,39,54–56^.

**Figure 2:**
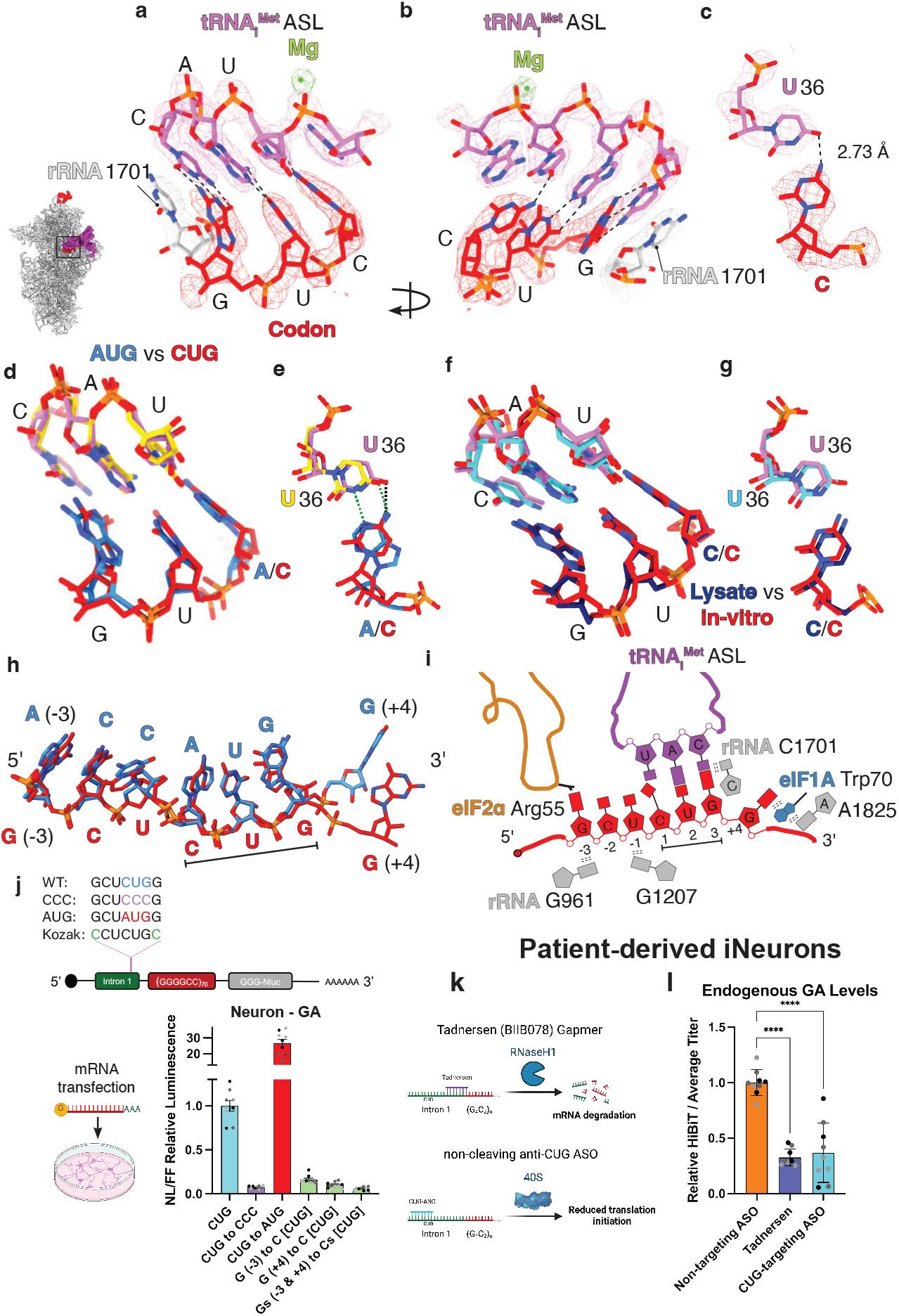
*C9orf72* RAN translation initiates at a CUG near-cognate codon and relies on a strong Kozak context. **a-b**, Structural overview of the decoding center in the C9orf72 RAN translation initiation complex shown in two orientations. **c**, Close-up view of U36 within the anticodon stem-loop (ASL) and the C in the CUG near-cognate start codon. A single hydrogen bond is observed between U36[O4] and C[N4], with the interatomic distance indicated in black. **d**, Superposition of the RAN translation initiation complex (colored in red and magenta) with the canonical 48S initiation complex (colored in dodger blue and yellow; PDB:8p09)^63^. **e**, Superposition of U36 in the ASL from RAN versus canonical initiation complexes. **f-g**, Comparison of the decoding center between our *in vitro* RAN late-stage initiation structure and our near-native late-stage RAN initiation structure acquired from HeLa cell lysate. Both structures reveal CUG codon usage with a single hydrogen bond formation. **h**, Superposition of mRNAs from the *in vitro* RAN late-stage initiation complex (red) and the canonical 48S complex (dodger blue; PDB: 8p09). **i**, Schematic representation of the molecular interactions that stabilize the Kozak sequence during RAN translation initiation. Hydrogen bonds are indicated by pairing bases, stacking interactions are indicated by parallel dashed lines between bases. **j**, Functional analysis of start codon fidelity: mutation of the near-cognate CUG codon to CCC, AUG or mutation of the Kozak sequence at −3 and/or +4 from G to C and subsequent mRNA transfection in rat hippocampal neurons in the GA reading frame. **k-l**, Patient-derived iNeurons obtained from Hölbling et al.^60^ were used to evaluate the endogenous levels of dipeptide repeats (DPRs) produced when treated with (**k**) Tadnersen, a gapmer that causes target RNA degradation, a non-cleaving CUG-targeting ASO or a non-targeting ASO at 5 µM final concentration. (**l**) Endogenous GA DPR production was evaluated using relative HiBiT signal and was normalized to average titer values. Error bars represent the mean ± SD (n=9) obtained from three independent replicates. Statistical significance: ****p <0.0001.

During RAN translation initiation, the near-cognate CUG codon occupies the same position in the P site of the ribosome as a canonical AUG codon (Fig. 2d). We assigned a C in the first position rather than an A by comparing the electron density between our RAN near-native late-stage initiation structure map and a canonical AUG map, which indicates that an A is incompatible when modelled into the RAN density (Extended Data Fig. 8a-d). However, the C at the first codon position is shifted by ~1 Å, allowing its N4 atom to form a hydrogen bond with the O4 atom of U36 in the ASL of the tRNA_i_^Met^. This interaction results in a cis Watson-Crick-Franklin/Watson-Crick-Franklin base pair (Fig. 2c, 2e), a non-canonical pairing previously observed in RNA-containing macromolecular complexes, including ribosomal structures^57^. Decoding of the first codon base is further facilitated by a subtle repositioning of U36 toward the center of the ASL (Fig. 2e). This conformational adjustment is stabilized by a magnesium ion bound to the phosphate backbone of the ASL (Fig. 2a,b), a feature not observed in structures of canonical AUG initiation^41,44–50^. Both the ~1 Å shift and formation of a single cis Watson-Crick-Franklin/Watson-Crick-Franklin base pair were observed in the *in vitro* reconstituted and native structures (Fig. 2f-g, Extended Data Fig. 8a-d).

To functionally validate the role of this CUG codon in RAN translation initiation, we transfected HeLa cells, HEK293 cells and primary rat hippocampal neurons with *in vitro* transcribed C9orf72 mRNA containing 70 G_4_C_2_ repeats. Consistent with our structure and most prior research^36–38,51,58,59^, mutation of the near-cognate CUG to the non-cognate CCC resulted in a significant reduction in RAN translation in the GA and GP reading frames but had minimal effect on the GR reading frame (Fig. 2j, Extended data Fig. 8e-k). To further evaluate the role of this CUG codon in a disease-relevant model system, we used patient-derived *C9orf72*-ALS iNeurons, described by Hölbling et al., that possess a HiBiT tag within the endogenous locus of the *C9orf72* gene^60^. These cells contain hundreds of G_4_C_2_ repeats at the endogenous *C9orf72* gene locus and provide a quantitative readout of endogenous DPR production in the GA frame. We treated these cells with a non-cleaving ASO targeting the near-cognate CUG codon, which significantly reduced endogenous GA production (Fig 2k-l). This result is consistent with prior work identifying non-cleaving ASOs targeting the near-cognate CUG codon as an effective strategy for reducing DPRs^37^. Together, these results support that the near-cognate CUG codon is a primary site of endogenous GA RAN translation initiation from C9orf72 mRNA, which is consistent with recent studies in patient-derived iNeurons where mutation of this CUG codon resulted in reduced production of both GA and GP DPRs^53^.

### *C9orf72* RAN translation initiation at a near-cognate CUG codon is promoted by a strong Kozak sequence

In canonical translation initiation, the efficiency of start codon recognition and selection is strongly influenced by the nucleotide context surrounding the AUG codon, known as the Kozak sequence (**G**/**A**NNAUG**G**). In mammals, deviations from this consensus sequence can markedly reduce the efficiency of start site selection^61,62^. Previous work demonstrated that mutating the Kozak sequence surrounding the near-cognate CUG codon in *C9orf72* significantly reduced RAN translation in a rabbit reticulocyte lysate (RRL) system, suggesting that the Kozak sequence promotes RAN translation^37^. Our structures support these findings and reveal the molecular interactions through which the Kozak context promotes selection of the near-cognate CUG codon during RAN translation initiation (Fig. 2h-i, Extended Data Fig. 9a-c).

eIF2, a central factor in translation initiation, not only delivers the tRNA_i_^Met^ to the ribosome but also contributes to the stringency of start codon selection^1^. Our structure reveals that the eIF2α residue Arg55 directly interacts with the Kozak sequence by stacking with the G nucleotide at the position −3 relative to the CUG codon (Extended Data Fig. 9a). This interaction is further stabilized by rRNA nucleotide G961. Additionally, the U nucleotide at position −1 is stabilized by an interaction with rRNA G1207 (Extended Data Fig. 9b), further anchoring the CUG codon within the P site of the ribosome. On the 3^*1*^ side of the CUG codon, the +4 G nucleotide interacts with eIF1A residue Trp70 and rRNA A1825 in helix 44 (Extended Data Fig. 9c), contributing to start site selection. These interaction networks closely resemble those observed during canonical AUG-mediated initiation^44,48,63^. Indeed, the spatial arrangement of the nucleotides surrounding the CUG codon closely resembles that observed in canonical late-stage initiation complexes. The +4 G adopts a slight shift in position, a feature also observed among structures of canonical 48S complexes (Fig. 2h)^41,44,46,48,49,63,64^. Thus, similar to AUG-mediated initiation, near-cognate CUG selection during C9orf72 RAN translation is supported by interactions between the surrounding Kozak sequence and the translation initiation machinery (Fig. 2i).

To functionally validate our structural findings, we generated constructs in which the Kozak sequence surrounding the near-cognate CUG was disrupted by mutating the −3 position from G to C, the +4 position from G to C, or both positions simultaneously. Mutating either the −3 or +4 position, or both positions together, significantly reduced RAN translation in the GA frame to levels comparable to those observed upon mutation of the CUG to CCC (Fig. 2j, Extended data Fig. 9d, 9f). Conversely, replacing the near-cognate CUG with a canonical AUG resulted in an approximately 25-fold increase in GA-frame DPR production, further supporting a model in which the CUG is recognized within a canonical-like Kozak context but initiates translation with substantially lower efficiency than AUG (Fig. 2j). Consistent with our analysis of CUG to CCC mutations, mutation of the −3 and +4 positions had limited impact on GR frame RAN translation in our system (Extended data fig. 9e, 9g-h). Collectively, our structural and functional data support a model in which GA-frame DPR production is predominantly initiated at the near-cognate CUG and depends on both an intact CUG codon and a favorable Kozak context for efficient RAN translation initiation.

Notably, ~10% of GA-frame RAN translation persisted following disruption of either the CUG codon or its Kozak context, bringing its expression in line with that observed basally for RAN translation in the GR reading frame (Fig. 2j, Extended Data Fig. 7). Additionally, neither the CUG nor Kozak mutations substantially affected RAN translation in the GR reading frame, at least under our experimental conditions. This continued translation is not easily explained by use of other alternative near-cognate codons^36^ or by translational frameshift events^37,65,66^. This residual translation suggests that a minor, but potentially important, fraction of DPR production occurs independent of the identified CUG start site, potentially through an IRES-like initiation pathway^38,40^.

### Ribosomal scanning in RAN translation initiation

The mechanism by which RAN translation initiates remains unresolved. One proposed model suggests that RAN translation proceeds through a canonical cap-dependent scanning pathway^67^. In this model, the repetitive RNA is first engaged by the cap-binding complex eIF4F, which facilitates recruitment of the 43S pre-initiation complex to form a 48S initiation complex. The ribosome then scans the mRNA and encounters the structured repeat region, which acts as a physical barrier to scanning, thereby increasing the dwell time of tRNA_i_^Met^ at near-cognate codons proximal to the repeats^67,68^ and ultimately promoting initiation at these sites. This model is consistent with the well-established role of downstream RNA hairpin structures promoting translation initiation at near-cognate start codons^69,70^. Moreover, it is consistent with the observation that RAN translation is largely cap-dependent^36–38,67^. An alternative model proposes that the repeat-containing mRNA directly recruits the ribosome, enabling initiation through an IRES-like mechanism^20,38,40,71^. Consistent with the latter model, RAN translation is upregulated during the integrated stress response (ISR)^36,40,72,73^, a condition in which canonical translation initiation is broadly downregulated^74,75^, and is enhanced by selectively disrupting eIF4E function in neurons^53^.

Our near-native cryoEM analysis of the C9orf72 RAN translation initiation complex identified a distinct class of particles corresponding to a scanning intermediate (Extended Data Fig. 10). This structure is characterized by the 40S subunit in an open conformation, mRNA positioned within the mRNA channel, and initiator tRNA_i_^Met^ in the P_OUT_ conformation (Extended Data Fig. 10k–l), all hallmarks of a scanning-competent state. As observed for the late-stage native initiation complexes, the scanning intermediate was resolved in particle classes both with and without a well-resolved G_4_C_2_–ES9^S^ interaction and eIF3 (Extended Data Fig. 10a–j).

### G_4_C_2_ repeat RNA binds to 18S ribosomal RNA expansion segment 9 (ES9^S^)

Our cryoEM structures (Fig. 1, Extended Data Fig. 4) reveal a direct interaction between the G_4_C_2_ repeat RNA and the ribosome. The repeat RNA binds to the 18S rRNA ES9^S^, located in the head region of the 40S subunit (Fig. 1a–f). Although this region of the ribosome is inherently flexible, focused 3D refinement with a local mask enabled reconstruction of a cryoEM map with sufficient local resolution (2.5 – 4.0 Å) to assign well-resolved nucleotide bases and discriminate between purines and pyrimidines in lower-resolution regions. This enabled us to build a complete atomic model of ES9^S^ and a partial model the G_4_C_2_ repeat RNA structure (Fig. 3a-f).

**Figure 3:**
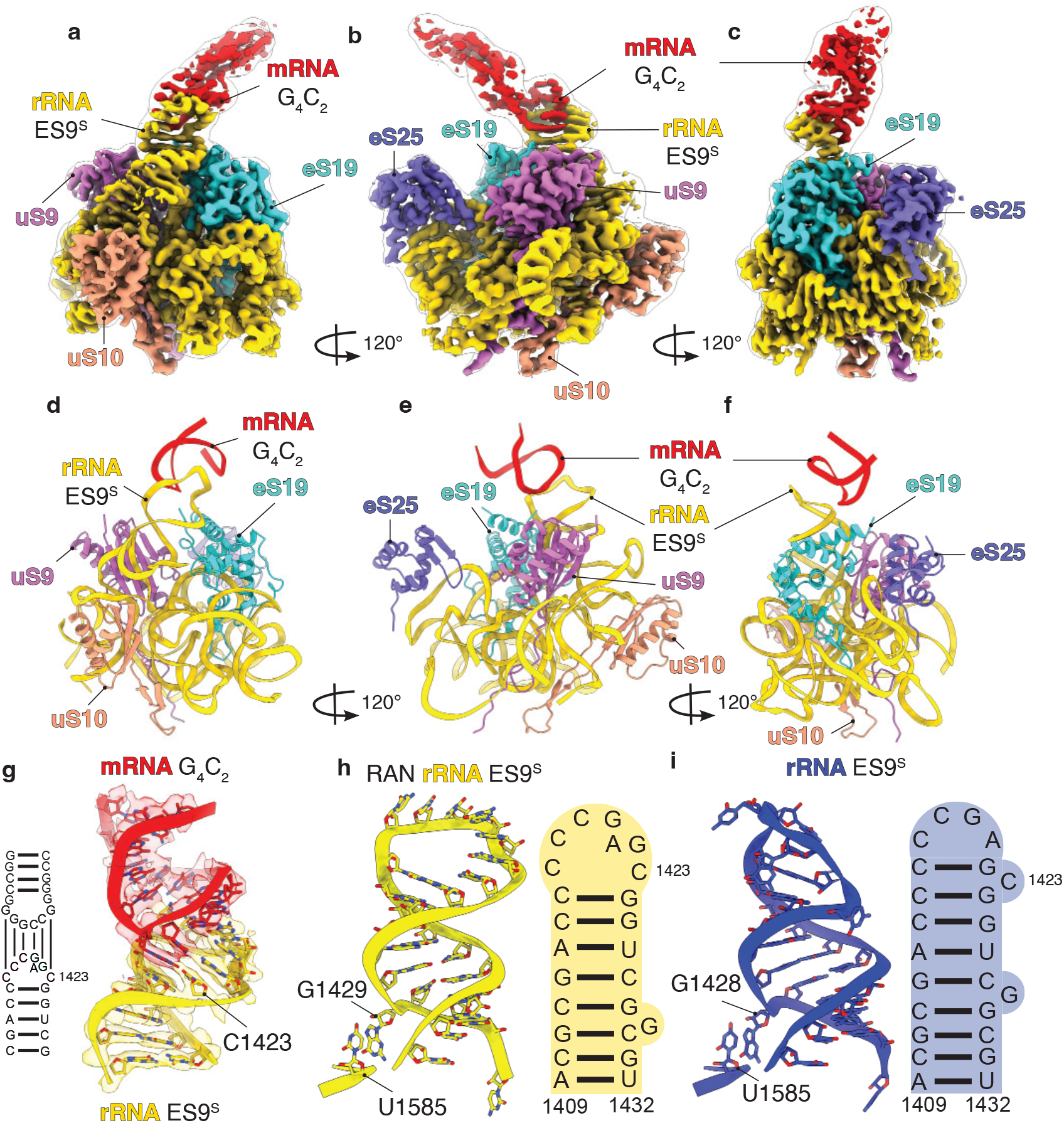
Structural basis of the interaction between 18S rRNA expansion segment 9 (ES9S) and G4C2 repeat mRNA. **a**–**c**, Overview of the cryoEM map of the ES9^S^–G_4_C_2_ mRNA region in the *in vitro* reconstituted human 48S RAN translation initiation complex shown in multiple orientations. The 18S rRNA is shown in gold, G_4_C_2_ repeat mRNA in red, and surrounding ribosomal proteins eS19 (turquoise), uS9 (orchid), uS10 (light salmon) and eS25 (purple) are highlighted. **d**–**f**, Atomic model view in the same orientations. **g**, Structure of ES9^S^–G_4_C_2_ fitted into the cryoEM map, highlighting the helical tails of the RNAs. **h-i**, Comparison of the ES9^S^ structure in the context of the RAN translation initiation complex (gold) versus the canonical 40S subunit lacking the G_4_C_2_ repeat (blue). The conformational rearrangement of ES9^S^ required for kissing-loop formation is evident. Each structure is accompanied by a schematic representation of the corresponding RNA secondary structure, illustrating the conformational differences.

G_4_C_2_ repeats can adopt both G-quadruplex and hairpin conformations *in vivo*^76–78^. Prior studies demonstrated that compounds which stabilize G-quadruplexes inhibit RAN translation, whereas increased expression of the G-quadruplex resolvase DHX36 promotes RAN translation^77–83^. Our structure reveals that, when bound to the ribo-some, G_4_C_2_ repeat RNA adopts a hairpin conformation and forms a kissing-loop interaction with the ES9^S^ region of the 18S rRNA, specifically involving nucleotides C1417 through C1423 (Fig. 3g). To investigate how RNA secondary structure affects RAN translation efficiency, we folded the C9orf72 repeat RNA under ionic conditions that favor G-quadruplex formation (high potassium)^77,84–86^ and compared it to RNA folded in ionic conditions that favor hairpin formation (high sodium)^77,86^. Consistent with our model, RNA folded under G-quadruplex-favoring conditions showed reduced RAN translation, whereas RNA folded under hairpin-favoring conditions maintained translation levels (Extended Data Fig. 11a-b). Further-more, heat-induced unfolding of the G-quadruplex prior to translation restored RAN translation (Extended Data Fig. 11c), supporting a structure-dependent inhibitory effect of G-quadruplex formation on RAN translation.

ES9^S^ is a conserved structural element of the ribosome. The C-rich segment that mediates contact with the G_4_C_2_ RNA is highly conserved across most mammals (Extended Data Fig. 12j), consistent with a potential functional role of the ES9^S^ region. This region is typically disordered in cryoEM structures, resulting in poor local resolution in previous structures of human ribosomes and precluding detailed structural analysis. To directly compare the native conformation of ES9^S^ to the conformation of ES9^S^ in the context of RAN translation initiation, we reanalyzed our previously published cryoEM dataset of the human 40S ribosomal subunit (EMD-47929)^87^. By applying focused 3D classification and refinement using a mask centered on the ES9^S^ region, we obtained a high-resolution structure that captures its native conformation (Extended Data Fig. 12a-i). This strategy enabled a detailed comparison of ES9^S^ before and after binding to the G_4_C_2_ repeat RNA hairpin loop (Fig. 3h-i and Supplementary Video 1).

In the native 40S ribosome structure, the ES9^S^ loop consists of four nucleotides (Fig. 3i). Upon binding to the G_4_C_2_ repeat RNA, the terminal G-C base pair within the loop is disrupted, resulting in widening of the loop (Fig. 3h) and the formation of six intermolecular base pairs with the repeat RNA (Fig. 3g, h). Accompanying the loss of the terminal G-C base pair, the entire ES9^S^ stem shifts upward by one base pair. Consequently, G1428, which normally participates in a stabilizing interaction with U1585 at the base of the ES9^S^ stem, is repositioned one nucleotide higher, while G1429 shifts into its place to establish a stacking interaction with U1585 (Fig. 3h, i and Supplementary Video 1). This structural rearrangement stabilizes the kissing-loop interaction between the repeat RNA and ES9^S^, potentially facilitating ribosome recruitment and promoting RAN translation initiation.

### The G_4_C_2_-ES9^S^ interaction promotes C9orf72 RAN translation and can be selectively inhibited

A major obstacle in studying the functional contribution of specific rRNA elements to translational control is the difficulty of directly manipulating rRNA. Mammalian cells contain hundreds of copies of rRNA genes^88^, making targeted genetic modification challenging. Moreover, perturbation of rRNA can have negative unintended consequences, including defects in ribosome biogenesis. Therefore, to assess the functional importance of the G_4_C_2_–ES9^S^ interaction without genetically altering the ribosome, we developed antisense oligonucleotides targeting ES9^S^ rRNA (ES9^S^-ASOs) designed to competitively disrupt the G_4_C_2_–ES9^S^ interaction (Fig. 4). We first validated direct binding of the ES9^S^-ASO to its intended ribosomal target by determining a 3.9 Å cryoEM structure of the 40S–ASO complex (Fig. 4a–d). As expected, the ASO binds directly to ES9^S^ and induces a structural rearrangement of ES9^S^ similar to that induced by the G_4_C_2_ repeat RNA in the RAN translation initiation structures.

**Figure 4:**
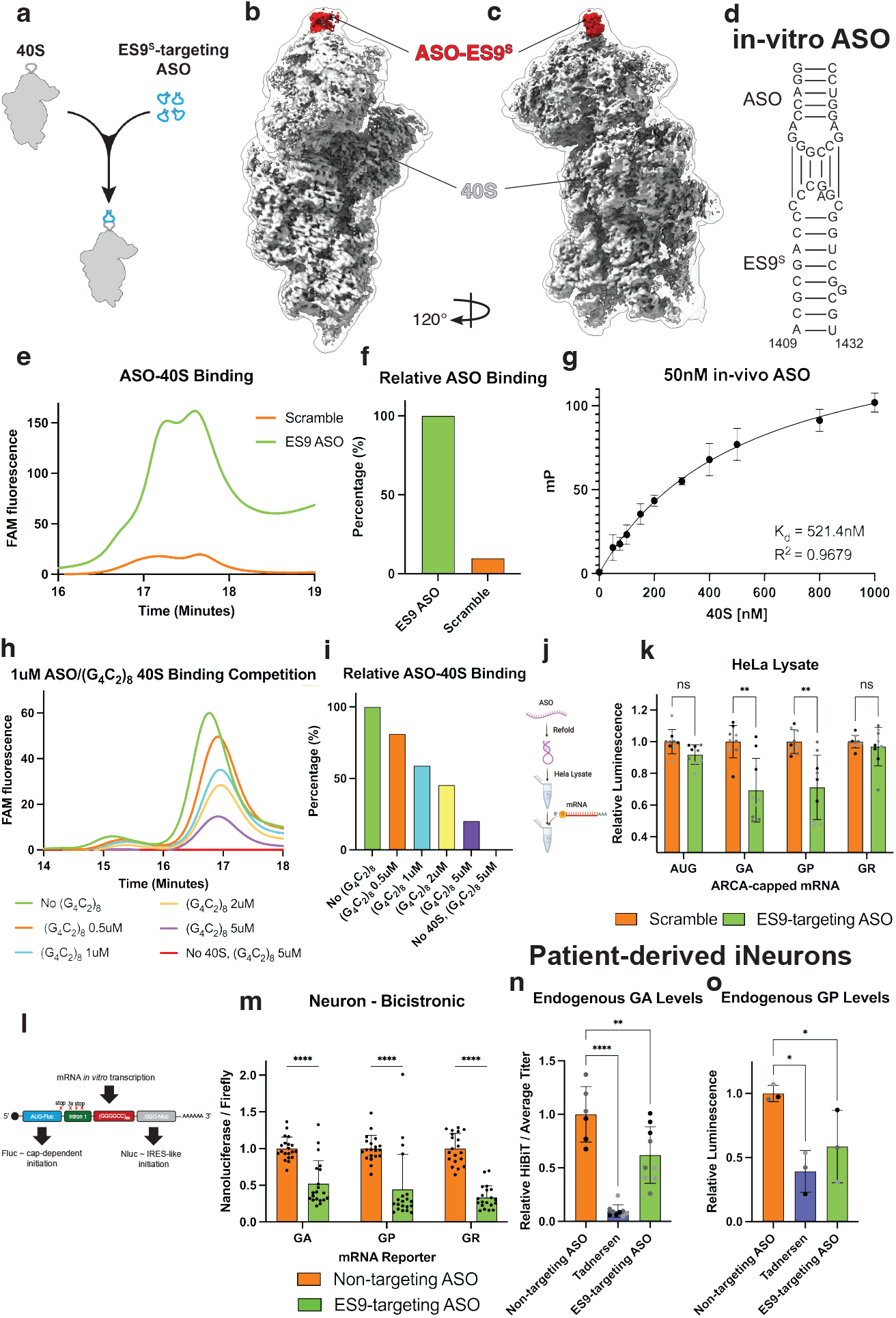
Antisense oligonucleotides (ASOs) targeting ES9^S^ selectively reduce *C9orf72* RAN translation via competitive inhibition of the ES9^S^-G_4_C_2_ kissing loop formation. **a**, Schematic representation of refolding the ES9^S^-targeting ASO and mixing with *in vitro* purified 40S ribosome. **b**–**c**, CryoEM structure of the ES9^S^-targeting ASO bound to the 40S ribosome at ES9. **d**, Schematic sequencing of the *in vitro* ASO-ES9^S^ binding interaction and the *in vivo* ASO. **e-f**, Assessment of 40S ribosome binding for a (FAM)-labelled ES9^S^-targeting ASO and a FAM-labelled scrambled ASO using an HPLC affinity analysis. Binding shown as (**e**) trace of the FAM signal over the 40S peak and (**f**) quantification of the area under the curve of the respective ASO’s FAM signals at the 40S peak. **g**, FAM-labelled *in vivo* ASO fluorescent anisotropy data for binding to the 40S ribosomal subunit identifies the ASO to have a K_d_ of 521.4 nM at 50 nM of ASO concentration. **h-i**, 1 µM *in vitro* FAM-labelled ES9^S^-targeting ASO mixed with *in vitro* purified 40S ribosome and varying concentrations of (G_4_C_2_)_8_ RNA oligonucleotide. 40S binding was then assessed using HPLC affinity analysis. Binding shown as (**h**) trace of the FAM signal over the 40S peak and (**i**) quantification of the area under the curve for each condition at the 40S peak. **j**, Schematic of ASO refolding for *in vitro* translation assays using cytoplasmic HeLa cell lysates. **k**, *In vitro* translation assays using cytoplasmic HeLa cell lysates and nanoluciferase reporters demonstrate that antisense oligonucleotides at 5 µM concentration reduce C9orf72 RAN translation in the GA and GP frames, but not the GR reading frame *in vitro*. Statistical significance: **p < 0.01. Error bars represent the mean ± SD (n=9) obtained from three independent replicates. **l-m**, Bicistronic reporters containing 66 repeats acquired from Ito et al.^59^ were *in vitro* transcribed into mRNA and subsequently transfected into rat hippocampal neurons and treated with 5 µM ASO, demonstrating reduction of IRES-like translation from the ES9^S^-targeting ASO in all three reading frames. Statistical significance: ****p < 0.0001. Error bars represent the mean ± SD (n=20) obtained from five independent replicates. **n-o**, Patient-derived iNeurons were used to evaluate the endogenous levels of dipeptide repeats (DPRs) produced after 5 µM of ASO treatment. **(n)** Endogenous GA DPR production evaluated using relative HiBiT signal and was normalized to average titer values. Error bars represent the mean ± SD (n=9) obtained from three independent replicates. Statistical significance: **p < 0.01, ***p < 0.001 **(o)** Endogenous GP DPR production evaluated using Meso Scale Discovery (MSD) Assay against the GP product. Statistical significance: *p < 0.05.

Next, we characterized binding of the ES9^S^-ASO to the 40S ribosomal subunit. We introduced a 3^*1*^ fluorescein phosphoramidite (FAM) label on the ASO (FAM-ASO) and measured its association with purified 40S subunits using a high-performance liquid chromatography (HPLC)-based separation assay, in which ASO binding was monitored by co-elution of the FAM signal with the 40S peak (Fig. 4e-f, Extended Data Fig. 13a-e). Consistent with our structural data, the ES9^S^-ASO bound selectively to the 40S subunit, exhibiting approximately 10-fold greater binding than an ASO containing a scrambled sequence (Fig. 4f-g). We next quantified the binding of the ES9^S^-ASO to the 40S ribosomal subunit through fluorescent anisotropy, identifying that the ASO binds the 40S at a K_d_ = 521.4 nM at 50 nM concentration of ASO. After quantifying our ES9^S^-ASO’s binding to the 40S, we tested whether G_4_C_2_ repeat RNA could compete with the ES9^S^-ASO for binding to the 40S subunit. Increasing concentrations of (G_4_C_2_)_8_ ssRNA resulted in a reciprocal decrease in ES9^S^-ASO binding to the 40S, supporting a model in which the G_4_C_2_ repeat RNA and ES9^S^-ASO compete for occupancy at ES9^S^ (Fig. 4h-i).

Having established that the ES9^S^-ASO can compete with G_4_C_2_ repeat RNA for occupancy at ES9^S^, we next examined its functional effect on RAN translation. We first assessed the effect of the ES9^S^-ASO on RAN translation of *in vitro* transcribed reporter mRNAs in HeLa cell lysates (Fig. 4j). RAN reporters exhibited a greater efficiency in the GA reading frame relative to GP and GR in HeLa lysates - consistent with their relative efficiencies in neurons (Extended Data Fig. 7d)^28,36,53^. In addition, these reporters exhibited a strong repeat-length dependence, with increased RAN translation at longer repeat lengths across all three reading frames except for the GR frame in rat hippocampal neurons (Extended Data Fig. 14a-b). Addition of the ES9^S^-ASO to the lysate had no significant effect on translation of an AUG-initiated nanoluciferase control construct relative to a scrambled control. In contrast, the ES9^S^-ASO significantly inhibited C9orf72 RAN translation in the GA and GP reading frames, although its effect on GR RAN translation in this context was not significant (Fig. 4j-k).

Since G_4_C_2_ repeat RNA interacts with the 40S ribosome in the absence of other translation factors^37^, we next assessed whether the ES9^S^-ASO might impact cap-independent RAN translation of G_4_C_2_ repeat RNA. We first tested this in HeLa lysates using repeat RNAs containing an Acap, which protects the mRNA from degradation, but is unable to engage the cap-binding protein eIF4E. RAN translation from these reporters was less efficient than from ARCA capped mRNAs, consistent with prior work^36,37^. However, addition of ES9^S^-ASO further diminished RAN translation in the GA and GP reading frames (Extended Data Fig. 14c), indicating that ES9^S^-dependent RAN translation in this lysate system is not restricted to a cap-dependent mechanism.

To extend this finding into a more physiologically relevant system, we used a series of recently described *in vitro* transcribed bicistronic constructs in which the upstream intronic region of C9orf72 and 66 G_4_C_2_ repeats are embedded between an AUG-FF-Luc and NLuc lacking an AUG start codon^51^ (Fig. 4l). When transfected into neurons, we detected NLuc expression from all three repeat-associated reading frames, although at levels two-to threefold lower than those driven by the cricket paralysis virus (CrPV) internal ribosome entry site (IRES) (Extended Data Fig. 14d). Given that the CrPV IRES itself exhibits relatively weak activity in human cells^89^, these results indicate that C9orf72 intron 1 containing an expanded G_4_C_2_ repeat possesses weak but detectable IRES-like activity. Notably, the CrPV IRES bicistronic mRNA initiates translation at an AUG codon, whereas the G_4_C_2_ bi-cistronic mRNA uses a near-cognate CUG codon, which we found to be approximately 25-fold less efficient than AUG (Fig. 2j). In these constructs, GA expression levels are closer to GP and GR levels than those observed with monocistronic reporters (Extended Data Fig. 7)^53^. We treated rat hippocampal neurons with an *in vivo*-compatible ES9^S^-ASO (Fig. 4d, Extended Data Table 1), then transfected them with the RAN bicistronic mRNA reporters. We observed a significant inhibition of RAN translation across all three reading frames compared to a scrambled control, further supporting a role for ES9^S^ in cap-independent translation of repeat mRNA (Fig. 4m).

Finally, to assess whether this G_4_C_2_-ES9^S^ interaction contributes to endogenous RAN translation, we used the *in vivo*-compatible ES9^S^-ASO in patient-derived iNeurons. Because the HiBiT reporter in these cells measures DPR production in the GA frame, we independently quantified GP DPR levels using a Meso Scale Discovery (MSD) assay that directly measures GP peptide abundance^77^. Consistent with our *in vitro* findings, the ES9^S^-targeting ASO significantly reduced endogenous DPR production in patient-derived iNeurons, decreasing both GA and GP levels (Fig. 4n–o). Collectively, these results demonstrate that disrupting the G_4_C_2_-ES9^S^ interaction with ES9^S^-targeting ASOs selectively suppresses the production of toxic C9orf72-derived DPRs and establishes ES9^S^ as a functionally important ribosomal determinant of RAN translation.

## Discussion

Defining the molecular features that enable RAN translation initiation is a significant challenge. Our structures and functional data reveal that C9orf72 RAN translation initiates predominantly at a near-cognate CUG codon located 24 nucleotides upstream of the G_4_C_2_ repeat region. This finding is consistent with prior biochemical studies indicating that this codon and its Kozak context are important for translation initiation, particularly in the GA reading frame^36,37^. The CUG codon is positioned within an optimal endogenous Kozak context, providing a rationale for its preferential selection over other near-cognate codons located upstream of the repeat. These findings do not preclude the possibility of additional initiation events occurring at other near-cognate or non-cognate codons, with potential contributions to translation in other reading frames. However, the favorable context of the −24 CUG provides a molecular explanation for why initiation at alternative sites may occur less efficiently and generally supports prior work suggesting that ribosomal frameshifting on or in the repeat sequence might contribute to generation of such products^37,58,90,91^.

The role of repeat RNA structure in promoting the near-cognate initiation process has, until now, remained enigmatic. Patients with C9orf72-associated neurodegenerative disease carry more than 30 G_4_C_2_ repeats in the STR region^24,92,93^. Indeed, our functional data in both rat hippocampal neurons and HEK293 cells show that increasing repeat length leads to an increase in RAN translation (Extended data Fig 14a-b). Our structural data provides a potential mechanistic explanation for this requirement – a long-range RNA–RNA interaction between the G_4_C_2_ repeat RNA and the ES9^S^ stem loop of the 18S rRNA. ES9^S^ is located near the mRNA channel exit site on the head of the 40S subunit. Based on our structural model, a G_4_C_2_ stem loop would require a minimum of ~30 repeats to span from the mRNA channel entry site to the region where it binds ES9^S^ (Extended Data Fig. 15). In our near-native dataset, we identified that the ES9^S^-G_4_C_2_ interaction is observed in both scanning and initiated structures (Extended Data Fig. 4, Extended Data Fig. 10). This supports a model in which the ES9^S^-G_4_C_2_ kissing loop can form during scanning and initiation, potentially occurring during ribosome recruitment. This restraint provides a structural rationale for how the repeat may play a role in ribosome recruitment during RAN translation.

RAN translation can also proceed via alternative initiation pathways that bypass the requirement for the cap-binding complex eIF4F in at least some contexts^38,40,53^. For example, constructs containing stop codons in all three reading frames upstream of the G_4_C_2_ repeat – and downstream of the near-cognate CUG codon – still support RAN translation, albeit at reduced efficiency^38,40,52,94^. Furthermore, RAN translation is enhanced under conditions that inhibit cap-dependent translation, such as activation of the integrated stress response (ISR)^36–38,40,72^. Consistent with this prior work, we observe that *in vitro* transcribed RNA synthesized from bicistronic reporters of C9orf72 RAN translation translates efficiently in all three reading frames when transfected into primary rat neurons (Extended Data Fig. 14d). As such, our findings support the existence of alternative recruitment pathways in addition to eIF4F-dependent initiation. It is thus intriguing that the ES9^S^-ASOs reduce RAN translation from both RNAs that lack a functional mRNA cap, which cannot efficiently bind eIF4F, and *in vitro* transcribed bicistronic reporters (Fig. 4l-m, Extended Data Fig. 14c). We therefore propose that the G_4_C_2_–ES9^S^ kissing-loop interaction may directly facilitate ribosome recruitment in the absence of canonical cap-binding factors and thereby contribute to an alternative, cap-independent initiation pathway.

Ribosomal expansion segments have historically been viewed as inert rRNA extensions, but a growing body of evidence has revealed discrete biological functions for these elements, including a recent report demonstrating that expansion segments promote disome formation during stress in neurons^95^. Our findings place ES9^S^ within this emerging functional framework and define, at nucleotide resolution, a sequence-specific RNA–RNA interaction between G_4_C_2_ repeat RNA and ES9^S^. A previous cryoEM analysis of the mouse *Hoxa9* RNA element identified low-resolution density corresponding to mRNA in direct contact with ES9^S^. Although the resolution of that reconstruction was insufficient to define the sequence underlying the ES9^S^–mRNA interaction, the *Hoxa9* element is GC-rich and contains a “GGGCCG” motif that the authors proposed to reside at the exposed end of a stem loop^96^. Notably, this motif closely corresponds to the ES9^S^ binding sequence identified here within the RAN mRNA. Structural comparison further shows that the two RNA elements superimpose at their point of contact with ES9^S^ and diverge only as they extend away from the 40S subunit, consistent with the two stem loops approaching ES9^S^ from slightly different orientations (Extended Data Fig. 16).

Together, these two examples of mRNA engagement with ES9^S^, along with our structure of an ASO bound at the same site, support a broader model in which the ES9^S^ may serve as a sequence-specific interaction surface for structured mRNAs. The G_4_C_2_-ES9^S^ kissing loop observed in our structures of C9orf72 RAN translation initiation may therefore represent one example of a broader mode of direct mRNA-rRNA recognition rather than a mechanism unique to RAN translation. Determining the prevalence of ES9^S^ engagement across the translatome, identifying the sequence and structural features that govern these interactions, and establishing whether ES9^S^-mediated recognition serves a regulatory function under specific cellular conditions represent important directions for future investigation.

Expanded G_4_C_2_ repeats can adopt multiple RNA conformations, and our structures do not exclude the formation of G-quadruplexes elsewhere within the repeat sequence. A recent preprint proposed that G-quadruplex formation is required for RAN translation initiation on expanded G_4_C_2_ repeats^97^. However, extensive evidence indicates that G-quadruplexes generally act as structural barriers to ribosomal scanning and elongation, thereby inhibiting translation^78–80,98–103^. Consistent with this inhibitory role, efficient translation of G-quadruplex-containing mRNAs often requires RNA helicases such as DHX36^79,80,101–104^, which has also been implicated in facilitating translation elongation through expanded G_4_C_2_ repeats^79,80^. In agreement with these findings, we observed reduced RAN translation when G_4_C_2_ repeat-containing RNA was folded under conditions favoring G-quadruplex formation, whereas translation efficiency was maintained under conditions favoring hairpin formation (Extended Data Fig. 11a,b). Importantly, our cryoEM structures reveal that the region of the G_4_C_2_ repeat that directly interacts with the ribosome adopts a hairpin conformation that forms a kissing-loop interaction with expansion segment ES9^S^ of the 18S rRNA. Together, these findings support a model in which expanded G_4_C_2_ repeats adopt distinct RNA conformations with opposing effects on translation: hairpin formation promotes RAN translation initiation through direct interactions with the ribosome, whereas G-quadruplex formation inhibits translation elongation, consistent with its established inhibitory effects on other G-quadruplex-containing transcripts^78–80,98–104^.

In summary, we present the first structures of a RAN translation initiation complex under *in vitro* and near-native conditions, revealing how the G_4_C_2_ repeat RNA physically binds the ribosome through a specific interaction with the ES9^S^ region of the 18S rRNA. Importantly, complementary antisense oligonucleotides that prevent this RNA-RNA interaction selectively suppress RAN translation in patient iNeurons. These findings suggest a potential therapeutic strategy for C9orf72-associated ALS and FTD, and possibly for other disorders in which ES9^S^-mediated mechanisms contribute to aberrant translation initiation.

## Methods

### Purification of Human Translation Initiation Factors and 40S Ribosomal Subunits

Translation initiation factors (eIFs) and tRNA_i_^Met^ were purified as previously described for canonical initiation^4,41,42,105–107^. Briefly, recombinant eIF1, eIF1A, eIF5, eIF4B, eIF4E, eIF4A and PABP were expressed in *E. coli* BL21(DE3) cells. Constructs included N-terminal His-MBP fusion tags followed by a TEV protease cleavage site. Proteins were purified using nickel affinity chromatography (HisTrap FF, Cytiva). Target proteins were eluted from the Ni-NTA resin using a linear imidazole gradient (0–250 mM). To remove the N-terminal affinity tag, TEV protease cleavage was performed overnight at 4°C in dialysis buffer containing 25 mM Tris (pH 7.5), 200 mM NaCl, 20 mM imidazole, 10% glycerol, and 5 mM 2-mercaptoethanol. The cleaved sample was subjected to a second Ni-NTA purification step to separate the His-MBP tag from the target protein. Subsequently, the protein was purified by ion-exchange chromatography using either HiTrap SP or HiTrap Q columns (Cytiva), selected based on the protein’s isoelectric point or localized surface charge. A linear KCl gradient (100–500 mM) was used for elution. Final polishing was achieved by size-exclusion chromatography on a Superdex 200 column. Protein purity was assessed by SDS-PAGE followed by Coomassie Brilliant Blue staining. The final protein sample was concentrated using Amicon centrifugal filters with a 10 kDa molecular weight cutoff. Native eIF3 was purified from Expi293F™ cells (ThermoFisher, A14527) using a combination of ammonium sulfate precipitation and sequential chromatography. Briefly, eIF3 was precipitated from clarified lysate using 40% ammonium sulfate saturation, and the precipitate was subjected to a three-step purification protocol consisting of Mono Q anion exchange, HiTrap Heparin affinity, and Superdex 200 size-exclusion chromatography. The same cells were used to purify the 40S small ribosomal subunit, using an established protocol^108^. This protocol was also used to purify the 40S small ribosomal subunit from the commercial cytoplasmic HeLa lysate (IPRATECH, CC-01-4050). Human eIF4G was expressed and purified as described before^106^. Briefly, eIF4G was expressed in Sf9 cells as a His_6_-tagged construct and purified using a combination of immobilized metal affinity, ion-exchange, and size-exclusion chromatography. Following elution from Ni-NTA Superflow resin, purified human eIF4A1 was added to eIF4G, and the proteins were incubated overnight at 4 °C to promote formation of the eIF4G–eIF4A heterodimer. Native eIF2 was purified using the same procedure as that used for native eIF3 purification, except that the ammonium sulfate concentration was increased to 50% during the precipitation step^109^. Briefly, the sample was purified from HeLa cell lysate using hydroxyapatite and size exclusion chromatography.

### *In Vitro* mRNA Transcription and Purification

For RNAs used in structural studies and functional studies, *in vitro* transcription was performed using linearized plasmid DNA (50 ng/µL) in transcription buffer (40 mM Tris-HCl pH 8.0, 30 mM MgCl_2_, 1 mM spermidine, 0.01% Triton X-100, 5 mM DTT) with 8 mM of each rNTP (ATP, GTP, CTP, UTP; Sigma), 15 µg/mL pyrophosphatase, 1.25 U/µL T7 polymerase (NEB M0251), and 1 U/mL RNasin Plus (Promega N2611), incubated at 37 °C for 3 hours. DNA was removed with DNase I (2.5 U/µL, NEB M0303) at 37 °C for 30 minutes, followed by ethanol precipitation. Pellets were resuspended in nuclease-free water.

Purified RNA was separated from abortive products by either denaturing PAGE or size exclusion chromatography. For PAGE, samples were mixed 1:1 with loading dye (NEB B0363S), heated to 95 °C for 2 minutes, and resolved on 4% acrylamide/7 M urea gels at 400 V overnight at 4 °C. RNA was visualized under 256 nm UV and electroeluted (Model 422 Electro-Eluter, Bio-Rad).

mRNA was capped using the Vaccinia capping system (NEB M2080) and polyadenylated using *E. coli* poly(A) polymerase (NEB M0276). Capped and tailed mRNA was cleaned using RNeasy Mini Kit (Qiagen 74104). RNA integrity and modification status were validated by denaturing gel electrophoresis. For size-exclusion purification, RNA was separated on a Superose 6 column (AKTA Pure™) with elution buffer (10 mM NH_4_OAc-HOAc, pH 5.0; 150 mM KCl; 5 mM MgCl_2_), then ethanol precipitated and processed as described. Column-purified GA70 RNA was used without refolding.

For RNAs used for RNA transfection in cells and *in vitro* functional studies, RNAs were synthesized using the HiScribe T7 High Yield RNA Synthesis Kit (NEB E2040S). They were co-transcriptionally capped either with a functional ARCA-cap (3′-O-Me-m7G(5′)ppp(5′)G RNA Cap Structure Analog, NEB S1411L) or a non-functional A-cap (m7G(5’)ppp(5’)A RNA Cap Structure Analog, NEB S1405) that stabilizes the RNA against degradation but does not allow for cap binding. RNA was then polyadenylated using *E. coli* poly(A) polymerase (NEB M0276) and cleaned using RNeasy Mini Kit (Qiagen 74104).

### RNA Refolding

Oligo refolding was conducted in 1× oligo refolding buffer (10 mM HEPES-KOH, pH 7.5; 100 mM KCl, 3mM MgCl_2_, 5% glycerol). RNA was heated to 95 °C for 4.5 minutes and cooled gradually: 65 °C (5 min), then decrementally at 60, 55, 50, 45, 40, 35, 30 °C for 1 minute each, followed by 25 °C for 30 minutes, and stored on ice for at least 30 minutes prior to use.

mRNA was folded based on a protocol modified from Junjie U Guo and David P Bartel^110^. Briefly, RNA was heated to 80°C for 2 minutes and cooled to 4°C for one minute in a solution containing 0.05M Tris (pH 7.5), 1mM MgCl_2_, and either 100mM NaCl (favoring hairpin folding), 150mM KCl (favoring G-quadruplex folding), or no additional salt.

### High-performance liquid chromatography (HPLC) Binding Assay

For the 40S ribosomal binding assessment assays, the FAM-labeled oligonucleotide was mixed with HEK293T 40S ribosome at a molar ratio of 3:1 (3 µM oligonucleotide and 1 µM of HEK293T 40S ribosome). The mixture was incubated at 30 °C for 10 min and then transferred to the HPLC autosampler at room temperature for injection. Size-exclusion chromatography was performed at room temperature using GTxResolve Premier SEC 1000 Å, 3 μm, 4.6 x 300 mm (Waters) at flow rate 0.2ml/min for 30min (Buffer: 20mM HEPES, 1mM DTT, 100mM KCl, 3mM MgCl_2_, pH 7.5). Fractions corresponding to 40S peak and free ASO were collected. The samples were run on 1% agarose gel in 1X Tris-acetate EDTA (TAE) buffer at room temperature at 180V for 30 minutes. Gels were either unstained and imaged in the FAM channel for identifying FAM signal or stained with 1 X GelGreen for RNA signal analysis. For the (G_4_C_2_)_8_ competition assay, 1 µM of the FAM-labeled oligonucleotide and 1 µM HeLa 40S ribosome were mixed with the (G_4_C_2_)_8_ at the specified concentrations (0.5 µM, 1 µM, 2 µM and 5 µM). The mixtures were incubated at 30 °C for 10 minutes. Size-exclusion chromatography was performed at room temperature using the same column, flow rate, run time and buffer as described above.

### Fluorescent Anisotropy Binding Assay

Fluorescent anisotropy (FA) experiments were conducted using an *in vivo* ES9^S^-targeting FAM-labelled ASO (FAM-ASO) at 50 nM in FA reaction buffer (20 mM HEPES, 100 mM KCl, 4 mM MgCl_2_, 1 mM DTT, pH 7.5). Serial dilutions of unlabeled 40S ribosomes (1,000, 800, 500, 300, 200, 100, 75, 50, 25, and 12.5 nM) in FA reaction buffer were added to PCR tubes containing the FAM-labelled ASO. Two controls were included: a FAM-ASO control (50 nM) and a buffer control. The samples were incubated at 30°C for 10 min, then immediately transferred to 384 well black plates and analyzed on a Tecan Spark^®^ Multimode Microplate Reader. Buffer control was used for blank and the ASO-only control was used for reference, G-factor was calibrated using blank and reference. mP readings were plotted as a function of 40S concentration and binding curve was calculated using the equation One site – Specific binding (Y = Bmax*X/(Kd + X)) in GraphPad Prism.

### *In Vitro* Translation Assay in Rabbit Reticulocyte Lysate (RRL)

RRL *In vitro* translation assays following RNA folding were conducted using the Flexi Rabbit Reticulocyte Lysate System (Promega, L4540), following the protocol as previously described^36^. In brief, folded RNAs were diluted to 6 nM into a reaction (10 μL) containing 30% RRL, 10 mM amino acid mixes, 0.5 mM magnesium acetate, and RNase Inhibitor (NEB). The additional salts in the RRL reaction that come along the diluted RNA from the folding reaction (3.75 mM KCl or 2.5 mM NaCl) were normalized across all experiments. The reaction was performed for 30 min at 30°C, then terminated by incubation on ice. After termination, reactions were diluted 1:7 with Glo Lysis Buffer (Promega, E2661), followed by the addition of NanoGlo Substrate freshly diluted 1:50 in NanoGlo Buffer (Promega, N1150). Luminescence was measured using a GloMax 96 Microplate Luminometer.

For data expressed in Extended Data Fig. 11c, folded RNA solutions were heated after dilution (so the available KCl is negligible) for the time and temperature indicated. The solution was allowed to equilibrate to room temperature for 15 minutes and then we proceeded with the RRL experiments as described above.

### *In Vitro* Translation Assay in HeLa Lysate

Translation was performed using commercial cytoplasmic HeLa lysate (IPRATECH, CC-01-40-50). Reactions (10 µL) contained: 5 µL lysate, 1 µL 10× translation buffer (final 1×: 16 mM HEPES-KOH, 20 mM creatine phosphate, 0.1 mg/mL creatine kinase, 0.1 mM spermidine, 50 mM KOAc), 1.75 µL refolding buffer (10 mM HEPES-KOH, 100 mM KCl, 10% glycerol), 0.25 µL RNasin Plus, 1 µL oligonucleotide or 1X refolding buffer (refolded as described above), and nuclease-free water as needed. All reactions with oligonucleotide were conducted at a final concentration of 5 µM oligonucleotide.

Reaction master mixes were prepared on ice and preincubated at 37 °C for 20 minutes. To initiate translation, 9 µL of master mix was combined with 1 µL mRNA in PCR tubes, gently mixed, and incubated at 37 °C for 30 minutes. Reactions were terminated by adding 10 µL of 4 µM puromycin in 2× passive lysis buffer (Promega E1910). Luciferase activity was measured using the Nano-Glo assay (Promega N1110), per manufacturer’s instructions. Translation was assessed at 24.6 nM RNA. Each dataset represents the mean of three independent biological replicates, each performed in technical triplicate.

### Cell Culture

HEK293 and HEK293T cells were cultured in high-glucose DMEM (Gibco, 11965118) supplemented with 10% fetal bovine serum (FBS; Bio-Techne, S11150) and 1% penicillin-streptomycin (Gibco, 15070063). HeLa cells were cultured in high-glucose DMEM supplemented with 10% FBS, 1% nonessential amino acids (NEAA; Gibco, 11140050), and 1% penicillin-streptomycin.

Primary rat hippocampal neurons were prepared from P0–P1 Sprague-Dawley rat pups. Hippocampi were dissected and dissociated using papain as previously described by Rodriguez et al.^111^. Neurons were plated at a density of 3.5 × 10^4 cells/well in poly-D-lysine-coated 96-well plates (Gibco, A3890401) in 200 µL/well of neuronal growth medium (NGM). NGM consisted of Neurobasal-A medium (Gibco, 10-888-022), B27 supplement (Gibco, 17-504-044), and GlutaMAX supplement (Gibco, 35-050-061) at a 100:2:1 ratio.

All animal procedures were conducted in accordance with National Institutes of Health guidelines and approved by the University of Michigan Committee on the Use and Care of Animals. All cells were maintained at 37°C in a humidified incubator containing 5% CO_2_.

### RNA Transfection and Luciferase Assay

Primary rat hippocampal neurons were plated in 96-well plates at 3.5 × 10^4 cells/well and cultured for 7 days prior to transfection. HeLa cells were plated at 1 × 10^4 cells/well, and HEK293 cells were plated at 2 × 10^4 cells/well, 24 h prior to transfection.

Primary rat hippocampal neurons, HEK293, and HeLa cells were transfected with reporter RNA and firefly luciferase RNA, which served as a transfection control, using Lipofectamine MessengerMAX reagent (Thermo Fisher Scientific, LMRNA008) according to the manufacturer’s protocol for *in vivo* NanoLuciferase assays. For CUG-to-CCC mutation experiments, Kozak mutation experiments and repeat-length NanoLuciferase assays (Fig. 2j, Fig. 4k, Extended data Fig. 7, Extended Data Fig. 8e-k, Extended Data Fig. 9d-h, Extended Data Fig. 14a-c), 1.23 nM of each of reporter RNA and 50 ng of firefly luciferase RNA was used.

We obtained bicistronic plasmids as a gift from Hideki Taguchi’s group (Ito et al. 2025, Fig 4l-m, Extended Data Fig. 14d). These constructs contain a Firefly luciferase reporter driven by an AUG start codon and stopping at a canonical stop codon. This is followed by partial 5′ UTR intron 1 of C9orf72 which contains stop codons in all three reading frames and is continuous with 66 G4C2 repeats and a nanoluciferase reporter with a “GGG” mutation of the start site of nanoluciferase. For our bicistronic reporter assays, we followed the same RNA amount protocol as above, with RNAs added at 2.53nM concentration.

Luciferase activity was measured 24 h after RNA transfection. Cells were lysed in 70 µL Glo Lysis Buffer (Promega, E26661) for 5 min on a rocker at room temperature. Lysates were then mixed 1:1 with either 1× Nano-Glo substrate prepared in Nano-Glo buffer (Promega, N1150) or One-Glo reagent (Promega, E6130) and incubated for 5 min on a rocker in the dark. Lysates were transferred to opaque 96-well plates. Luminescence was measured using a GloMax 96 Microplate Luminometer.

### Patient-derived iPSC maintenance and iNeuron differentiation

*iPSCs* were obtained from American Type Culture Collection (ATCC) as fibroblasts (ATCC PCS-201-010) and differentiated into iPSCs as in Maltby et al., 2024 (48). Poly-GA HiBiT iPSC line was obtained from Adrian Isaacs’s lab at UCL. Cells were cultured in mTeSR plus (Stem Cell, 100-0276) or TeSR-E8 (Stemcell 05990) and maintained at 37°C, 5% CO2.

*iNeurons* were differentiated as in Miller et al., 2025 (20). Briefly, a neurogenin-1 and neurogenin-2 (Ngn1/2) overexpression cassette was delivered into iPSCs. iPSCs were cultured in mTeSR plus (Stem Cell, 100-0276) or TeSR-E8 (Stemcell 05990) media plus 2μg/mL doxycycline for 2 days and frozen for further use. To begin iNeuron differentiation (day 1), cells were thawed onto dPGA/Geltrex coated plates in TeSR-E8 containing N2 with 1x NEAA supplement (Gibco 11140050), 10ng/mL BDNF, 10ng/mL NT3, 0.2ug/mL laminin, 2μg/mL doxycycline and 1ul/mL of 10mM Rock inhibitor. On day two, media was changed to transition media using half TeSR-E8 media, half DMEM F12 (Gibco 11320033) and containing N2 with 1x NEAA supplement (Gibco 11140050), 10ng/mL BDNF, 10ng/mL NT3, 0.2ug/mL laminin, 2μg/mL. On day three, media was changed to Neurobasal-A (Gibco 10888022) containing 1x Culture One, 1x B27 supplement (Gibco 17504044), 1x Glutamax supplement (Gibco 35050061), 10ng/mL BDNF, 10ng/mL NT3, 0.2μg/mL laminin, and 2μg/mL doxycycline. From day 6 until maturity (day 14), iNeurons were cultured in maintenance media which contained 1x B27 supplement (Gibco 17504044), 1x Glutamax supplement (Gibco 35050061), 10ng/mL BDNF, 10ng/mL NT3, and 0.2μg/mL laminin, in Neurobasal-A (Gibco 10888022). Cells were maintained at 37°C, 5% CO2 and underwent a 50% media change twice weekly.

### HiBiT Lytic Detection and Cell Titer Analysis Assay

Neural progenitor cells plated in 96 well plates were differentiated for 14 days as previously described^60^. After differentiation, iNeurons were treated with 5 µM of ASO and 0.2uL of Endoporter (GeneTools, OT-EP-PEG-1) at a final well volume of 100uL. On day 3 or 4 of ASO treatment, an additional 100uL of media alone was added to cells, diluting the effective ASO concentration to 2.5 µM. Cells were removed from the incubator on day 7 of ASO treatment and wells were either measured for HiBiT signal using the NanoGlo® HiBiT Lytic Detection System (Promega, N3040) per manufacturer’s protocol or assessed for Titer value by first lysing cells in 100uL of Glo Lysis Buffer (Promega, E26661) for 5 minutes on a rocker, then adding 100uL of CellTiter-Glo® (Promega, G7572) and rocking an additional 5 minutes. Data is presented as (HiBiT luminescence values / average titer luminescence value). Relative values are normalized to values acquired on the same plate. Luminescence was measured using a GloMax 96 Microplate Luminometer.

### Patient-derived iNeuron Meso scale discovery (MSD) assay

We adapted the GP MSD assay from Su et al^77^. We used an in-house rabbit polyclonal GP antibody as both the capture and detection antibody. The assay was validated using lysates from transfected HEK293 cells expressing GP70 GGG-NL-3xFlag and C9 BAC transgenic mice. For iNeuron MSD analysis, neural progenitor cells plated in 12 well plates were differentiated for 14 days as previously described^60^. After differentiation, iNeurons were treated with 5 µM of ASO and 1uL of Endoporter (GeneTools, OT-EP-PEG-1) at a final well volume of 500uL. On day 3 or 4 of ASO treatment, an additional 500uL of media was added to cells, diluting the effective ASO concentration to 2.5 µM. Cells were removed from the incubator on day 7 of ASO treatment and MSD assay was conducted as previously described^77^.

### Reconstitution of Human 48S Pre-Initiation Complex for CryoEM

The 48S complex was assembled *in vitro* by sequentially reconstituting the 43S pre-initiation complex and a cap-binding complex, followed by mixing the components in a final volume of 50 µL. Initiation factors were stored in factor buffer (20mM HEPES-NaOH, pH 7.5, 200mM NaCl, 10% glycerol, 5mM ß-Me) while the 40S ribosome was stored in reconstitution buffer (20 mM HEPES-KOH, pH 7.5, 70 mM KOAc, 4 mM Mg(OAc)2, 0.1 mM spermidine, 1 mM DTT). To assemble the 43S complex, 0.5 µM of the 40S subunits were incubated with 1.67 µM eIF3, 1.67 µM ternary complex (eIF2-GTP-tRNA_i_^Met^), 0.83 µM eIF1, eIF1A, eIF5, and 0.83 µM GMPPNP in reconstitution buffer. The reaction was incubated at 30 °C for 10 minutes in a final volume of 15 µL. The cap-binding complex was prepared by incubating 2.5 µM mRNA with 2.5 µM poly(A)-binding protein (PABP), 2.5 µM eIF4F complex (eIF4E, eIF4A, eIF4G), 5 µM additional eIF4A, and 1.25 mM ATP-γ-S (pH 7.0) in reconstitution buffer, in a final volume of 10 µL, at 30 °C for 10 minutes. The complete 48S complex was assembled by combining the 43S and cap-binding mixtures with 0.92 µM eIF4B, bringing the total reaction volume to 27 µL. This final mixture was incubated at 30 °C for an additional 10 minutes. After adding 20uL reconstitution buffer to a final concentration of 50uL prior to plunging, the final buffer conditions were (10.85mM HEPES-KOH, 4.323mM HEPES-NaOH, pH 7.5, 43.23mM NaCl, 37.975mM KOAc, 2.17mM Mg(OAc)2, 0.05mM spermidine, 2.2% glycerol, 1.08mM ß-Me, 0.54mM DTT).

### *In vitro* Reconstitution of Human 40S-ES9^S^ targeting ASO complex

The 40S-ES9^S^ targeting ASO complex was assembled *in vitro* by first refolding the ES9^S^ targeting ASO in 1X oligo refolding buffer as described above. Next, the complex was assembled at 20uL with 1 µM eIF1 and eIF1A, 2 µM ASO and 150nM 40S purified from HEK293T cells. Reconstitution buffer (20 mM HEPES-KOH pH 7.5, 70mM KAc, 4 mM MgAc, 0.1mM spermidine and 1mM DTT) was used to dilute the sample to 20uL. The complex was then incubated at 30 °C for 10 minutes and placed directly on ice prior to plunging.

### Purification of Native Human Initiation Complexes from Lysate

The sample was prepared by MNase treating commercially available HeLa Cell lysate (IPRACell sku: CC-01-40-50) for 5 minutes at 25C shaking 300 rpm with MNase (NEB M0247S) at a final concentration of 0.015 units/ul. The reaction was quenched with EGTA at a final concentration of 3 mM. Samples were prepared by taking 1129 ul of a MNase treated lysate and adding 160 ul of 10X Translation Buffer (160 mM HEPES-KOH pH 7.6, 200 mM Creatine Phosphate, 1 mM Spermidine, 500 mM Potassium acetate, 25 mM magnesium acetate, and 1 ug/ul of creatine phosphate kinase), 5 ul of RNaseIN Plus (Promega N2611), 32 ul of complete amino acid mix (Promega L4461), ATP (Thermofisher Scientific 50591512) at a final concentration of 1 mM, GMP-PNP (Sigma G0635) and magnesium acetate mixture at a final concentration of 5 mM each, and an in-vitro transcribed mRNA containing the beginning of intron 1 of the gene C9orf72 followed by 70 repeats of GGGGCC (G4C2) at a final concentration of 200 nM. The reaction was incubated for 20 minutes at 37C shaking 300 rpm before being loaded onto a 10-30% sucrose gradient (20 mM HEPES KOH pH 7.6, 100 mM Potassium acetate, 3 mM Magnesium acetate, 1 mM DTT and 1 protease inhibitor tablet (Roche 11836170001)). The sample was centrifuged for 5 hours at 40,000 rpm using a SW41 Ti rotor. Fractions were taken to collect the main 40S peak and avoid 60S contamination. Fractions were pooled and centrifuged for 2 hours and 30 minutes at 108,000 rpm using a TLA 120.2 rotor. Supernatant was removed and the pellet was resuspended in 48S Complex Buffer (20 mM HEPES-KOH pH 7.6, 70 mM potassium acetate, 4 mM magnesium acetate, 1 mM DTT, 0.1 mM spermidine).

### CryoEM Grid Preparation and Data Collection

The late-stage initiation complexes were crosslinked with 1.5 mM bis(sulfosuccinimidyl)suberate (BS3) (Thermofisher Scientific A39266) on ice for 45 minutes and diluted to 150 nM with reconstitution buffer for grid preparation. 3 µL of sample was applied to UltrAuFoil R1.2/1.3 300-mesh gold grids coated with graphene oxide (Sigma, 763705) as previously described^41,112^. Grids were prepared using a Vitrobot (FEI) at 4 °C and 100% humidity, blotted for 8 or 9 seconds at −15 blotting force, and plunge-frozen in liquid ethane at 93 K. CryoEM data were acquired using a Titan Krios G4i microscope (ThermoFisher) equipped with a K3 direct electron detector and BioQuantum imaging filter (Gatan) at 105,000 magnification, pixel sizes 0.8487 Å or 0.8320 Å (post-Nov 2023). SerialEM was used with multi-shot mode (2 shots per hole), and defocus values ranged from −1.2 to −3.0 µm. Electron doses per frame were: 1.48 e^−^/Å^2^ (Data 1), 1.45 e^−^/Å^2^ (Data 2), and 1.317 e^−^/Å^2^ (Data 3).

The ASO-40S complexes were not crosslinked and were applied to QUANTIFOIL® Holey Carbon (ultra thin +2nmA). Grids were prepared using a Vitrobot (FEI) at 4 °C and 100% humidity, blotted for 7 or 8 seconds at −15 blotting force, and plunge-frozen in liquid ethane at 93 K. Acquisition microscope, detector and analysis software were the same as the late-stage initiation complexes.

For native complexes, sample was crosslinked on ice for 45 minutes using a final concentration of 1.5 mM BS3 (Thermofisher Scientific A39266). Following crosslinking, the sample was diluted to 200 nM of ribosomes using the 48S complex buffer. 3 ul of 200 nM 48S complex was applied to glow-discharged QUANTIFOIL R1.2/1.3 gold 300 mesh with 2 nm carbon grids. CryoEM grids were prepared using a FEI Vitrobot Mark IV at 4C and 100% humidity, with blot force of 7 and blot time of 6 seconds. The grids were plunged into liquid ethane at 93 K in a precision cryostat system.

Two independent data sets were collected on a Titan Krios G2 microscope (ThermoFisher) equipped with Gatan K III direct electron detector (FEI) at a magnification of 105,000x and at pixel sizes of 0.834 Åpix-1. SerialEM was used with multi-shot mode (2 shots per hole) with defocus values ranging from −1.2 to −3.0 µm and electron doses per frame of 0.677 e^−^/Å^2^ (Data 1) and 0.744 e^−^/Å^2^ (Data 2).

### Image Processing

Motion correction and dose-weighting were performed using RELION 5^44,107,108^ with a 5×7 patch alignment. Contrast transfer function (CTF) estimation was carried out using CTFFIND4^115^. Automated particle picking was seeded using reference-based auto picking. A total of 250 classes were generated via 2D classification, and selected classes were further refined using 3D auto-refinement. Focused 3D classification was conducted early in processing for both the *in vitro* and native structures to identify ribosome-containing particles for further classification. These particles were then refined using CTF refinement and Bayesian polishing in Relion^116^. After polishing, 3D classification on ribosomal subdomains (e.g., head region, ternary complex, ES9^S^) was performed (see Extended data Fig. 2 and 5). For focused 3D classification, a 3 Å molmap was created of the region of interest from a model ridgid body fit into the region in ChimeraX (6zmw)^42^. The molmap was resampled to the refined map to be sorted and a mask was created around the region of interest. For differentiating scanning vs initiated particles in our native structure, the particles were first classified by head position, then were classified for either ternary complex into eIF3 or ES9^S^-G_4_C_2_ followed by the ternary complex into eIF3. Final resolutions were estimated based on the gold-standard Fourier Shell Correlation (FSC) at the 0.143 criterion.

### Model Building and Refinement

Initial models were derived from previously published human 48S structures (PDB: 6zmw, 8PJ2)^42,44^. A kissing-loop interaction at ES9^S^ was modeled using the TAR-R06 kissing-loop crystal structure^117^, with sequence adaptations based on ES9 and repeat region alignments. Atomic model building was conducted in Coot^118^ followed by real-space refinement using Phenix^119^. Figures were made using ChimeraX^120^.

### Figures and text

All structural figures were generated using UCSF ChimeraX^120^. AI-based language tools, including Grammarly and ChatGPT, were used to correct typographical errors and improve clarity by reducing word count.

## Data availability

Atomic coordinates have been deposited in the Protein Data Bank under accession codes 39RN (in vitro RAN late-stage initiation complex), 39RL (in vitro ES9^S^ region), and 39RT (near-native IC). Corresponding cryoEM density maps are available in the Electron Microscopy Data Bank under accession numbers EMD-79722 (RAN late-stage initiation complex), EMD-79720 (ES9^S^ region), and EMDB-79727 (near-native IC). All relevant source data are provided with this paper.

## ACKNOWLEDGEMENTS

We thank Caroline Hsieh and the Sami Barmada lab for their recommendations and expertise in iNeuron experimental conditions, the LSI CryoEM Facility for help with electron microscopy data collection, the Petrucelli lab for assistance with setting up the GP-MSD assay, and Christopher Lilienthal and Bradford Battey for assistance with scientific computing. We thank Adrian Isaacs’s lab at UCL for their gift of the C9orf72 patient-derived Poly-GA Hi-BiT iPSC cell line^60^.

## Author contributions

W. D., S. L., M.S, C.S.F and J.B.Q purified eIFs. R.K., and J.B.Q purified mRNA, assembled the *in vitro* reconstituted late-stage initiation complexes. B.S. performed native complex purification. R.K, W.D, B.S. and J.B.Q performed cryoEM data collection, processing, atomic model building, and refinement. R. K., C.W., I.P. and B.S. performed *in vitro* translation. E.L. and I.P. performed iPSC and iNeuron differentiation and management. R.K. and I.P. performed experiments on patient iNeurons. I.P. and C.W. performed mRNA transfection experiments. R.K. designed ES9^S^-targeting ASOs. A.K. conducted the MSD assays. R.K., B.S. and J.B.Q. interpreted the structures. R.K, P.K.T. and J.B.Q wrote the manuscript with contributions from all authors. P.K.T and J.B.Q. conceived the project and obtained the funding to support the project.

## EXTENDED DATA

**Extended Data Figure 1:**
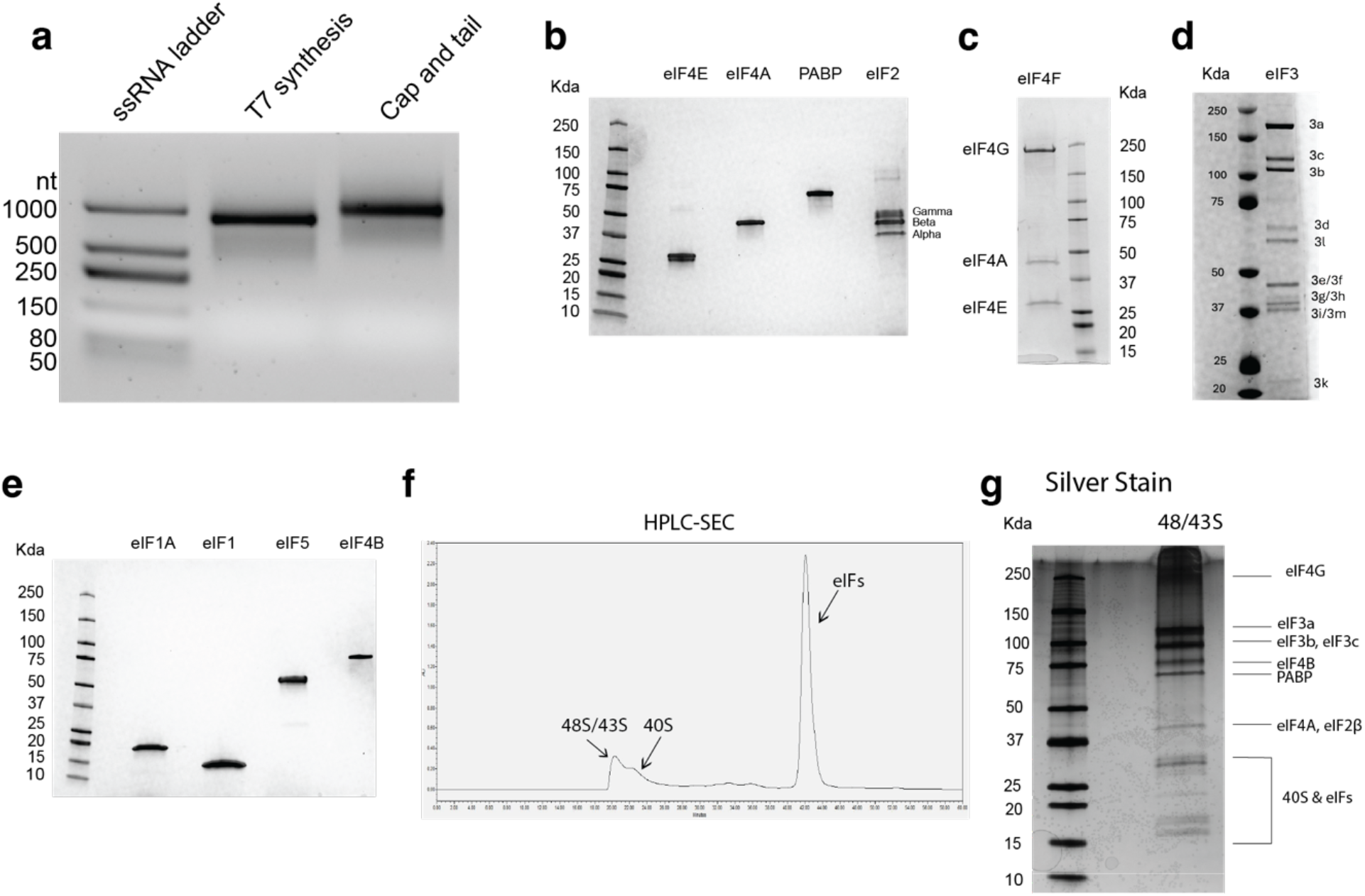
Purification of RNA and eukaryotic initiation factors. **a**, Purified RNA visualized on a 1% agarose gel stained with 1× GelGreen. RNA is shown before (middle) and after (right) enzymatic capping and polyadenylation. **b**–**e**, Coomassie-stained SDS-PAGE gels of purified initiation factors used for assembly of the *in-vitro* reconstituted RAN 48S complex: **b**, ternary complex (eIF2–GTP–Met-tRNAi^Met^), **c**, eIF4F cap-binding complex, **d**, eIF3, and **e**, additional factors including eIF1, eIF1A, eIF5, and eIF4B. **f**, HPLC size exclusion trace of the fully assembled complex. **g**, Silver stain of the assembled 48/43S peak obtained from the HPLC.

**Extended Data Figure 2:**
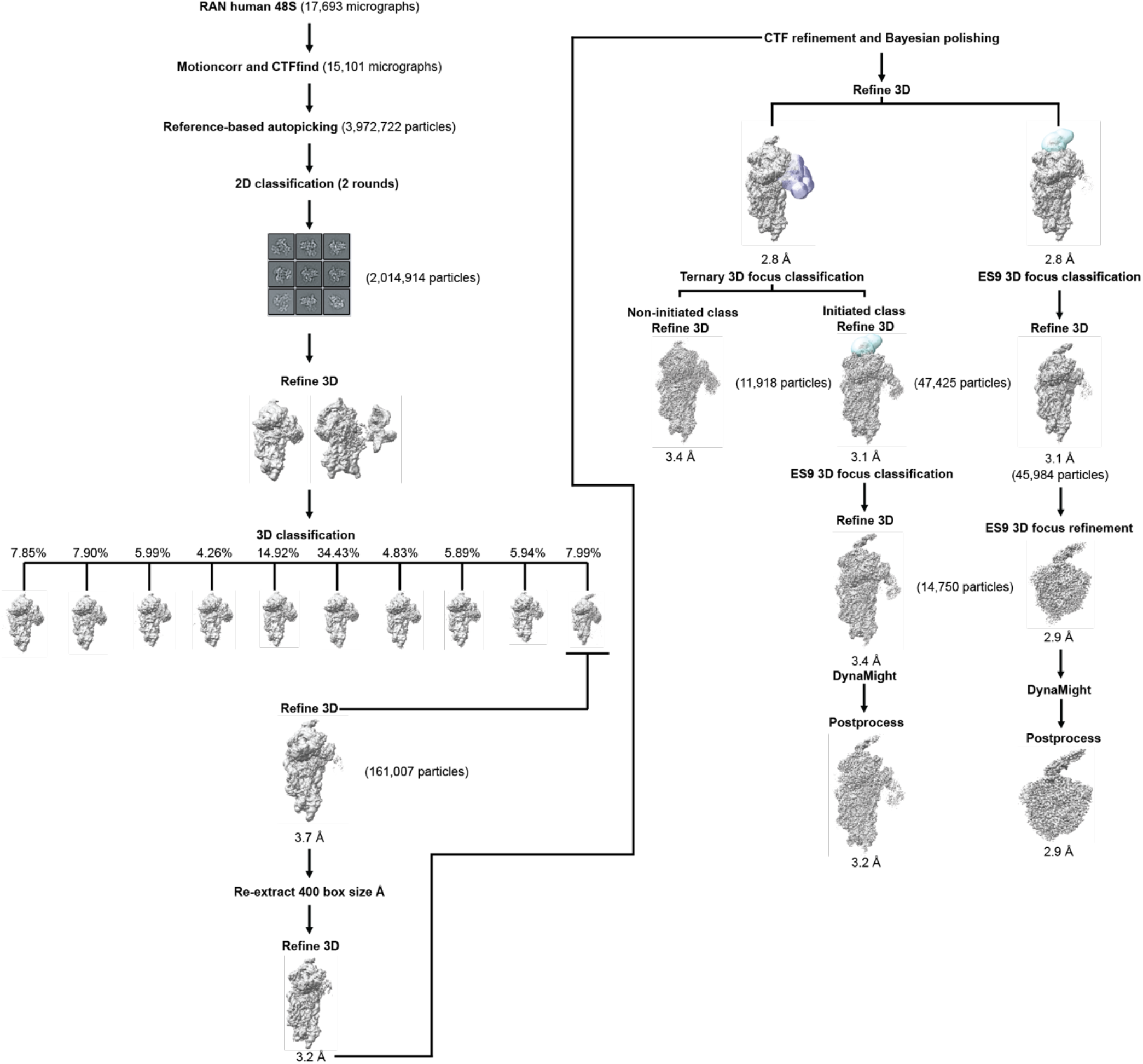
CryoEM data processing workflow for the *in vitro* reconstituted RAN 48S initiation complex. Schematic of cryoEM data processing for the *in vitro* reconstituted RAN 48S complex. Final reconstruction of the overall complex was generated from 14,750 particles selected after 3D focus classification. The ES9^S^-focused 3D refinement used 28.6% of classified particles. The overall map was derived from 9.2% of particles after sequential classification.

**Extended Data Figure 3:**
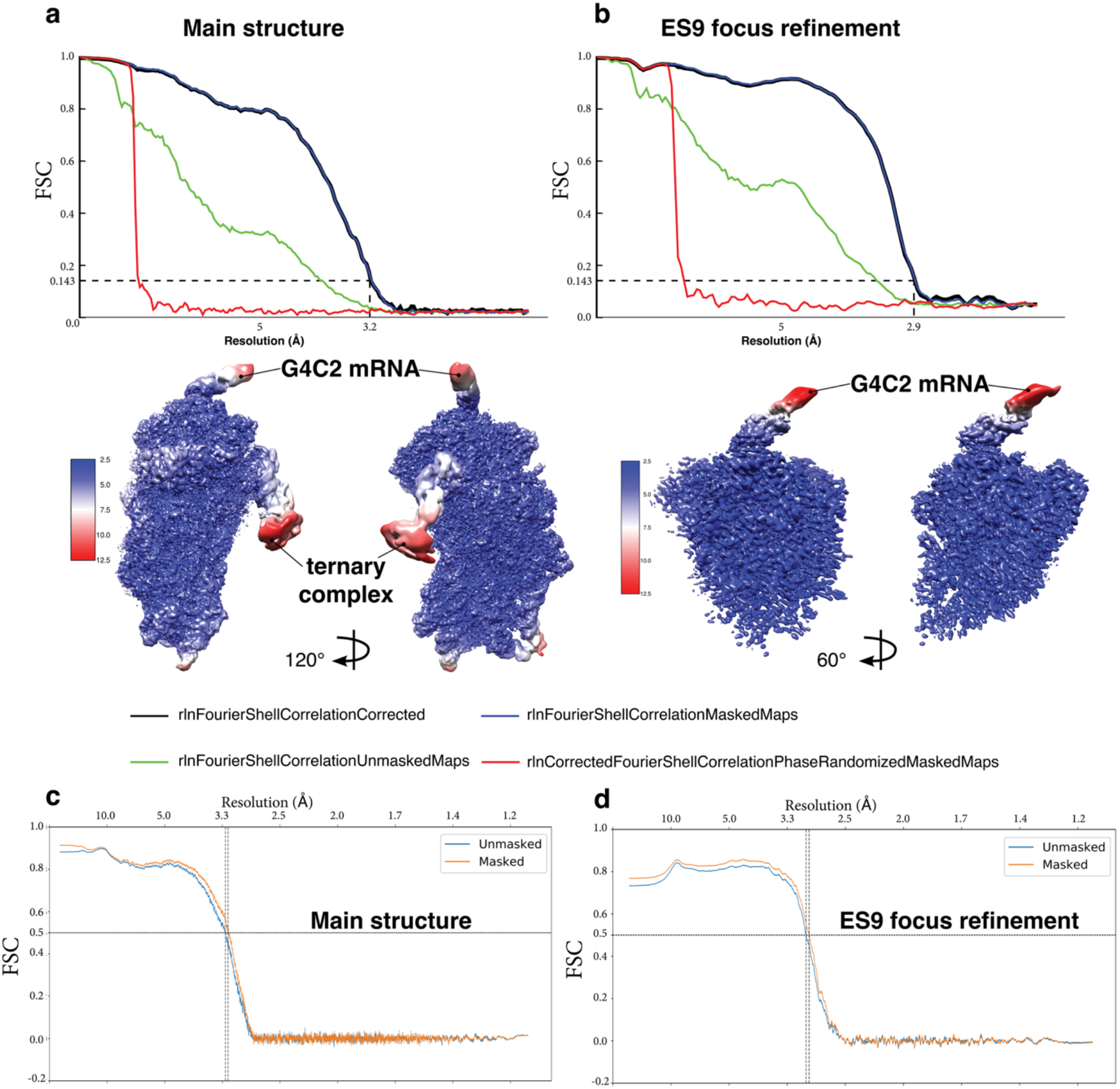
Local resolution and model validation for the *in-vitro* reconstituted RAN 48S initiation complex. **a**–**b**, Local resolution maps and Fourier shell correlation (FSC) plots for main structure and ES9^S^ focus refinement. Local resolution maps for both structures presented underneath the curves in two orientations. **c-d**, Model-to-map FSC curves calculated using Mtriage in Phenix, indicating resolution at FSC = 0.5.

**Extended Data Figure 4:**
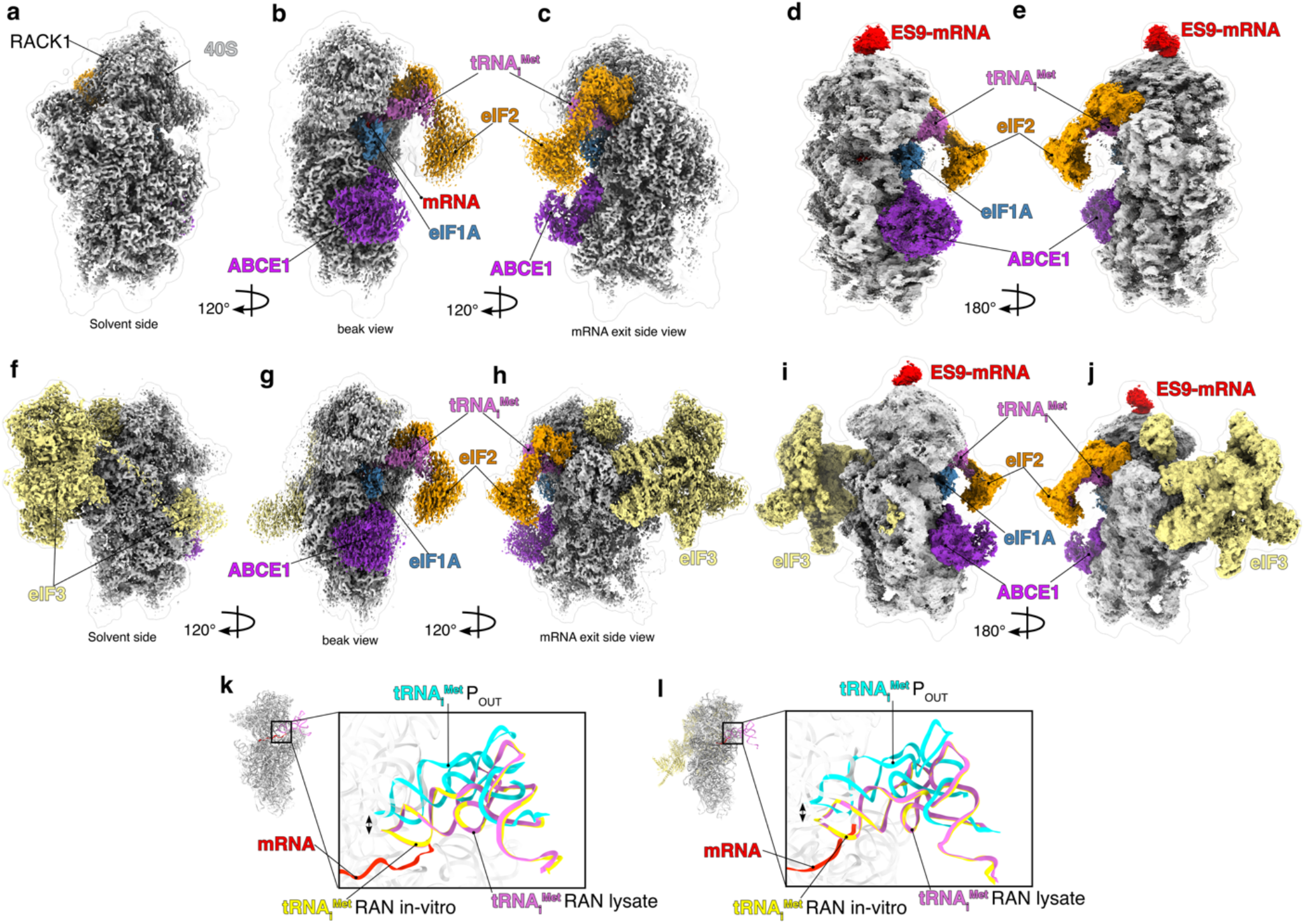
Structure of the RAN 48S initiation complex purified from human cell lysate. **a-c**, Overview of the electron density map of the 3.54 Å RAN initiation structure presented without the presence of eIF3. eIF2, tRNA_i_^Met^, ABCE1, eIF1A and mRNA are highlighted in the structure. **d-e**, Overview of the electron density map without eIF3 and with the ES9^S^-G_4_C_2_ kissing loop interaction. **f-h**, Overview of the electron density map of the 4.76 Å RAN initiation structure with eIF3 present. **i-j**, Overview of the electron density map with eIF3 and with the ES9^S^-G_4_C_2_ kissing loop interaction. **k-l**, Superposition of the tRNA_i_^Met^ position between a canonical scanning 48S complex (cyan P_out_, PDB: 8PJ1) and the *in-vitro* reconstituted RAN 48S initiation complex (yellow RAN *in-vitro*, PDB: 9PG5) reveals the 48S RAN translation initiation complex is in a closed conformation (orchid). Superpositions shown for (**k**) the lysate structure without eIF3 and (**l**) with eIF3.

**Extended Data Figure 5:**
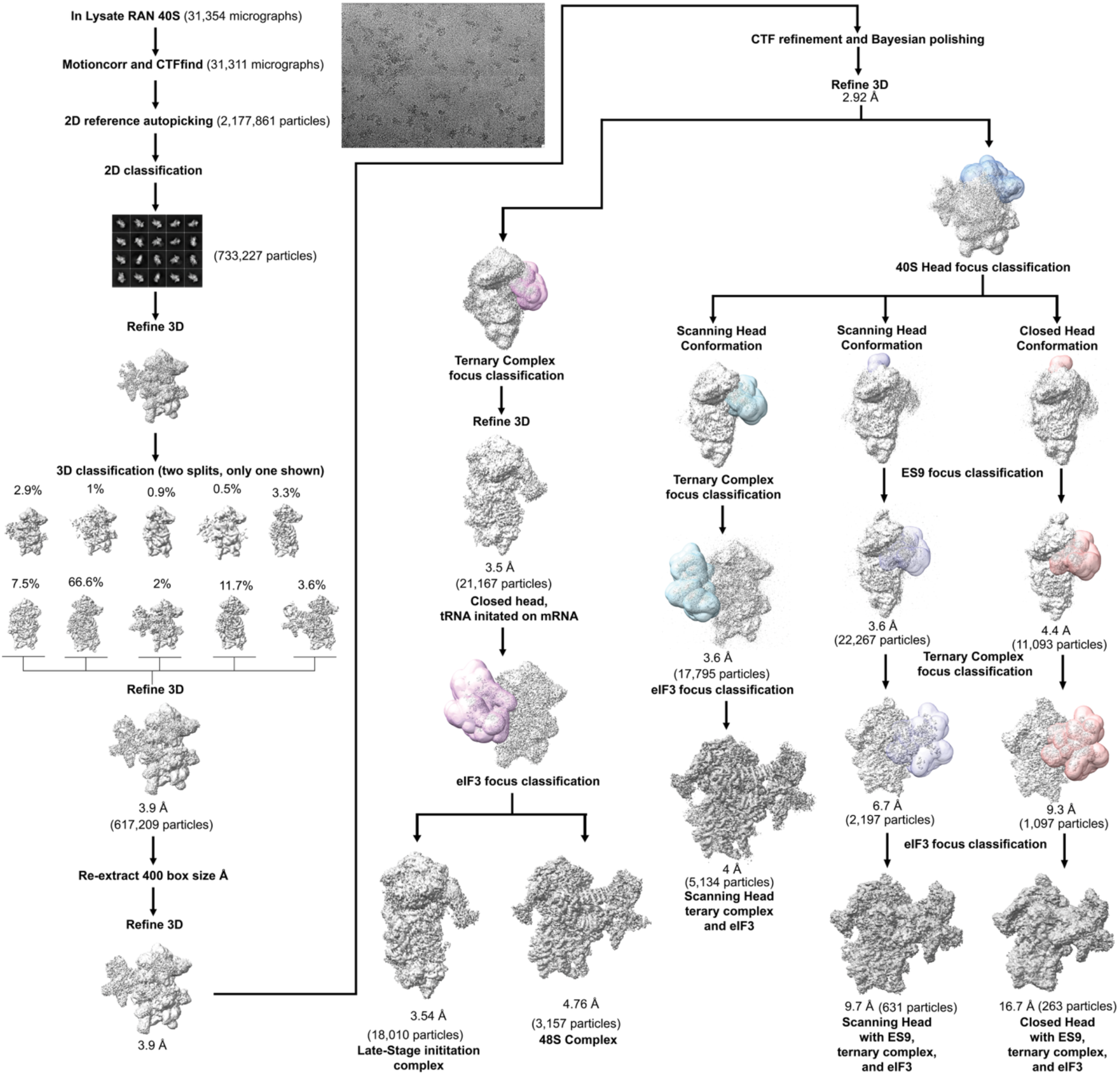
CryoEM data processing workflow for the RAN 48S initiation complex purified from human cell lysate. Schematic of cryoEM data processing for the RAN 48S complex purified from HeLa lysate. Initiated complexes without the ES9^S^-G_4_C_2_ kissing loop were resolved from 3D classification of the ternary complex (2.9% of particles) followed by eIF3 (0.5% of particles). Reconstructions of initiated complexes with the ES9^S^-G_4_C_2_ kissing loop were resolved from 3D classification of the head, followed by focus classification on ES9^S^ (1.8%), the ternary complex (0.2%) and eIF3 (0.04%). Scanning complexes without the ES9^S^-G_4_C_2_ kissing loop were resolved from 3D classification of the head, followed by the ternary complex (2.9%) and eIF3 (0.8%). Reconstructions of scanning complexes with the ES9^S^-G_4_C_2_ kissing loop were resolved from 3D classification of the head, followed by focus classification on ES9^S^ (3.6%), the ternary complex (0.4%) and eIF3 (0.1%).

**Extended Data Figure 6:**
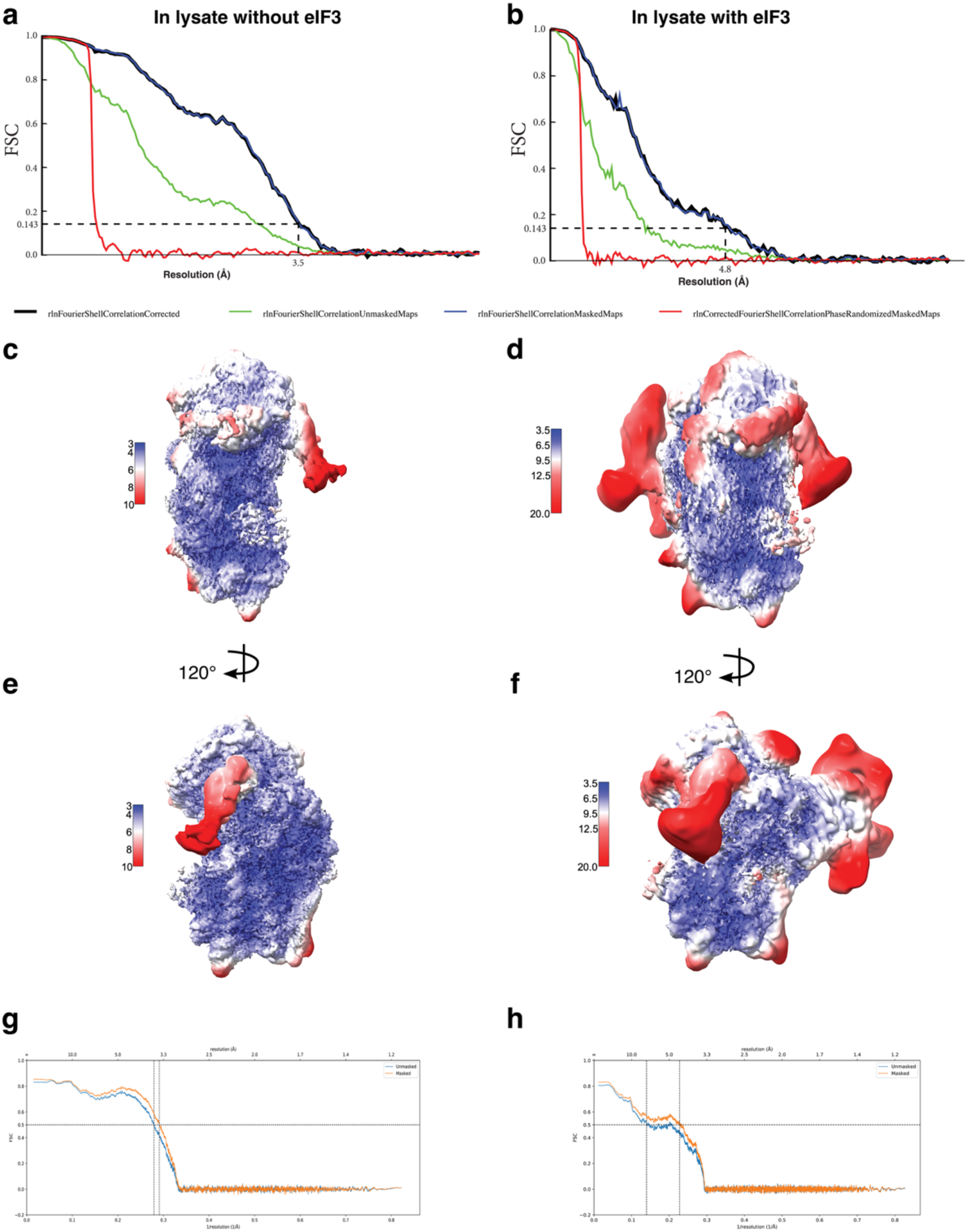
Local resolution and model validation for the RAN 48S initiation complex purified from human cell lysate. **a**–**b**, Local resolution maps and Fourier shell correlation (FSC) plots for main structure and ES9^S^ focus refinement for the in-lysate RAN 48S initiation complexes (**a**) without eIF3 and (**b**) with eIF3. **c-f**, Local resolution maps for both structures presented in two orientations. **g-h**, Model-to-map FSC curves calculated using Mtriage in Phenix, indicating resolution at FSC = 0.5.

**Extended Data Figure 7:**
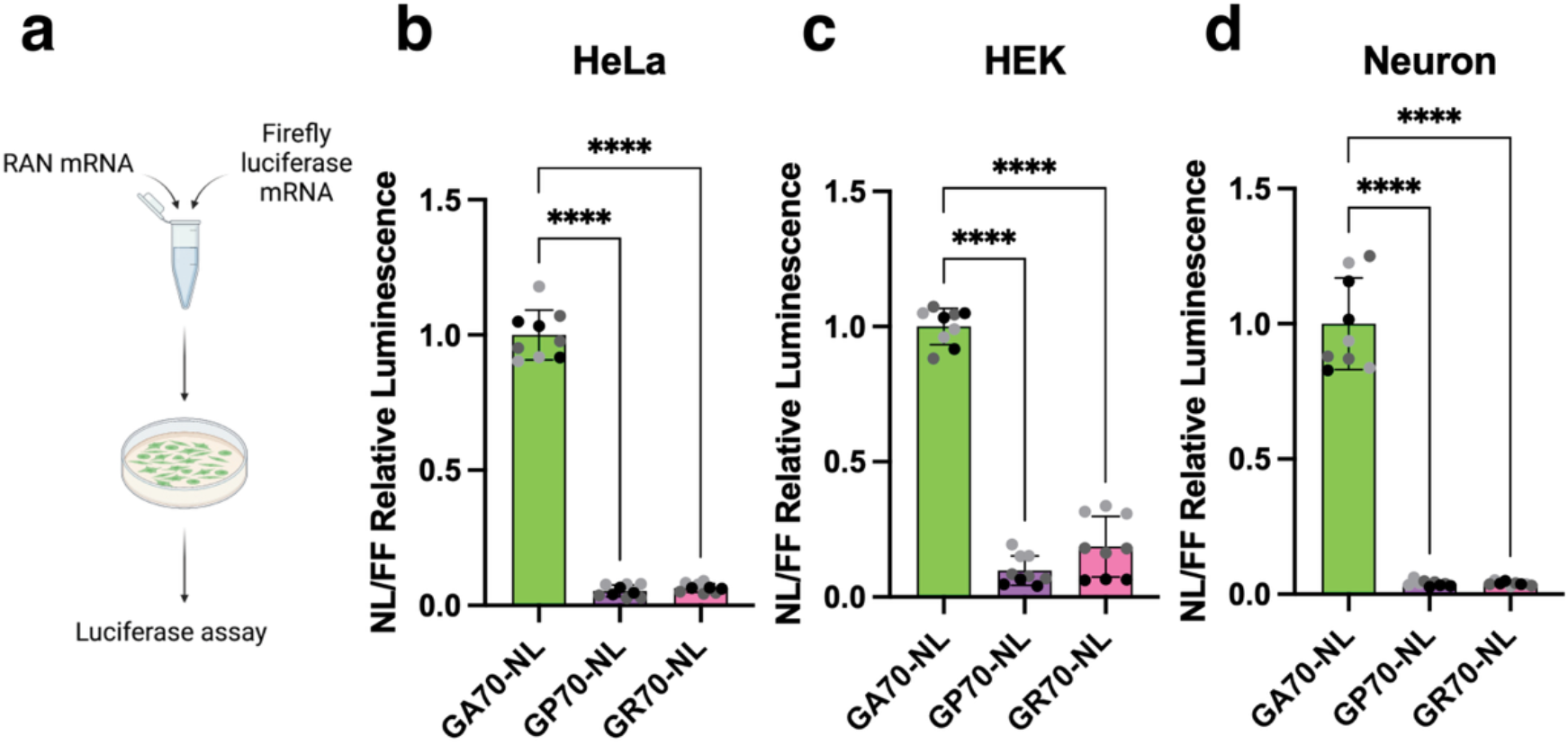
RAN translation produces DPRs most efficiently in the GA frame across multiple cell types. **a**, Schematic depicting mRNA transfection of RAN translation nanoluciferase reporters mixed with a firefly luciferase mRNA transfection control. **b-d**, GA70-NL reporters translate more efficiently than GP70-NL or GR70-NL in (**b**) HeLa cells (**c**) HEK cells and (**d**) rat hippocampal neurons.

**Extended Data Figure 8:**
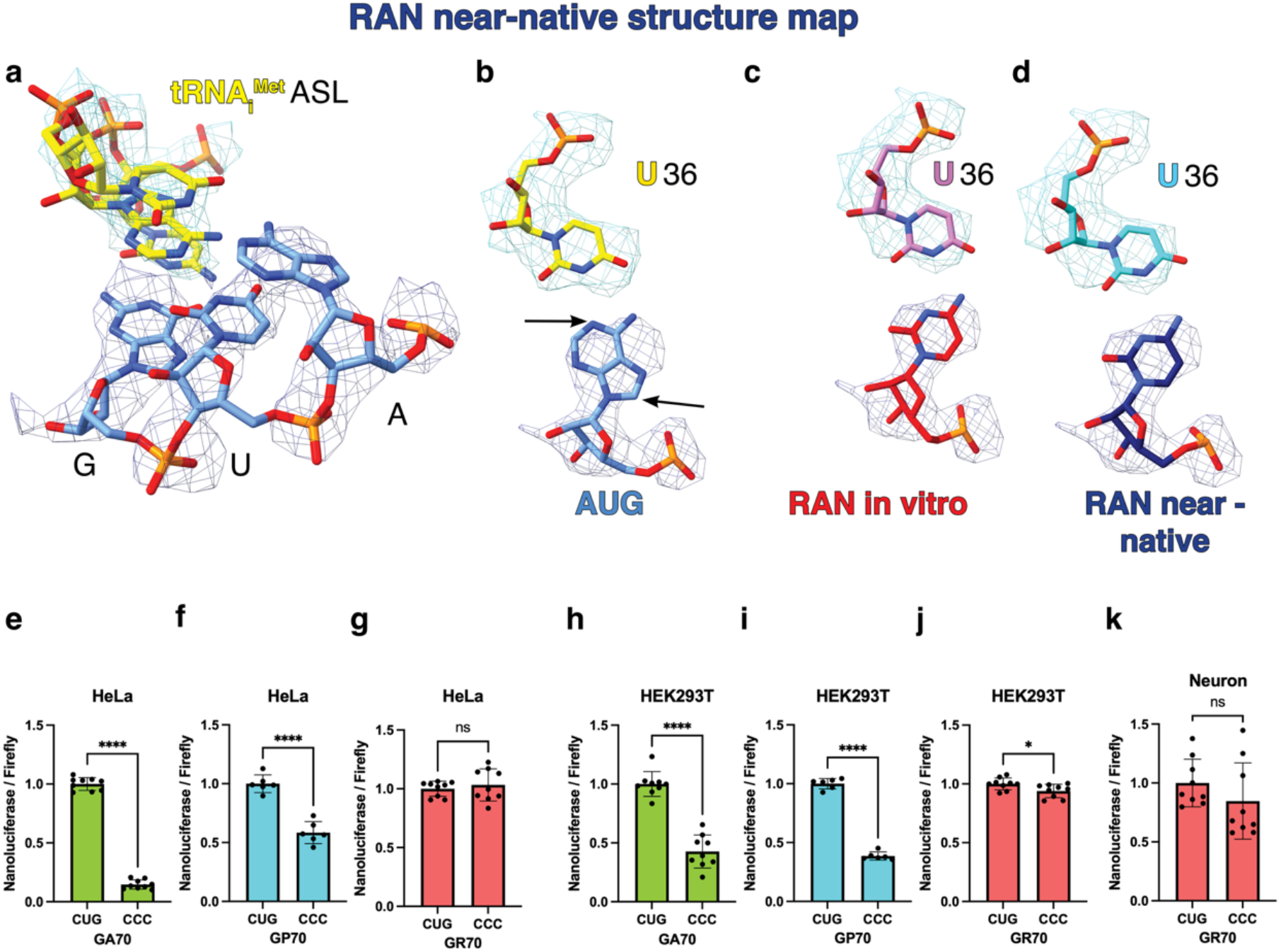
C9orf72 RAN translation initiates at a near-cognate CUG codon *in* vitro and in native conditions. **a-d**, Our near-native RAN initiation map fit with (**a-b**) the canonical AUG model, (**c**) the *in vitro* RAN initiation structure model and (**d**) the near-native RAN initiation structure model. The N1 and C8 atoms on the adenine do not fit in the RAN initiation density, indicating that this initiation base cannot be an adenine. **e-k**, Transfection of C9orf72 mRNA reporters in the GA, GP and GR frames with a competent CUG vs a mutated CCC codon in (**e-g**) HeLa cells, (**h-j**) HEK293 cells or (**k**) rat hippocampal neurons. Our data supports that mutation of the CUG near-cognate significantly reduces GA and GP production but does not significantly reduce RAN translation in the GR frame in HeLa cells or neurons and supports a minor reduction in HEK293 cells. Error bars represent the mean ± SD (n=9) obtained from three independent replicates. For GP, (**f, i)** n=6. Statistical significance: *p < 0.05, ****p <0.0001.

**Extended Data Figure 9:**
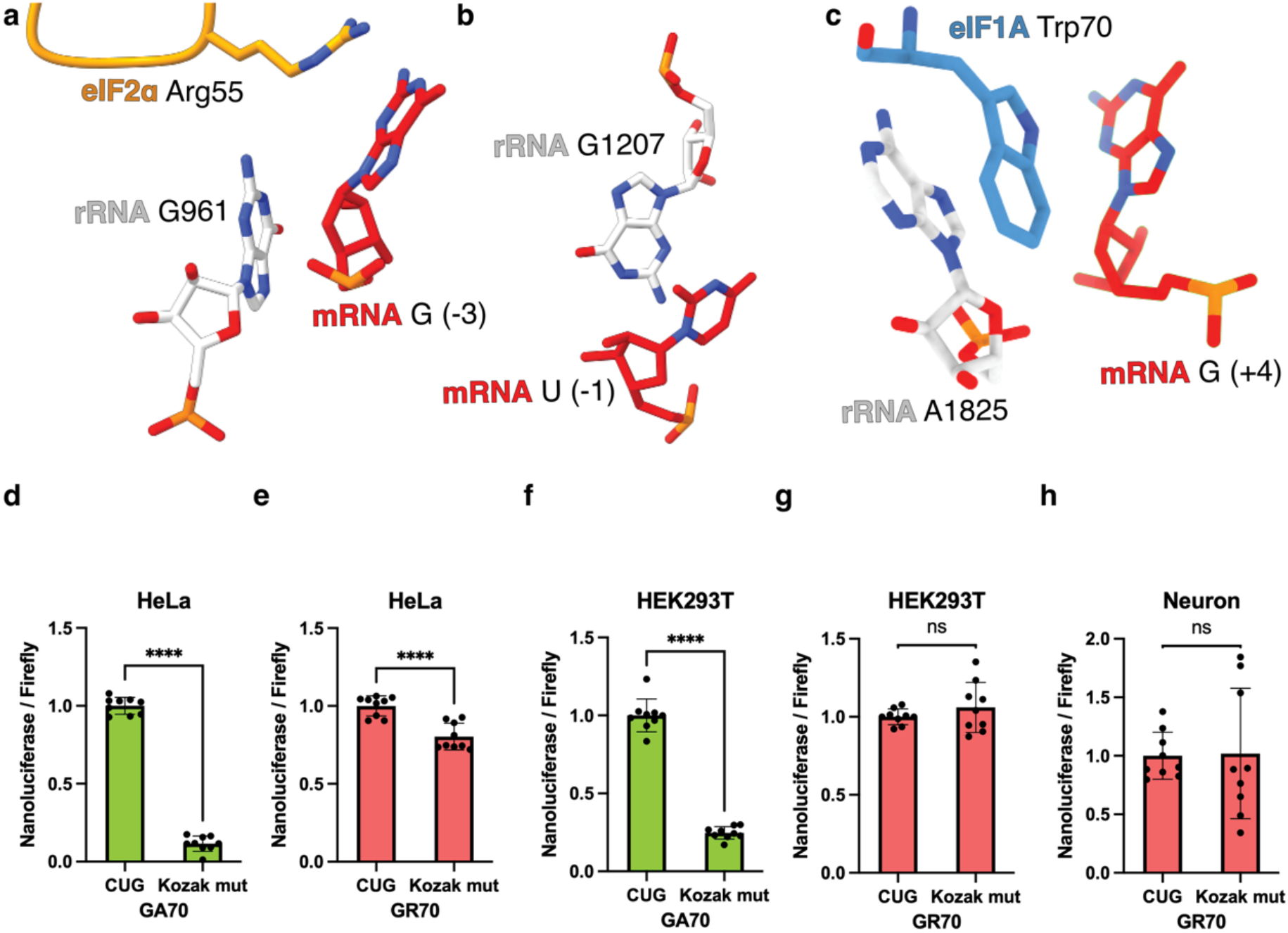
The Kozak sequence surrounding the CUG near cognate codon is structurally and functionally important for C9orf72 RAN translation. **a-c**, Structural model illustrating stabilization of the Kozak context in the RAN 48S initiation complex through interactions with components of the mRNA binding channel: **a**, eIF2α Arg55 interacts with the nucleotide at position –3 and with rRNA G691; **b**, rRNA G1207 contacts the nucleotide at position –1; **c**, rRNA A1825 and eIF1A Trp70 interact with the nucleotide at position +4. **d-h**, transfection of C9orf72 mRNA reporters in the GA and GR frames with a competent CUG and Kozak sequence vs a mutated Kozak sequence (−3G & +4G to Cs) in (**d-e**) HeLa cells, (**f-g**) HEK293 cells or (**h**) rat hippocampal neurons. Our data supports that mutation of the Kozak sequence significantly reduces GA production but does not significantly reduce RAN translation in the GR frame in HEK293 cells or neurons and supports a minor reduction in HeLa cells. Error bars represent the mean ± SD (n=9) obtained from three independent replicates. Statistical significance: ****p <0.0001.

**Extended Data Figure 10:**
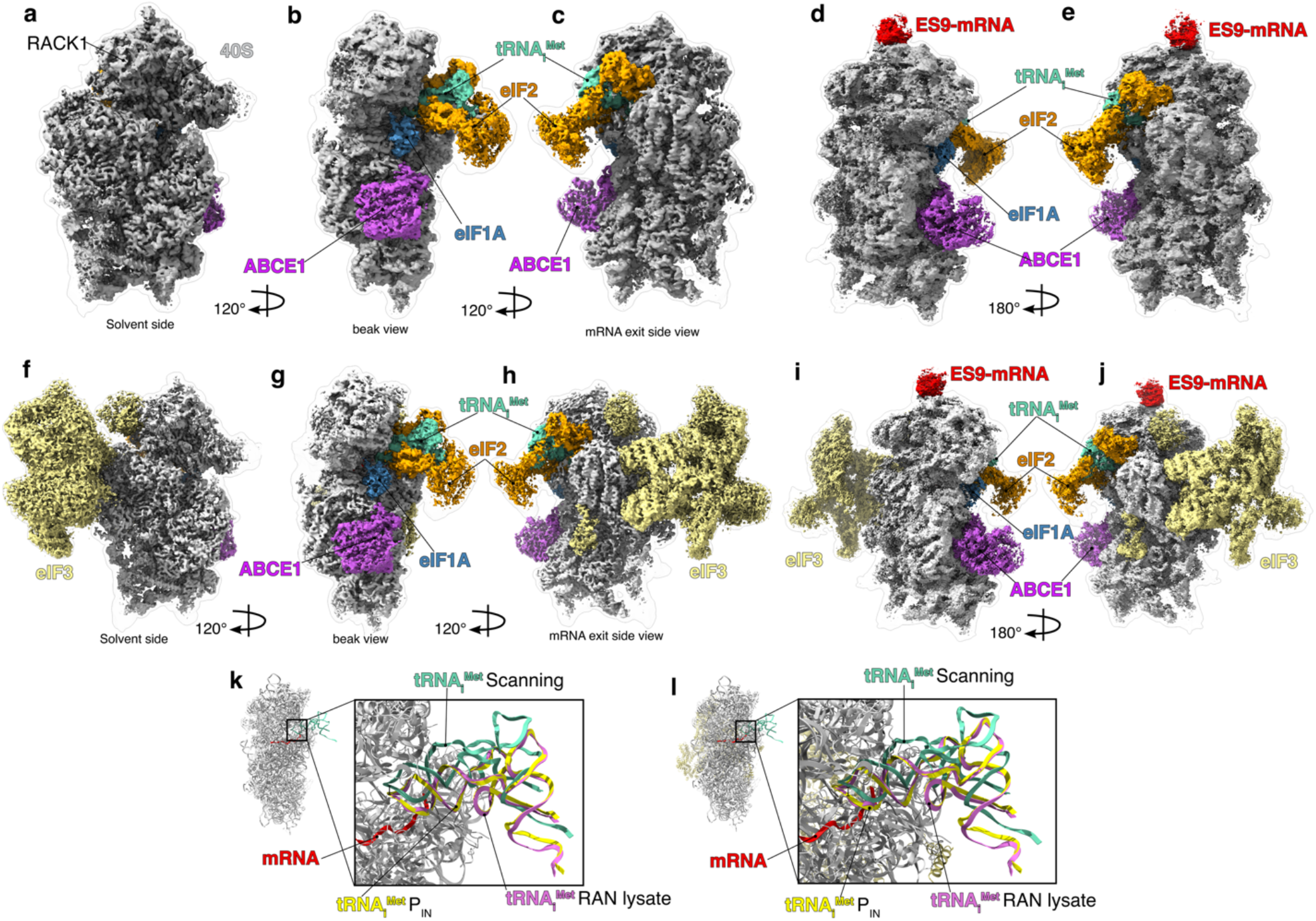
Near-native RAN 48S scanning complexes identified during data processing. **a-c**, Overview of the electron density map of the 3.54 Å RAN scanning structure presented without the presence of eIF3. eIF2, tRNA_i_^Met^, ABCE1, and eIF1A are highlighted in the structure. **d-e**, Overview of the electron density map without eIF3 and with the ES9^S^-G_4_C_2_ kissing loop interaction. **f-h**, Overview of the electron density map of the 4 Å RAN scanning structure with eIF3 present. **i-j**, Overview of the electron density map with eIF3 and with the ES9^S^-G_4_C_2_ kissing loop interaction. **k-l**, Superposition of the tRNA_i_^Met^ position between a canonical initiation 48S complex (yellow P_in_, PDB: 8PJ2), the near-native RAN 48S initiation complex (orchid RAN lysate) and our near-native RAN scanning structure (seafoam green scanning) reveals the 48S RAN translation scanning complex is in an open conformation (seafoam green). Superpositions shown for (**k**) the scanning structure without eIF3 and (**l**) with eIF3.

**Extended Data Figure 11:**
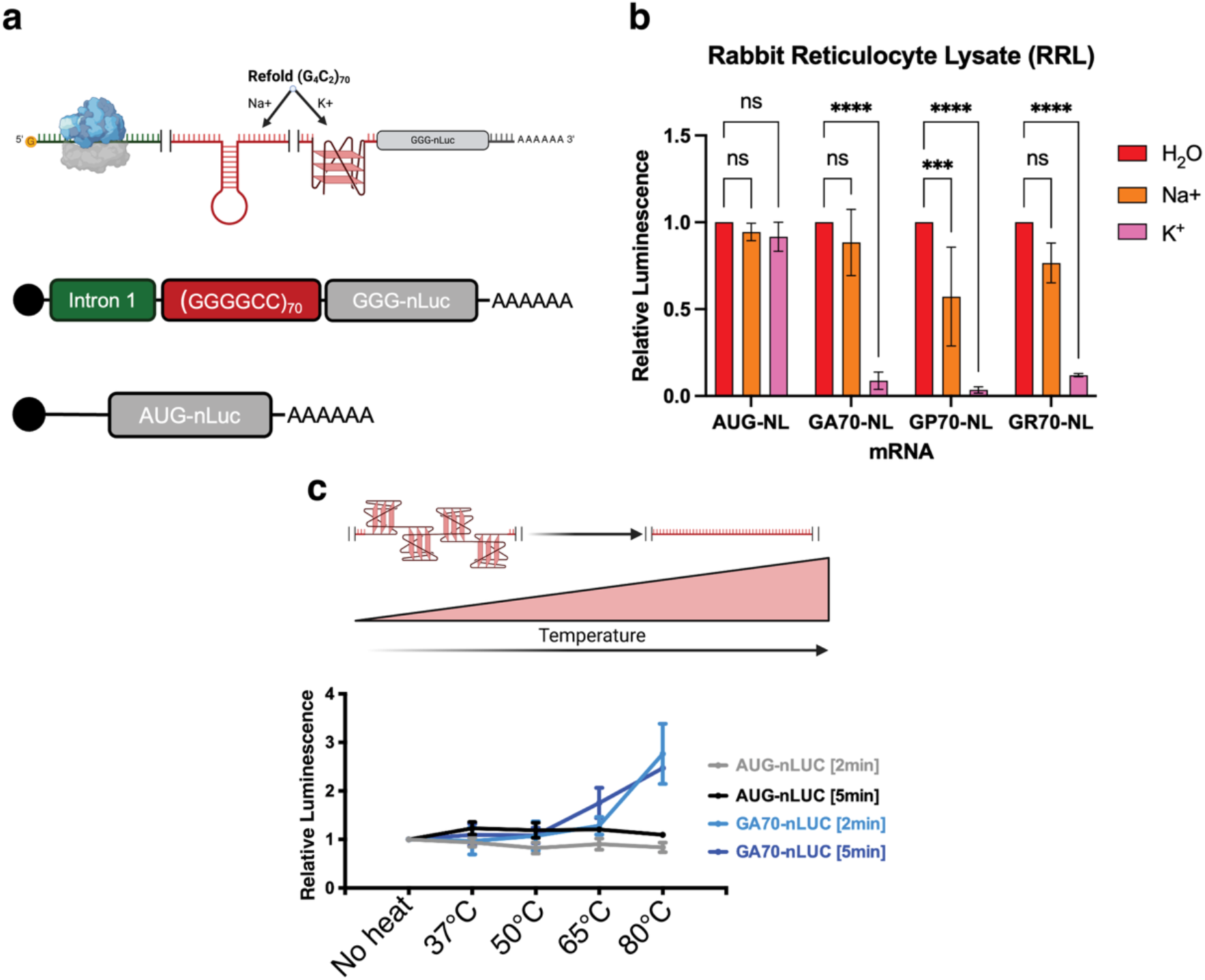
G-quadruplex formation inhibits RAN translation *in vitro*. **a**, Model of refolding of C9orf72 (G_4_C_2_)_70_ RNA in a hairpin or quadruplex formation. **b**, *In vitro* translation of AUG-NLuc RNA and C9orf72 (G_4_C_2_)_70_-NL RNA in the GA, GP and GR reading frames in rabbit reticulocyte lysate (RRL) after folding in K^+^ or Na^+^-containing buffers. K^+^ promotes G-quadruplex formation, which significantly inhibits RAN translation *in-vitro*. **c**, Heat denaturation of (G_4_C_2_)_70_-NL GA70 RNA folded in K^+^ buffer restores translation. RNAs were heated at indicated temperatures for 2 or 5 minutes prior to translation.

**Extended Data Figure 12:**
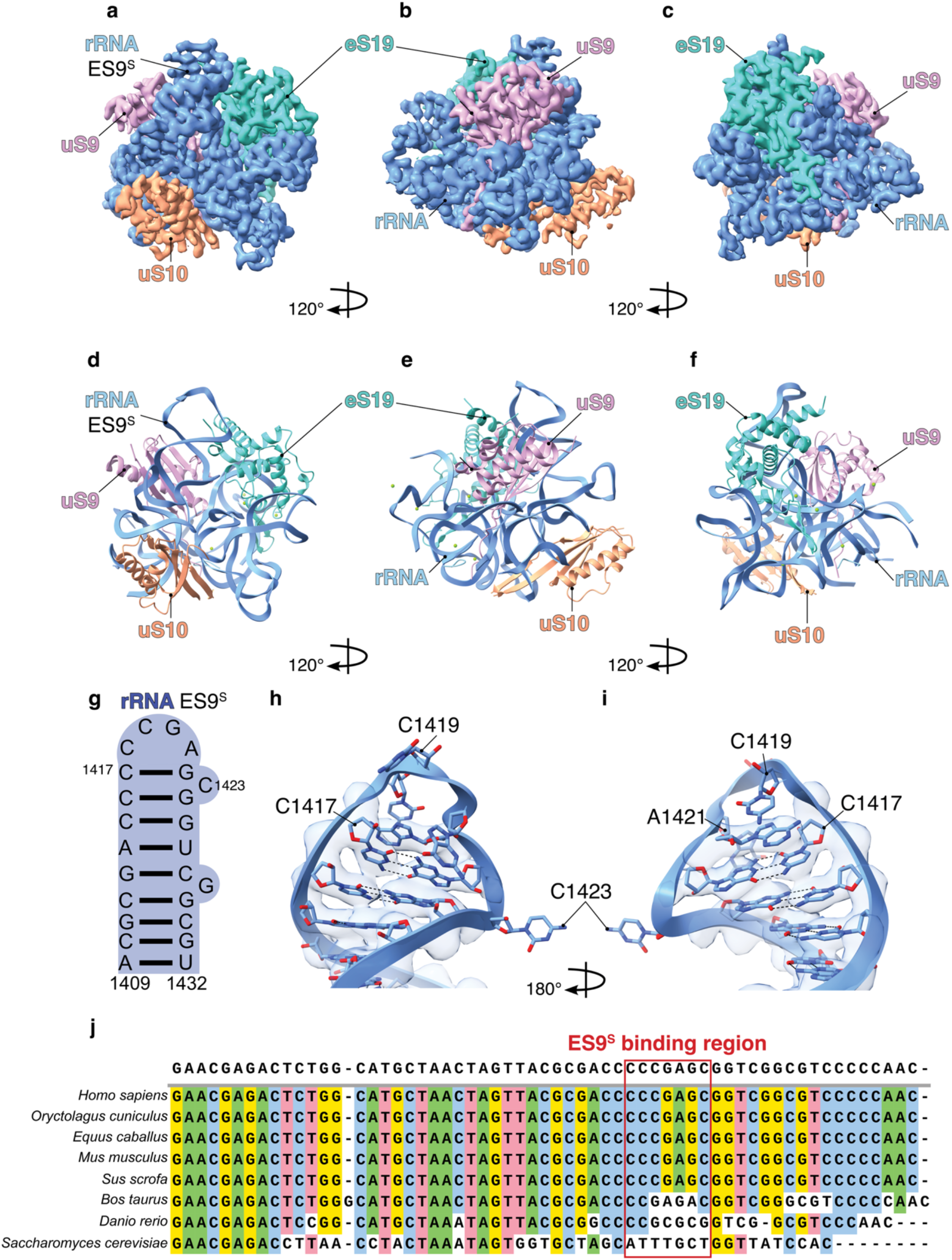
Structure and sequence conservation of 18S rRNA expansion segment 9 (ES9^S^). **a**–**c**, Electron density maps of the 40S ES9^S^ region in three orientations. rRNA (cornflower blue), uS9 (pink), uS10 (orange), and eS19 (turquoise); maps filtered with Gaussian filter (σ = 1). **d**–**f**, Molecular model fitted into the density. **g**, Schematic of the ES9S secondary structure. **h**–**i**, Isolated views of the ES9S turn region, highlighting canonical Watson–Crick base pairing between C1417 and G1422 and four unpaired nucleotides spanning the loop. **j**, Multiple sequence alignment showing evolutionary conservation of the G_4_C_2_-binding region in mammals but not in *Saccharomyces cerevisiae*.

**Extended Data Figure 13:**
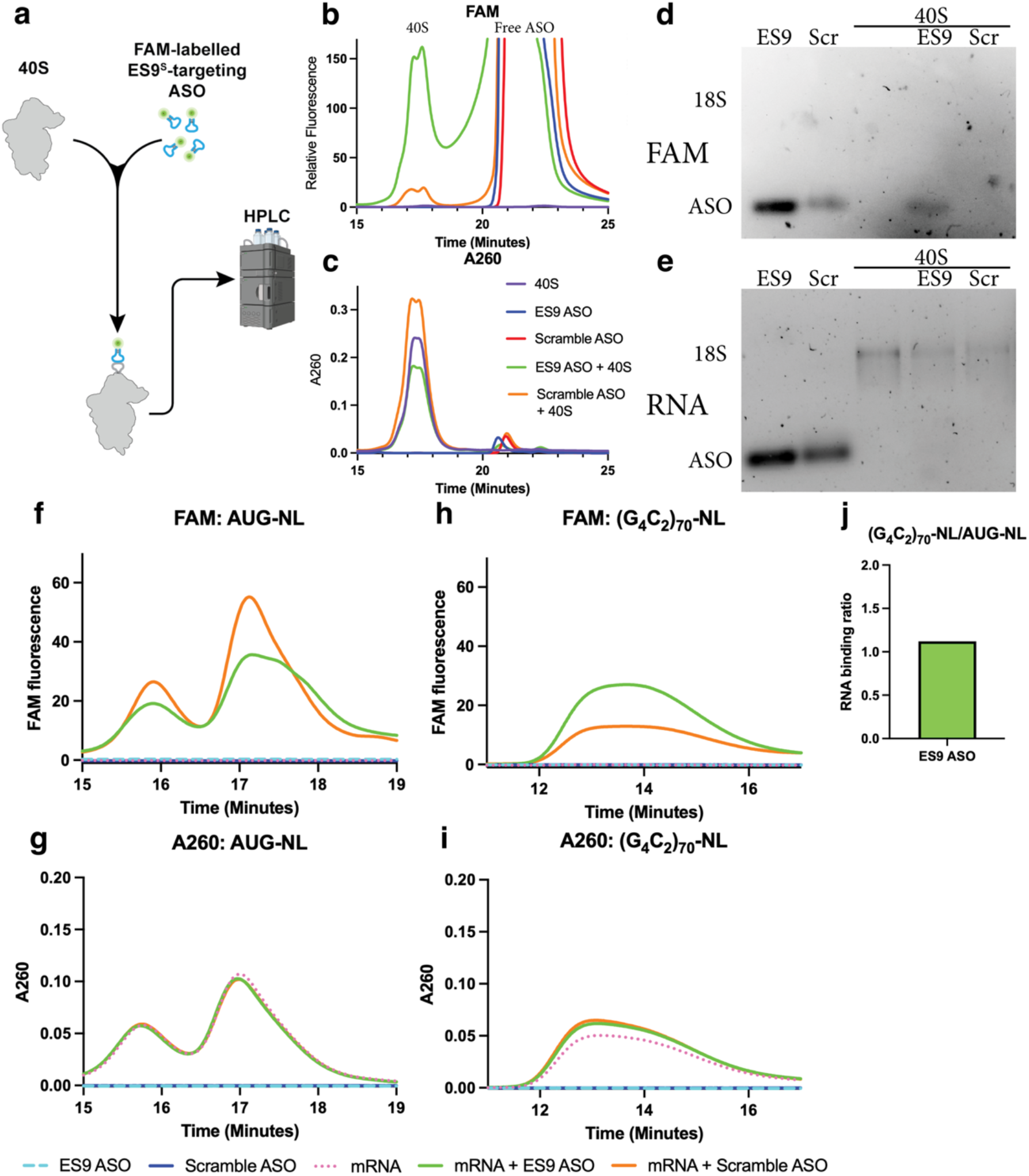
The ES9^S^-targeting ASO strongly and selectively binds the 40S ribosome. **a**, Schematic representation of refolding the FAM-labelled ES9^S^-targeting ASO and mixing with in-vitro purified 40S ribosome then analyzing binding using an HPLC binding assay. **b-c**, HPLC traces of the 40S alone (purple), ES9^S^ alone (blue), Scramble ASO alone (red), ES9^S^ ASO + 40S (green) and Scramble ASO + 40S (orange) shown in the (**b**) the FAM channel identifying the FAM-labelled ASOs and (**c**) the A260 channel depicting RNA. **d-e**, 1% agarose gels showing fractions from the free ES9 and Scramble ASO, then the fractions from the 40S peak for the 40S alone, ES9 ASO + 40S and Scramble ASO + 40S. 18S RNA and ASO bands are highlighted. Gels imaged in (**d**) the FAM channel without dye or (**e**) 1× GelGreen. (**f-i)** mRNA was mixed with either our FAM-labelled ES9^S^-targeting ASO or a scramble and was analyzed for binding efficiency using the HPLC binding assay. RNA either (**f-g**) contained a canonical AUG start codon and nanoluciferase or (**h-i**) the *C9orf72* 5′ UTR-(G_4_C_2_)_70_-nanoluciferase without an AUG start codon. The traces are presented from (**f**,**h**) the FAM channel or (**g**,**i**) the A260 channel. Traces are centered over the respective A260 peaks of the mRNAs. (**j**) Area under the curve (AUC) analyses of the FAM peak(s) corresponding presented as a ratio of AUG-NL/(G_4_C_2_)_70_-NL binding. Data shows that the ES9 ASO showed equivalent binding between the AUG-NL and (G_4_C_2_)_70_-NL mRNAs-suggesting that the ASO is not binding selectively to the repeat mRNA.

**Extended Data Figure 14:**
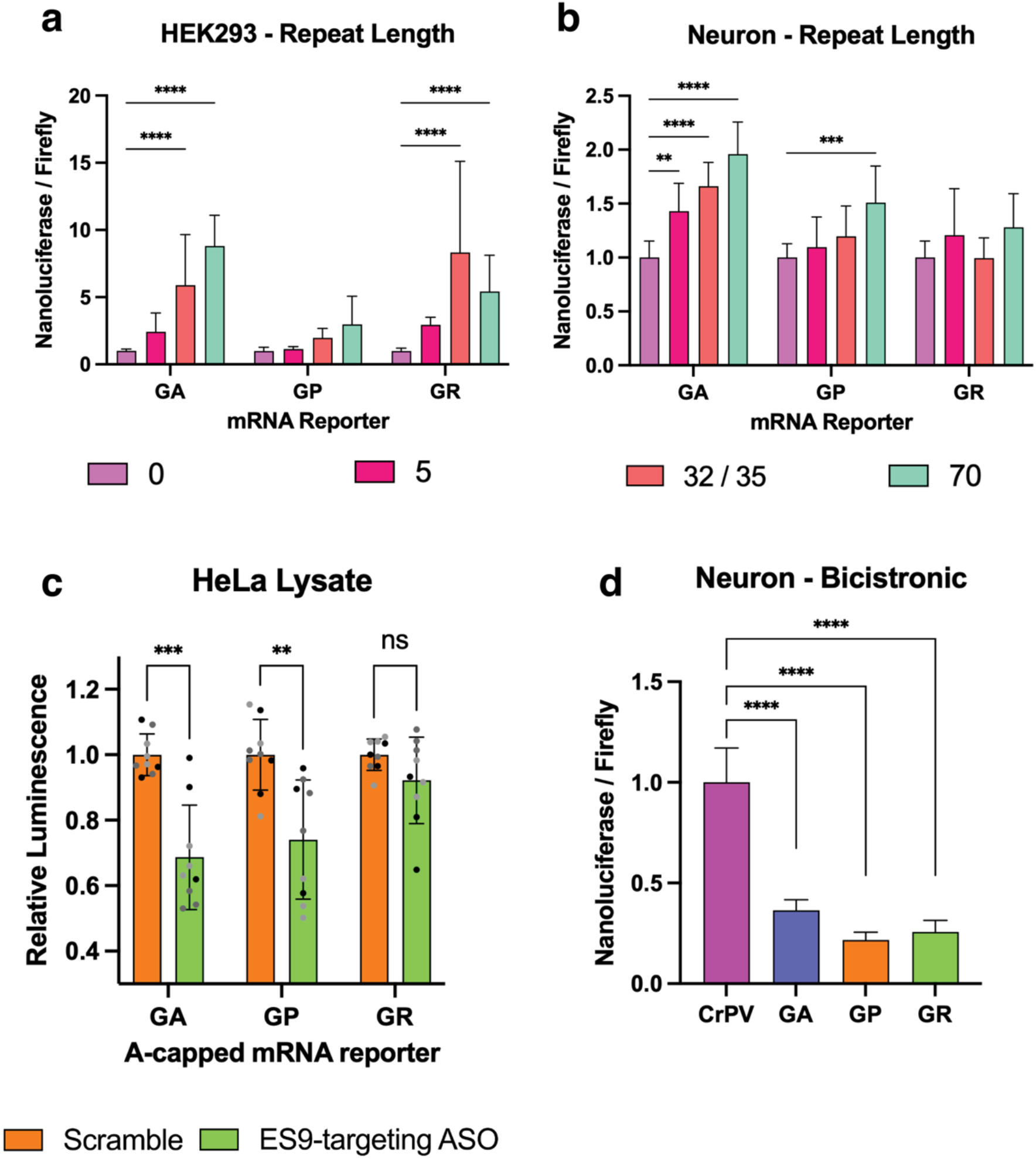
C9orf72 RAN translation can occur through cap-independent mechanisms that are supported by the ES9^S^-G_4_C_2_ kissing loop. **a-b**, RNA of increasing repeat length was transfected in (**a**) HEK293 cells and (**b**) Rat hippocampal neurons in all three reading frames, showing a general repeat length dependence in all conditions tested except for GR in neurons. GP increased in HEK cells but was not statistically significant (neuron n=10, HEK293 n=15). **c**, *In vitro* translation assays using cytoplasmic HeLa cell lysates and nanoluciferase reporters capped with an A-capped analog that does not bind the eIF4F cap binding complex. The A-capped RNAs demonstrate that antisense RNA oligonucleotides at 5 µM concentration reduce C9orf72 RAN translation in the GA and GP frames, but not the GR reading frame when RNAs do not interact with the cap binding complex (n=9). **d**, Bicistronic reporter expression in Rat hippocampal neurons relative to CrPV expression. Error bars for all bar graphs represent the mean ± SD obtained from three independent replicates for all experiments. Statistical significance: **p < 0.01, ***p < 0.001, ****p < 0.0001.

**Extended Data Figure 15:**
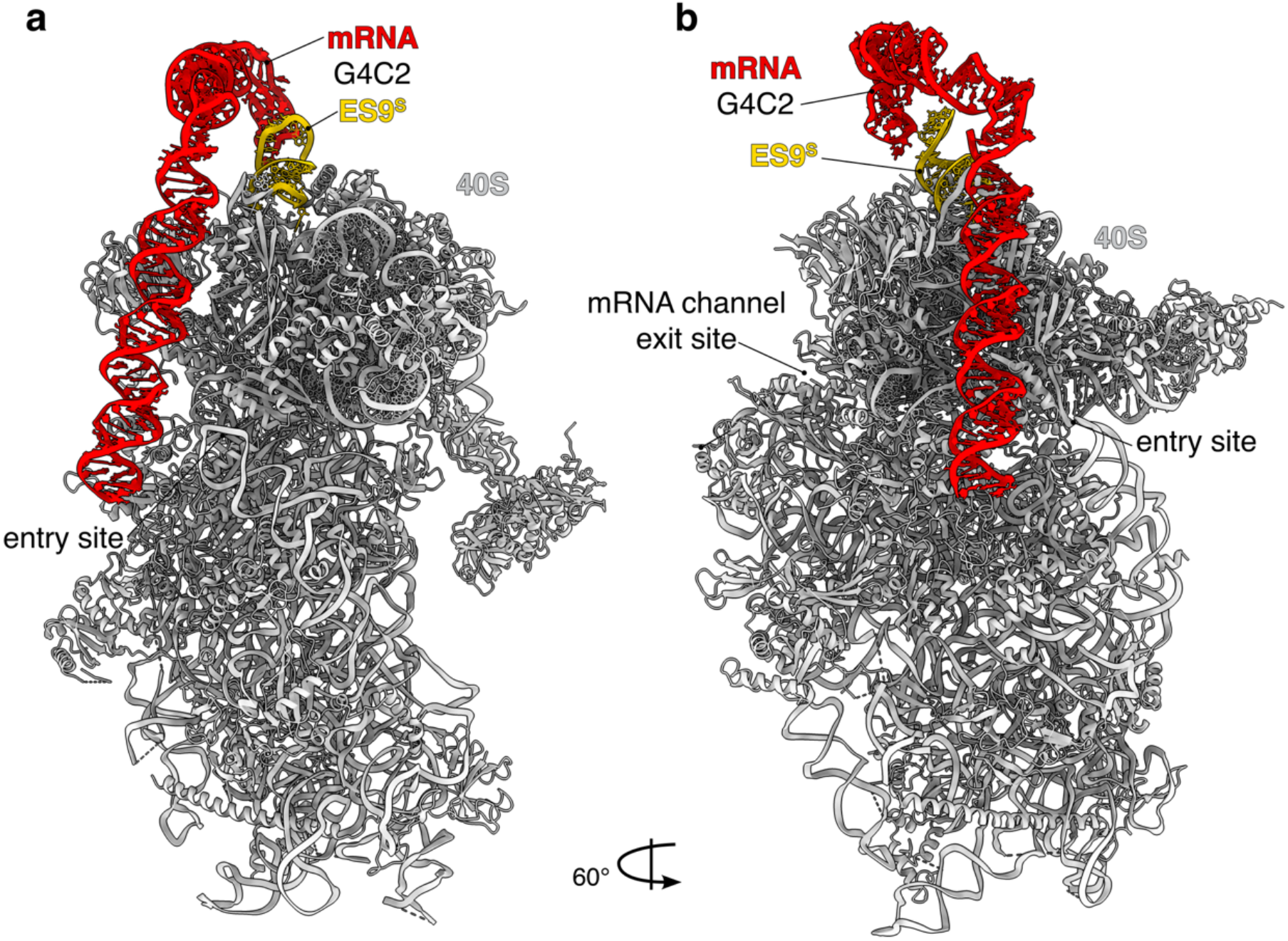
Theoretical model of G_4_C_2_ repeat extension from entry site to ES9^S^. Schematic representation of a helical G_4_C_2_ repeat RNA spanning from the mRNA entry channel to the ES9^S^ region. Based on spatial constraints, a minimum of 28–30 G_4_C_2_ repeats is estimated to be required for this interaction.

**Extended Data Figure 16:**
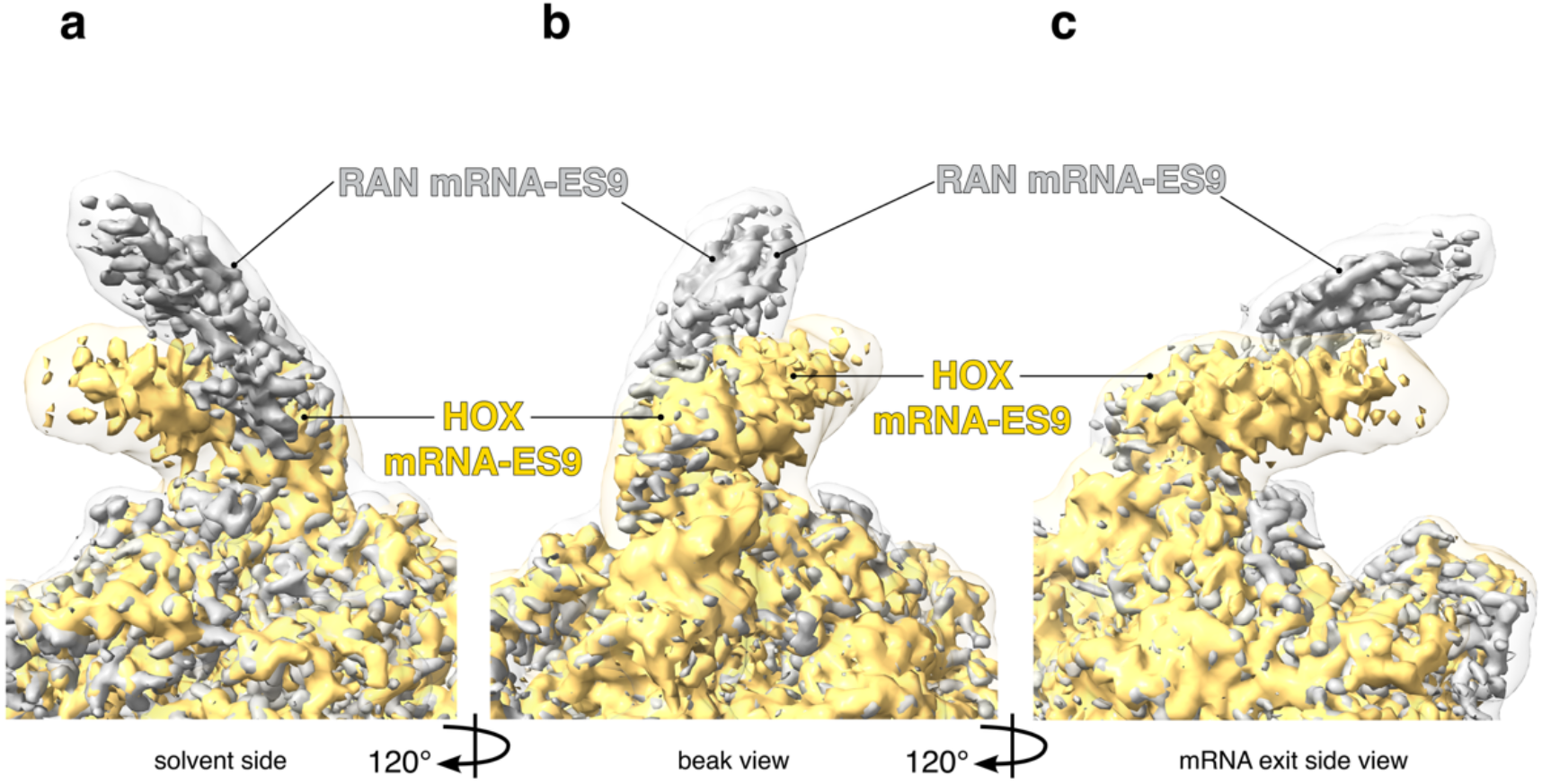
HOX mRNA and RAN mRNA bind at ES9^S^ with different angles of mRNA invasion. **a-c**, Three views identifying the overlap of our *in vitro* reconstituted focus refinement structure (PDB 9PG5) with the mouse *Hoxa9* IRES-like element from Leppek et al.^97^. Comparison of the densities reveals that the two mRNAs overlap at the mRNA-ES9^S^ interaction, with different angles of the invading stem loops approaching the ES9^S^.

**Extended Data Table 1: Sequences of RNA constructs and antisense oligonucleotides**.

**Extended Data Table 2: CryoEM data processing and refinement statistics**.

**Extended Data Table 1.**
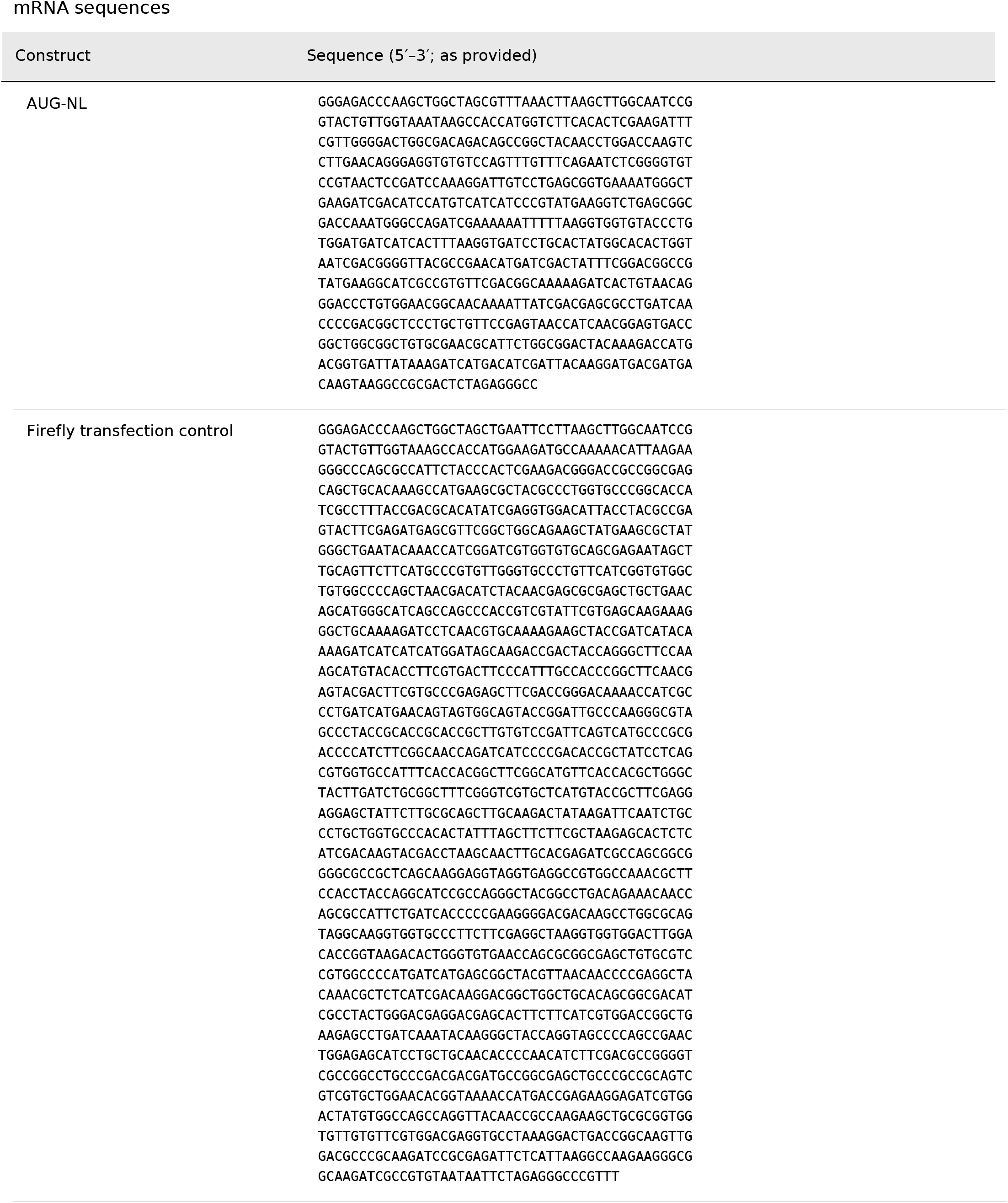
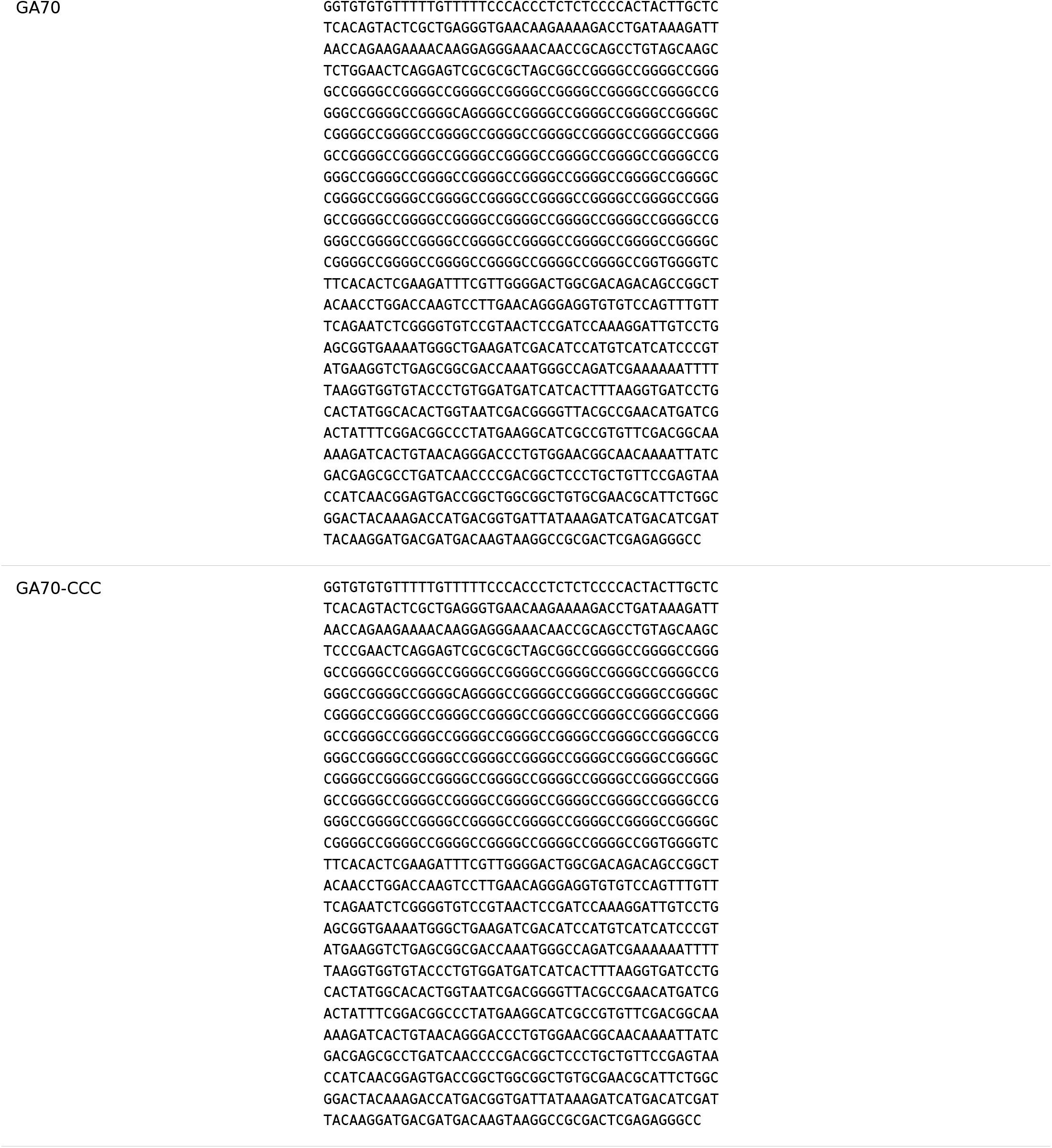
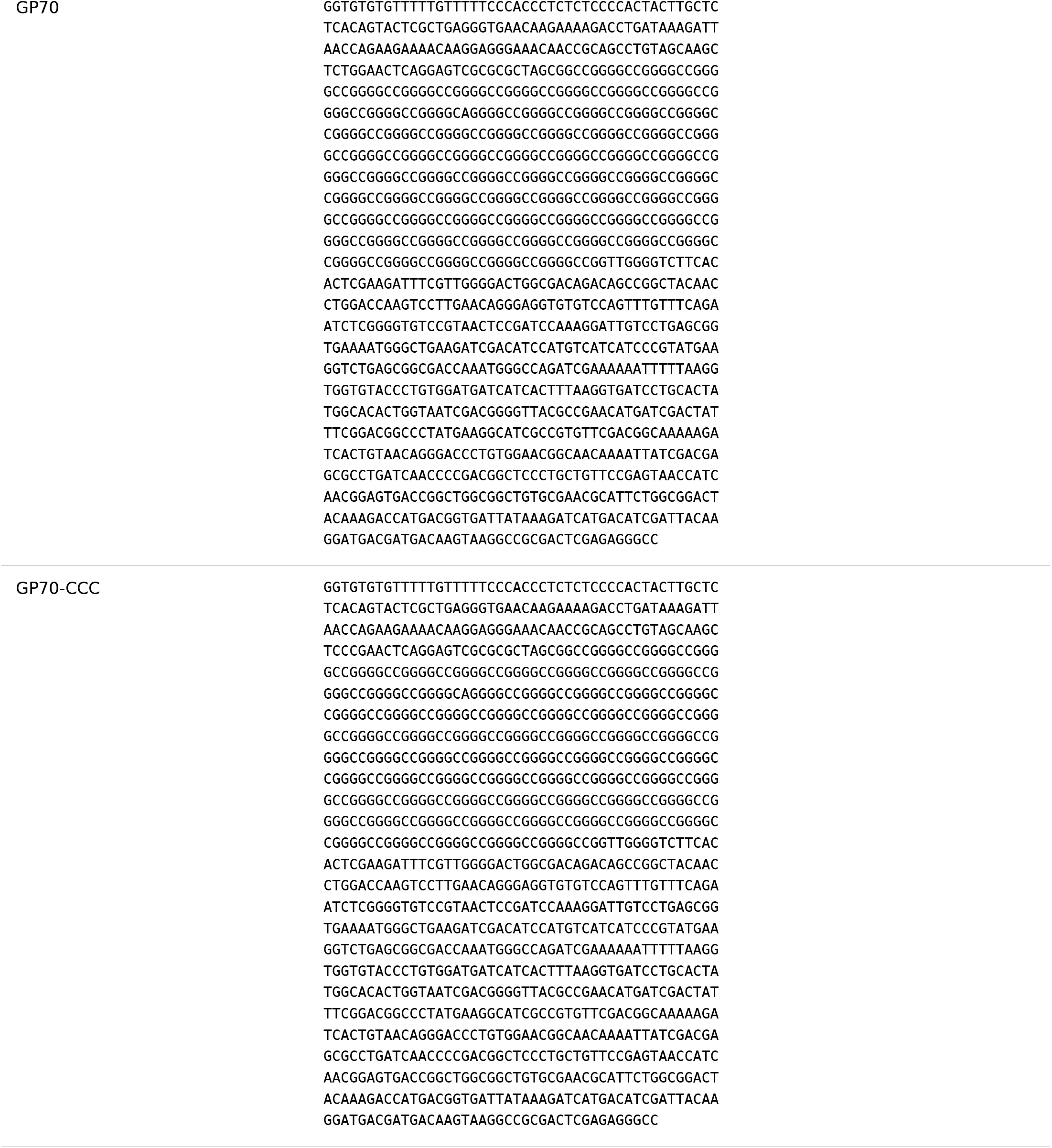
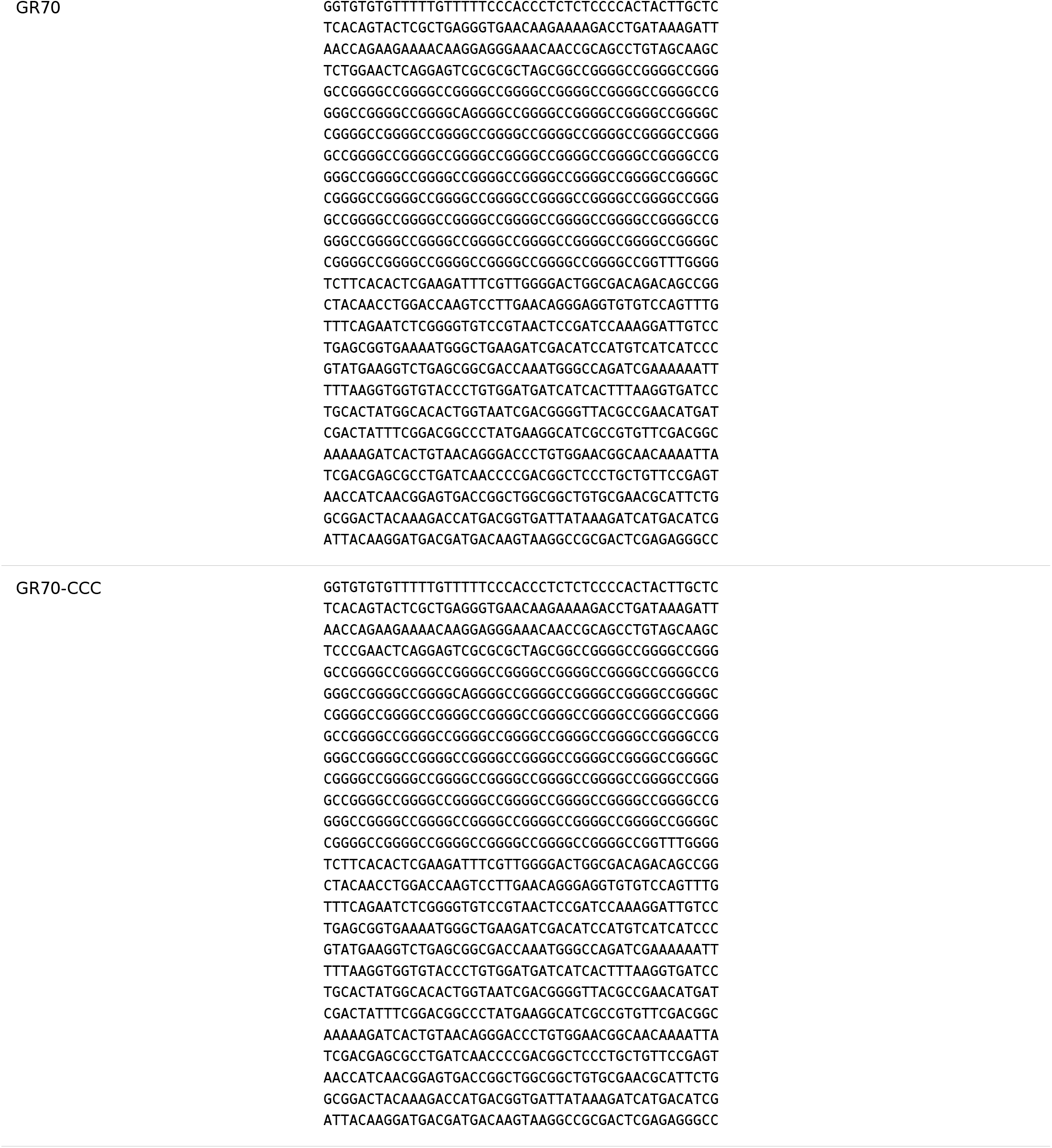
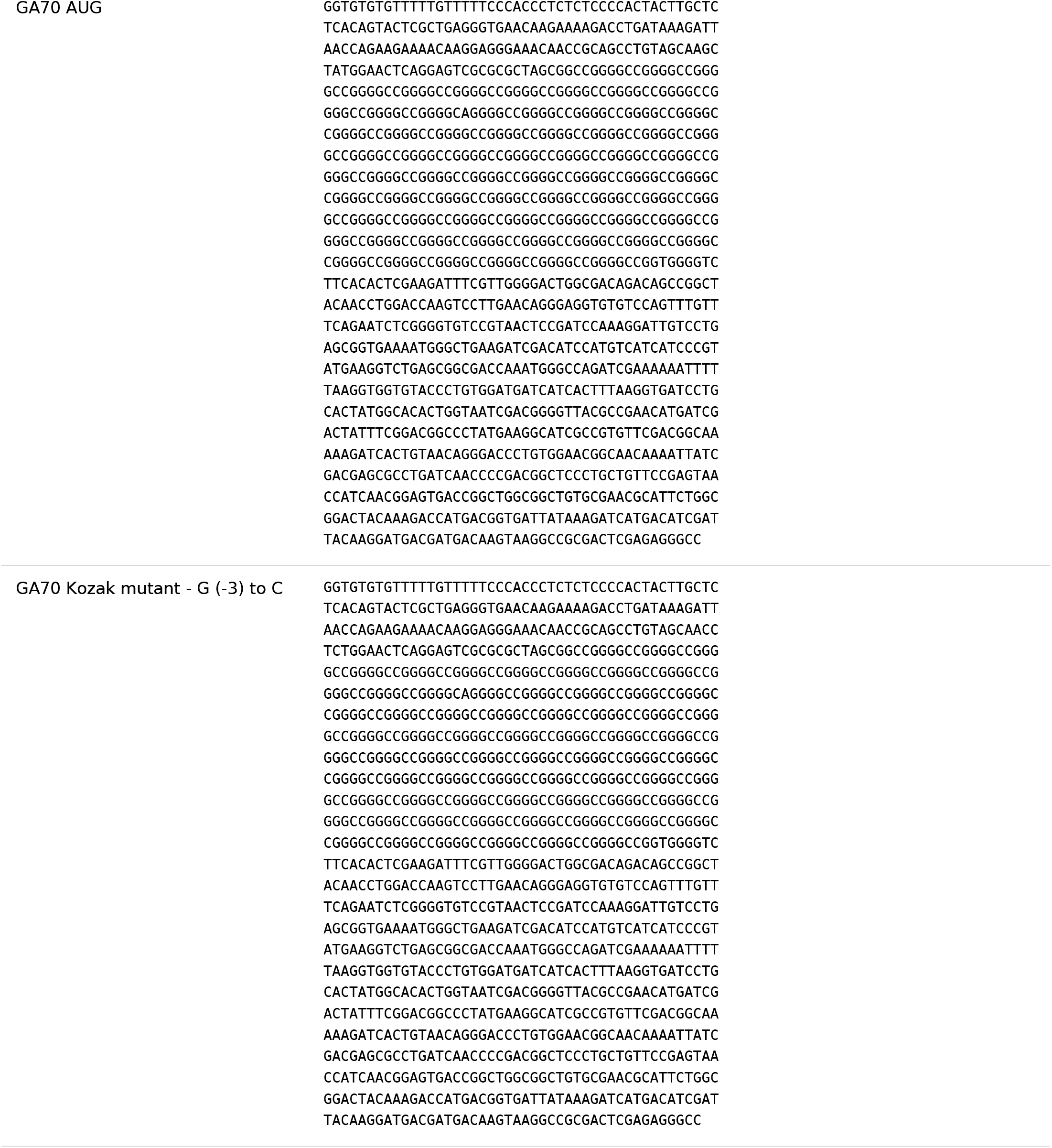
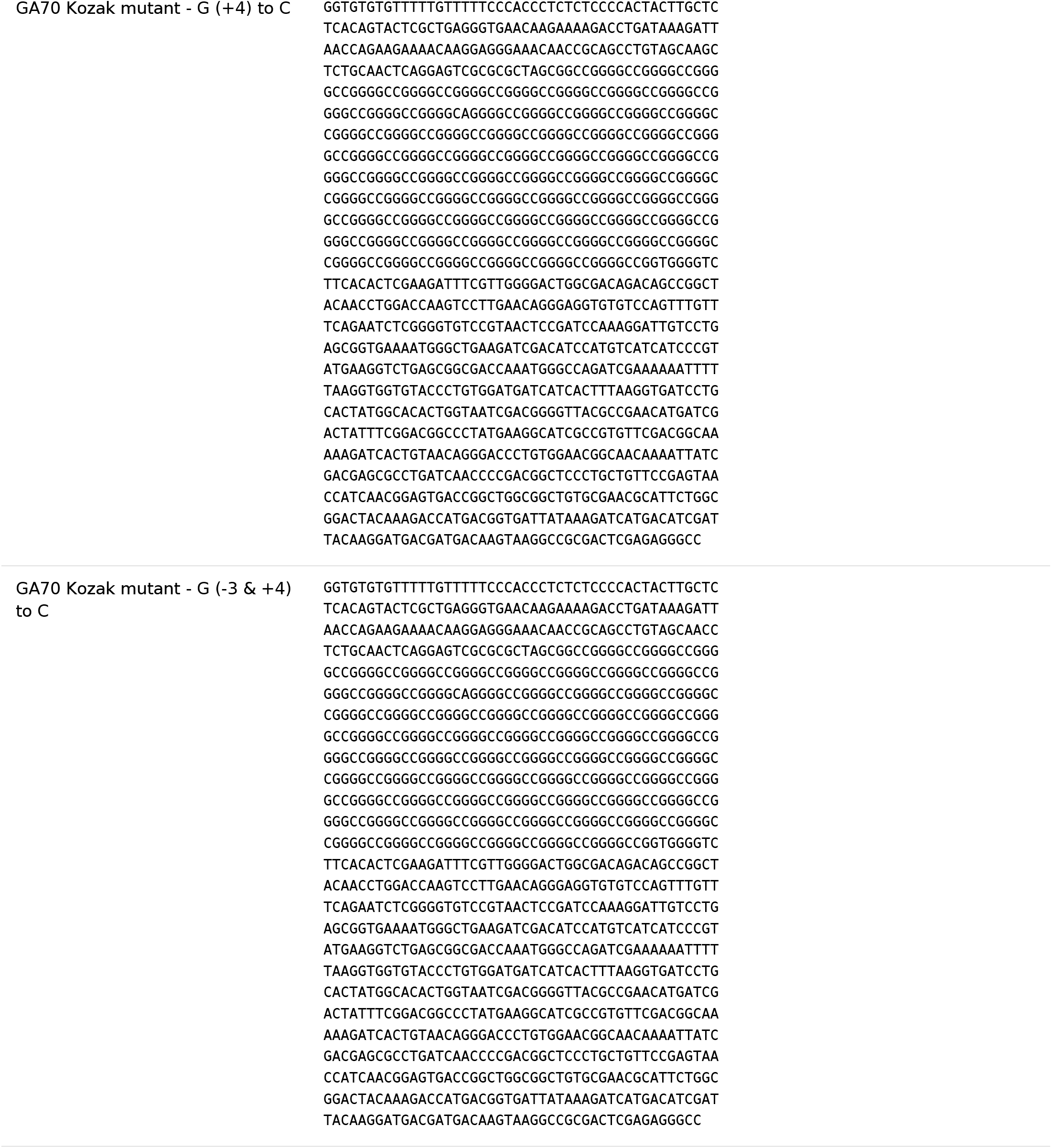
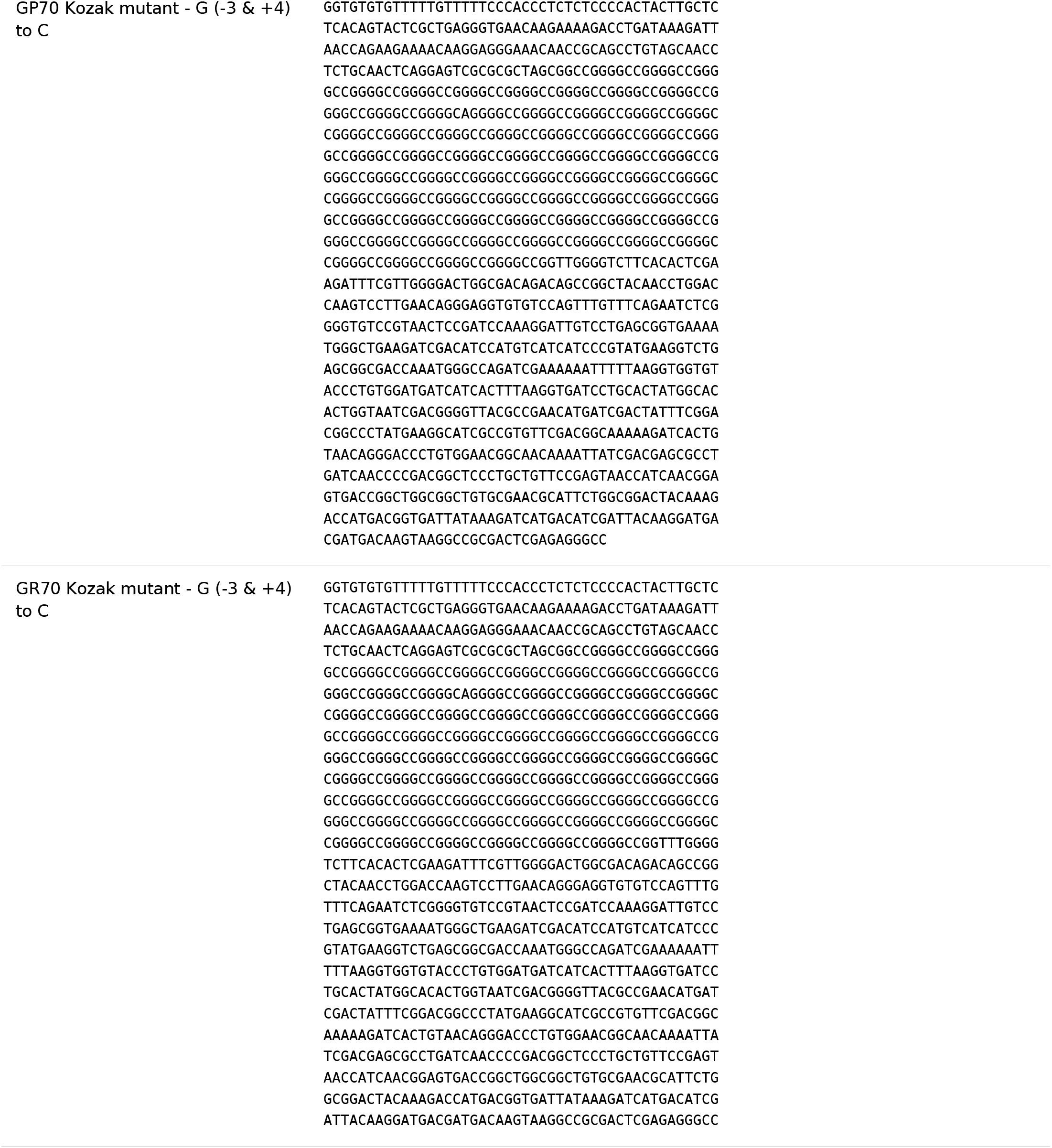
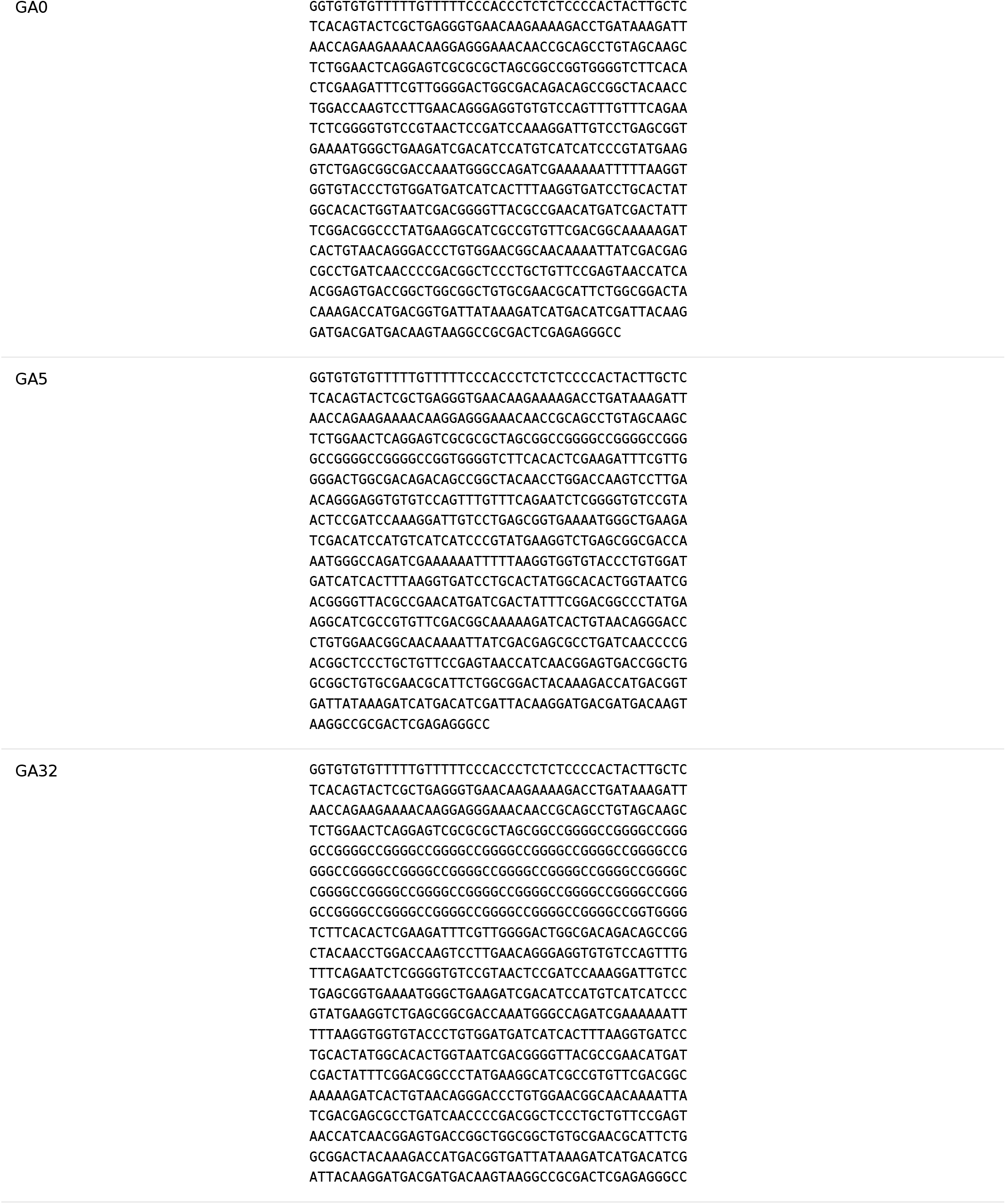
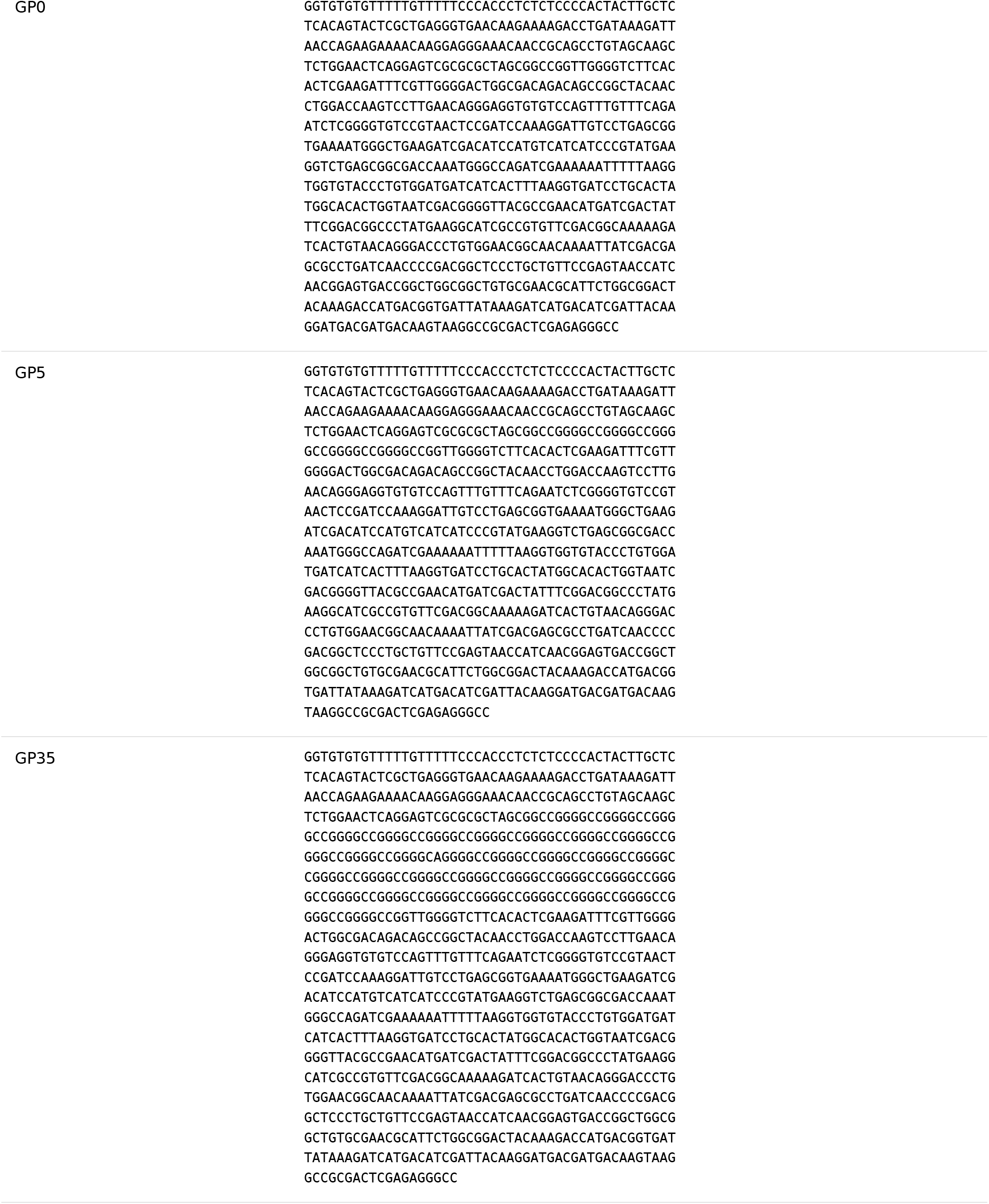
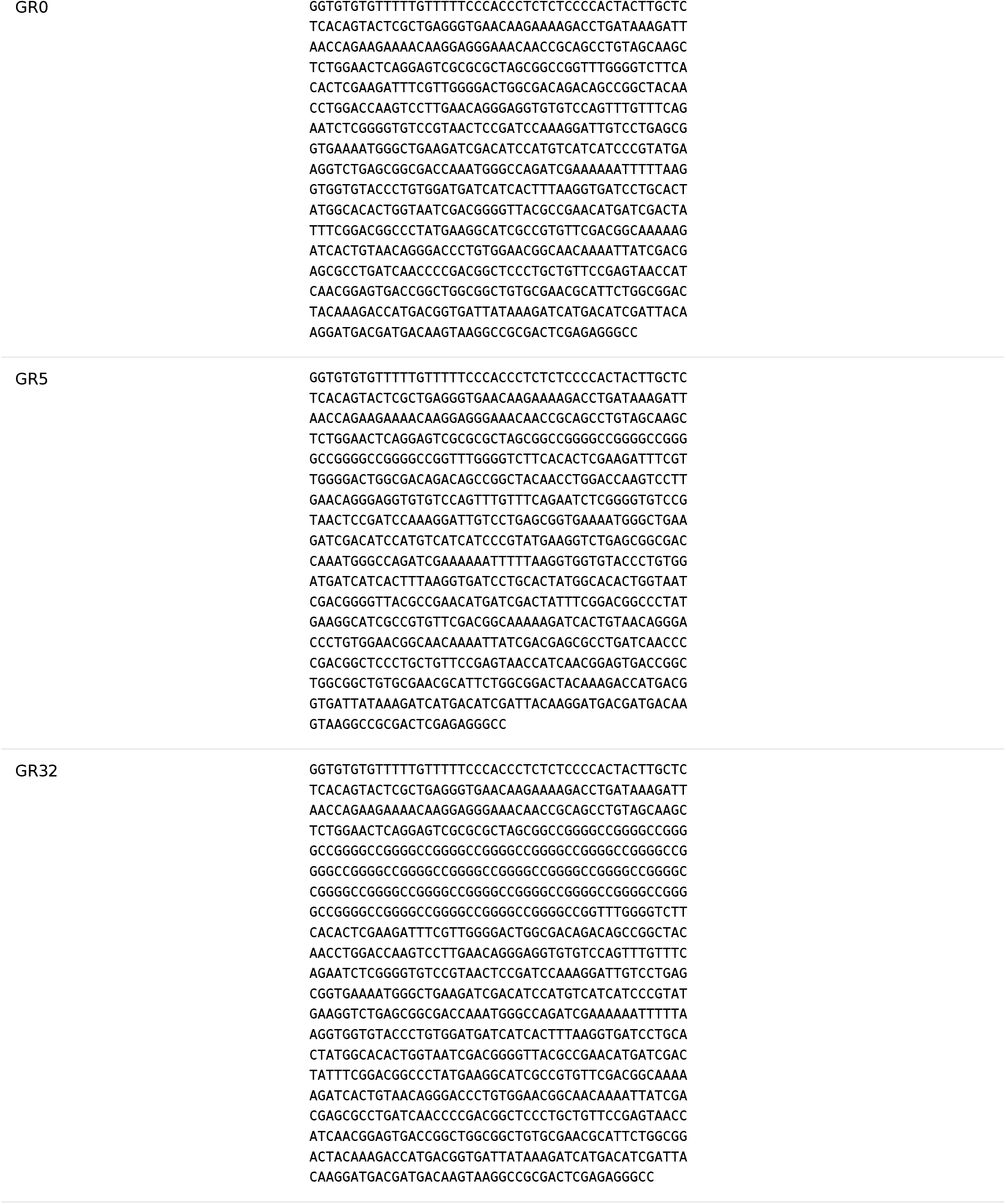
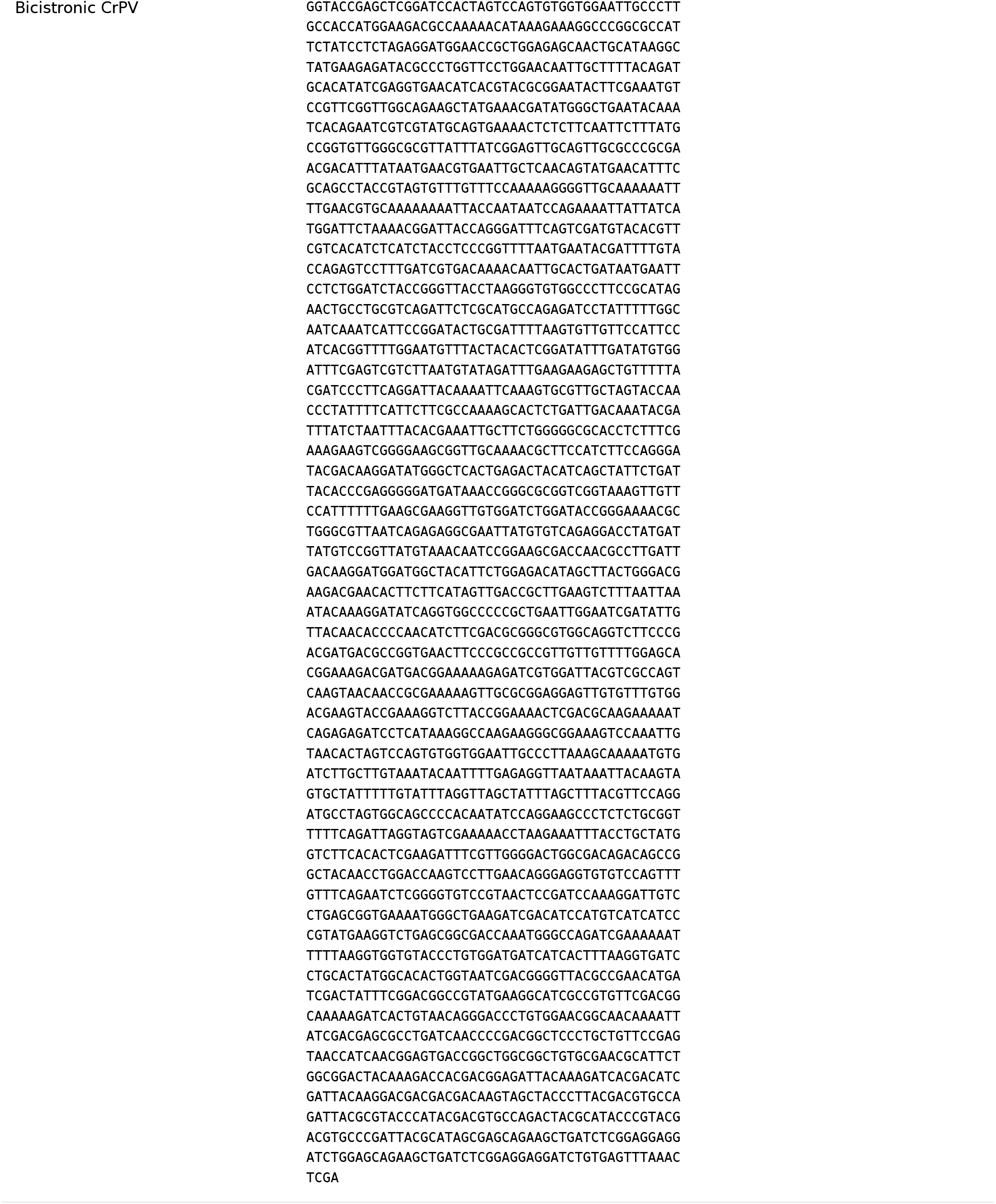
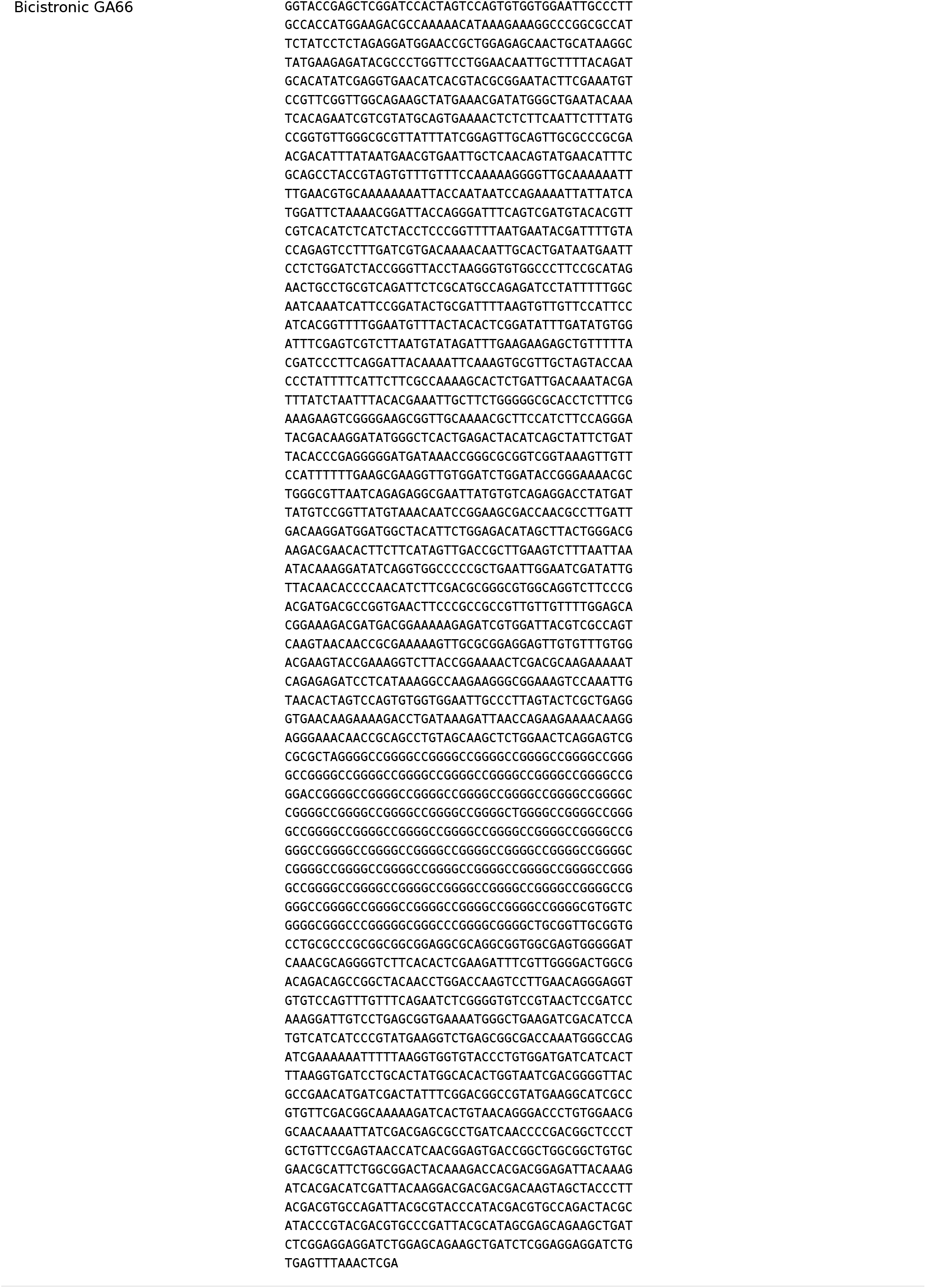
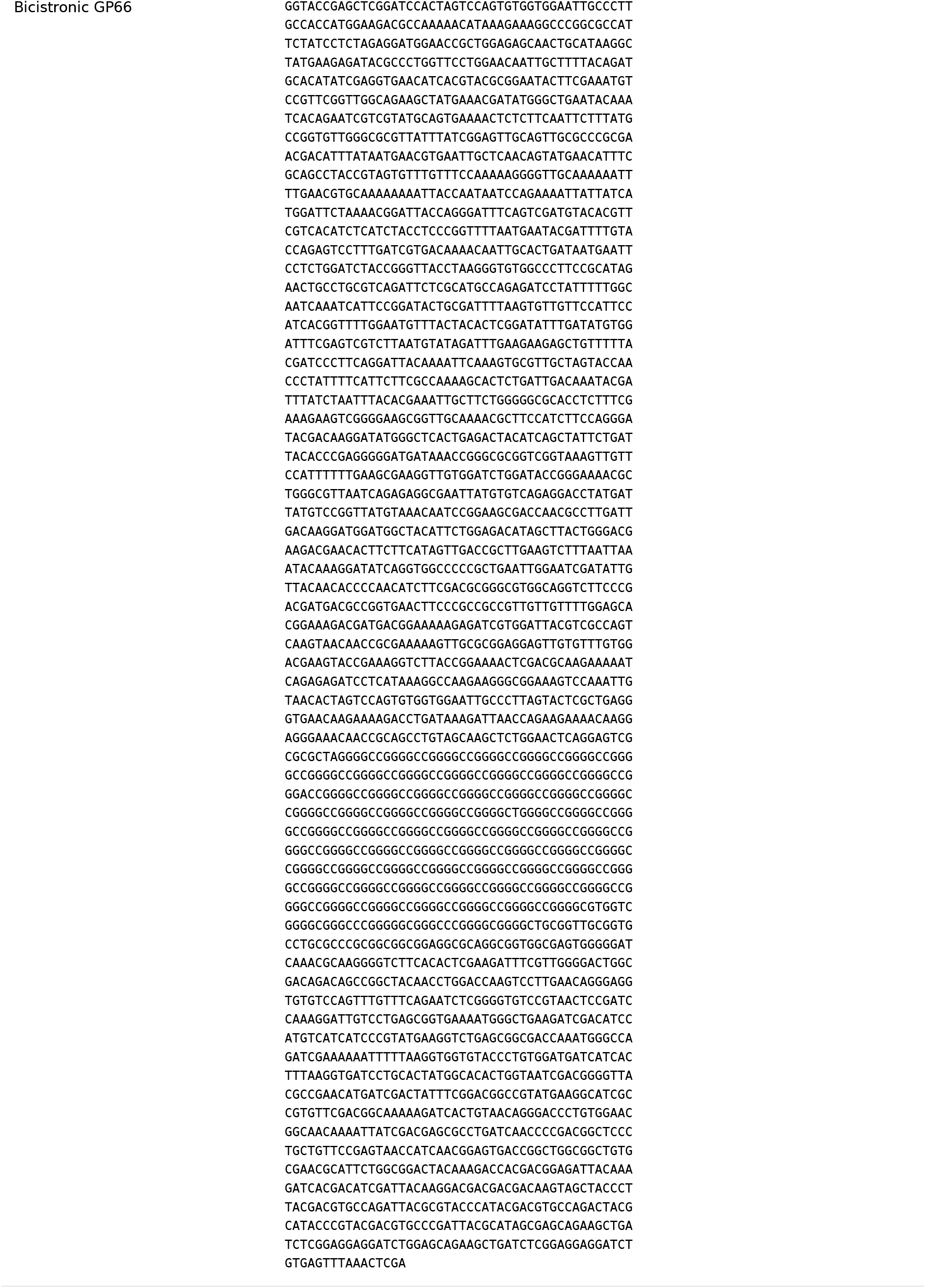
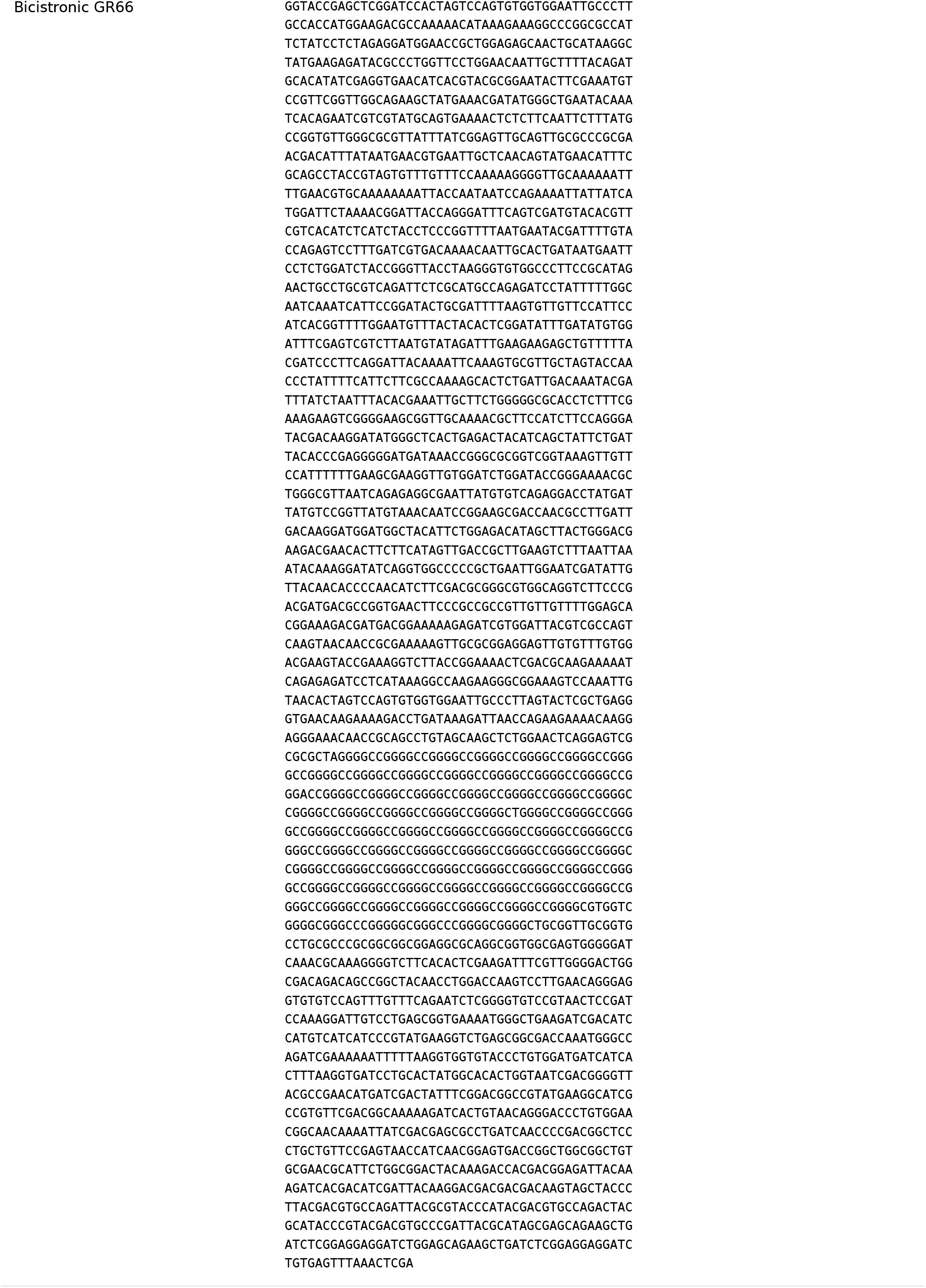
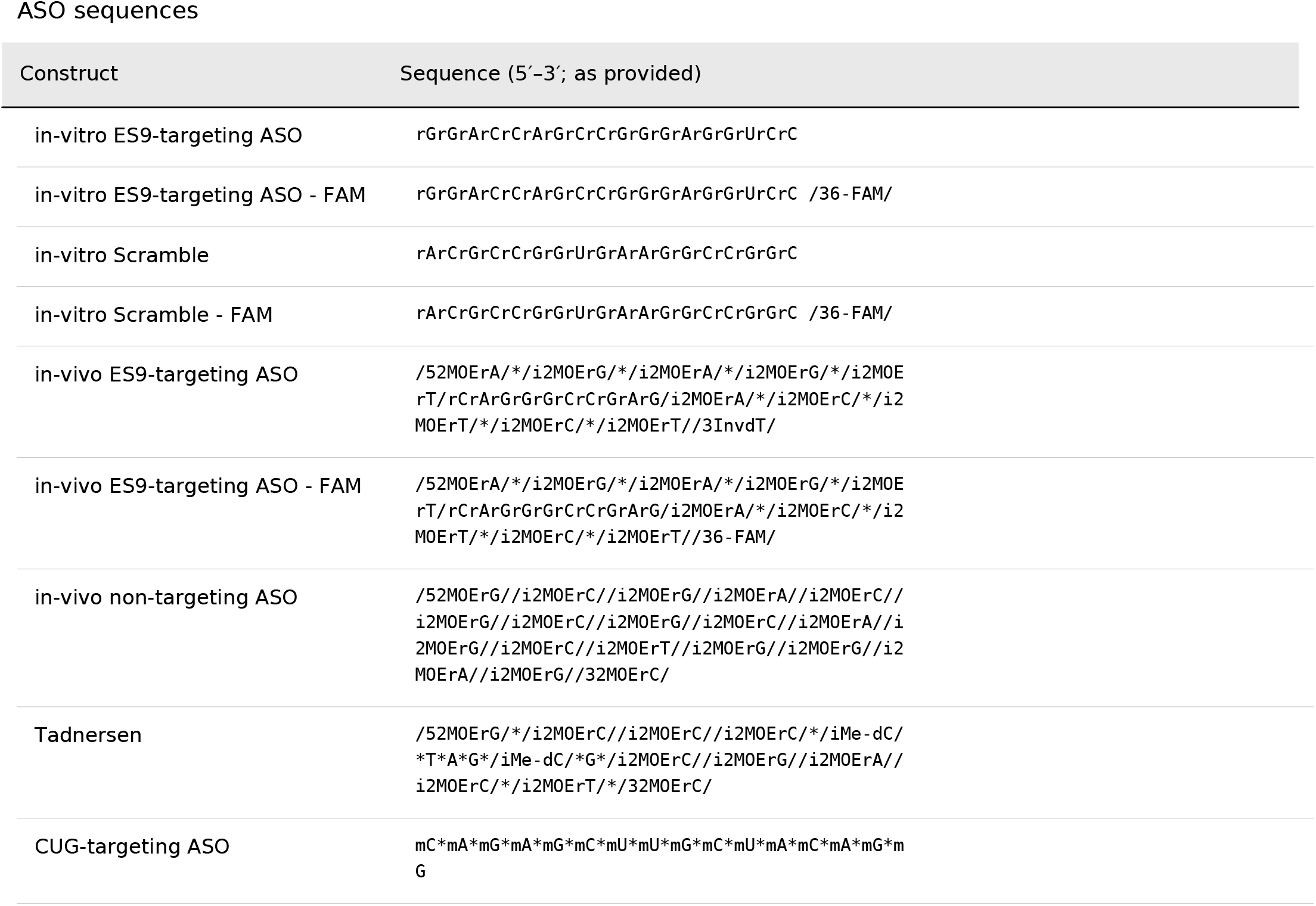
mRNA and ASO sequences.

**Table S2:** Cryo-EM data collection, refinement and validation statistics.

|  | *#in vitro* late-stage<br>initiation complex<br>(EMDB-79722)<br>(PDB: 39RN) | *# in vitro* ES9 focus<br>refinement<br>(EMDB-79720)<br>(PDB: 39RL) | *# near-native* late-stage initiation<br>without eIF3<br>(EMDB-79727)<br>(PDB: 39RT) |
| --- | --- | --- | --- |
| **Data collection and processing** |  |  |  |
| Magnification | 105,000 | 105,000 | 105,000 |
| Voltage (kV) | 300 | 300 | 300 |
| Electron exposure (e<sup>-</sup>/Å<sup>2</sup>) | 1.0461 | 1.0461 | 0.7439 |
| Defocus range (μm) | -1.2 to -3.0 | -1.2 to -3.0 | -1.2 to -3.0 |
| Pixel size (Å) | 0.832 | 0.832 | 0.834 |
| Symmetry imposed | C1 | C1 | C1 |
| Initial particle images (no.) | 2,014,914 | 2,014,914 | 2,177,861 |
| Final particle images (no.) | 14,750 | 45,984 | 18,010 |
| Map resolution (Å) | 3.1 | 2.9 | 3.5 |
| FSC threshold | 0.143 | 0.143 | 0.143 |
| Map resolution range (Å) | 2.5 to 12.5 | 2.5 to 12.5 | 3 to 20 |
| **Refinement** |  |  |  |
| Initial model used (PDB code) | 8pj5 | 8pj5 | 6YAL |
| Model resolution (Å) | 3.0 | 3.0 | 3.4 |
| FSC threshold | 0.5 | 0.5 | 0.5 |
| Model resolution range v(Å) | x to x | 2.5 to 3.0 | 3.0 to 3.4 |
| Map sharpening *B* factor (Å<sup>2</sup>) | -10 | -15 | -35 |
| Model composition |  |  |  |
| Non-hydrogen atoms | 51,166 | 16,363 | 89,481 |
| Protein residues | 3,000 | 1002 | 6,322 |
| Nucleotide | 1,267 | 389 | 1,833 |
| Ligands | 61 | 28 | 5 |
| *B* factors (Å<sup>2</sup>) |  |  |  |
| Protein | 49.87 | 51.51 | 52.57 |
| Nucleotide | 68.41 | 48.66 | 58.29 |
| Ligands | 24.72 | 37.35 | 99.08 |
| R.m.s. deviations |  |  |  |
| Bond lengths (Å) | 0.003 | 0.008 | 0.005 |
| Bond angles (°) | 0.566 | 0.729 | 0.643 |
| Validation |  |  |  |
| MolProbity score | 1.69 | 2.14 | 2.54 |
| Clashscore | 8.63 | 9.34 | 7.28 |
| Rotamer outliers (%) | 0.27 | 3.14 | 4.60 |
| Ramachandran plot |  |  |  |
| Favored (%) | 96.57 | 96.07 | 86.47 |
| Allowed (%) | 3.19 | 3.93 | 13.10 |
| Outliers (%) | 0.24 | 0.00 | 0.43 |

